# Molecular phylogenetics of Neotropical chrysomeline beetles: Evidence for a constrained history of host plant use

**DOI:** 10.64898/2026.01.30.702876

**Authors:** Guillaume J. Dury, Donald M. Windsor, Barbara J. Sharanowski, Lukáš Sekerka, Jacqueline C. Bede

## Abstract

This study reconstructs the phylogeny of an expansive set of Neotropical leaf beetles in the subfamily Chrysomelinae. From 33 species in the genus *Platyphora* Gistel, and an additional 37 species representing 16 beetle genera, five genes, three nuclear, and two mitochondrial, were sequenced and used to obtain a well-supported molecular phylogeny using both Bayesian and Maximum Likelihood. The subtribes Chrysomelina and Doryphorina (sensu Daccordi 1982) were monophyletic, while the genus *Platyphora* was polyphyletic. The genus *Leptinotarsa* Chevrolat is confirmed to be distinct from *Stilodes* Chevrolat. Host plant family was recorded for both adults and larvae using direct observations where possible. Ancestral host plant use was reconstructed using Bayesian trait analyses. A complicated history of host plant switches among a restricted set of plant families is revealed: In the paraphyletic *Platyphora*, one clade that includes *Proseicela* and *Leptinotarsa* had two switches from Asclepiadiodeae to Solanaceae, one switch to Moraceae, and one switch to Malpighiaceae, another *Platyphora* clade had switches between Asteraceae and Rauvolfioideae, and from Rauvolfioideae to Asclepiadiodeae, with other members of the same clade feeding on Boraginaceae and Convolvulaceae. All species included in the clade containing *Tritaenia* and *Stilodes* fed on Malpighiaceae, and all species included in the *Cosmogramma* and *Calligrapha* clade fed on Malvaceae.

## Introduction

The classification of genera in the Chrysomelinae has been problematic, with convergent evolution limiting the use of morphological characters of shape, colour or sculptures for generic delimitation (Daccordi, 1994). For example, the beetles *Platyphora paradoxa* (Achard) and *Euryceraea wagneri* Steinheil are similar in terms of colouration, size, and shape (see photos of *Pl. paradoxa* and another sympatric species *Pl.* sp. nov. “OTG” in the appendix) but differ in other respects such as the shape of the meta- and mesosternum (Daccordi et al., 1999, Steinheil, 1877). Morphological characters such as the intercoxal process have been used to delimit genera by Bechyné (1980) and Daccordi (1994); however, other taxonomic characters such as the chemical composition of defensive secretions (Pasteels, 1993) cast doubt on the classification and suggest some of the characters are homoplastic via convergence. In the subfamily Chrysomelinae, the genus *Platyphora* Gistel contains over 474 species making it, by number of species, the most speciose (Daccordi, 1993, Daccordi, 2017)—though the actual number is likely closer to 467 described species. Similarly to other Chrysomelinae, natural groups are hard to isolate within the genus *Platyphora*, despite attempts by Motschulsky (1860) and Achard (1922, 1923), perhaps because these authors used mostly characters of outer morphological appearance, such as colouring and size and type of elytral punctation (Daccordi, 1993).

Because the phylogenetic relationships of Neotropical Chrysomelinae remain largely unresolved (Hsiao, 1994, Termonia et al., 2002, Gómez-Zurita et al., 2007, 2008, Montelongo and Gómez-Zurita, 2014), here we focus on using molecular data to better understand the phylogenetic history of this group. The most recent taxonomic treatment of the Chrysomelinae divides the subfamily into two tribes: the monotypic Timarchini and the Chrysomelini. The Chrysomelini were divided into twelve subtribes by Daccordi (1982), who merged several in 1994 and proposed to keep three tribes in the Chrysomelini: Entomoscelina, Chrysolinina, and Chrysomelina. We focus especially on the subtribe Doryphorina, which is entirely endemic to the Americas (but introduced elsewhere; Daccordi, 1982). Within Doryphorina, *Platyphora* and several other genera are restricted to the Neotropics (Blackwelder, 1957, Daccordi, 1982, Daccordi, 1993). To date, the largest phylogeny of *Platyphora* included 20 species and was rooted with species from two other genera of Doryphorina (Termonia et al., 2002). The genus *Platyphora* was monophyletic in these studies, although species in some of the most closely related genera were omitted (Pasteels et al., 2003, 2004, Timmermans et al., 1992). Other studies could not verify the monophyly of *Platyphora* because these included only one or two exemplar species per genus (Hsiao, 1994, Gómez-Zurita et al., 2007, 2008). The relation of *Platyphora* to more closely related genera within Doryphorina is poorly known, despite rare and interesting natural history traits. For example, in several species from the genera *Platyphora* and *Doryphora* Illiger, and in most species of *Proseicela* Chevrolat, females care for larvae until pupation (Windsor et al., 2013).

Chrysomelinae are overwhelmingly host-specific at the species level (Jolivet and Hawkeswood, 1995, Strauss, 1988). Within genera of broad-shouldered leaf beetles (Chrysomelinae), host-plant families are often associated with groups of beetle species, which could be useful for the recognition of natural genera or sub-genera (Pasteels et al., 2004). Moreover, a phylogenetic framework would help clarify the evolution of plant host-specificity, which involves the coevolution of maternal oviposition preferences and larval nutritional development. To build a more encompassing phylogenetic framework, we collected specimens from 63 species and 12 genera from Central and South America, and 7 species and 4 genera from North America, Asia, Europe, and Australia. These include representatives of two subtribes Chrysolinina and Chrysomelina (sensu Daccordi, 1994) and representatives from four of twelve subtribes (sensu Daccordi, 1982): Phyllodectina, Doryphorina, Chrysomelina, and Chrysolinina. Determinations of host plant usage and host plant identifications for all but the three Brazilian species were made directly by the authors (GJD, DMW). Our sampling, the largest to date, includes 36.6% (15) of the 41 genera of Chrysomelinae in the Neotropical region according to Daccordi (1994), and half (11) of the 22 genera in the subtribe Doryphorina (Table 1, photographs in supplementary materials). Here, we infer the evolutionary relationships of Chrysomelinae using five genes, three mitochondrial and two nuclear, and use the molecular phylogeny to infer ancestral host plant use in Neotropical members of the subfamily.

**Table 1.**
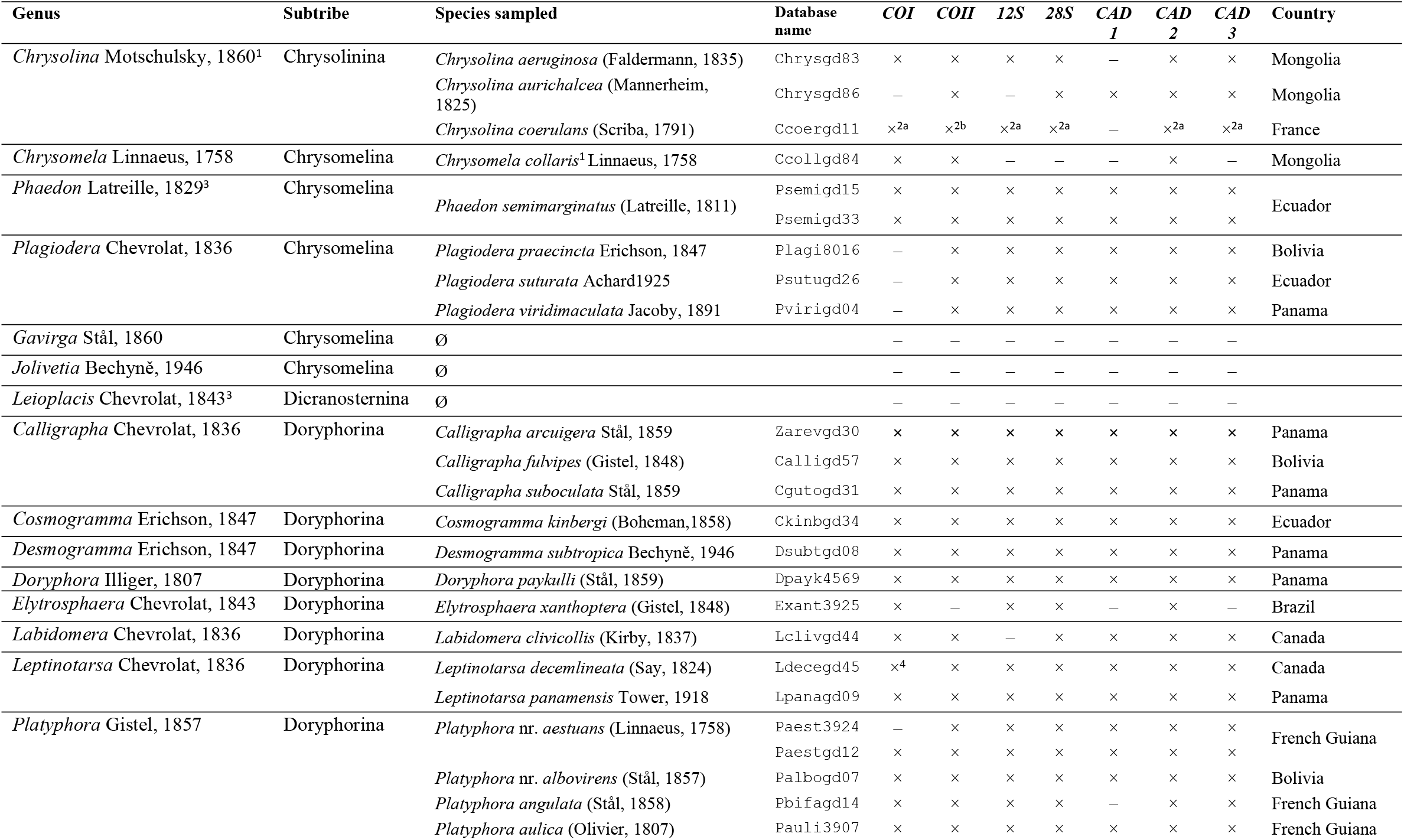
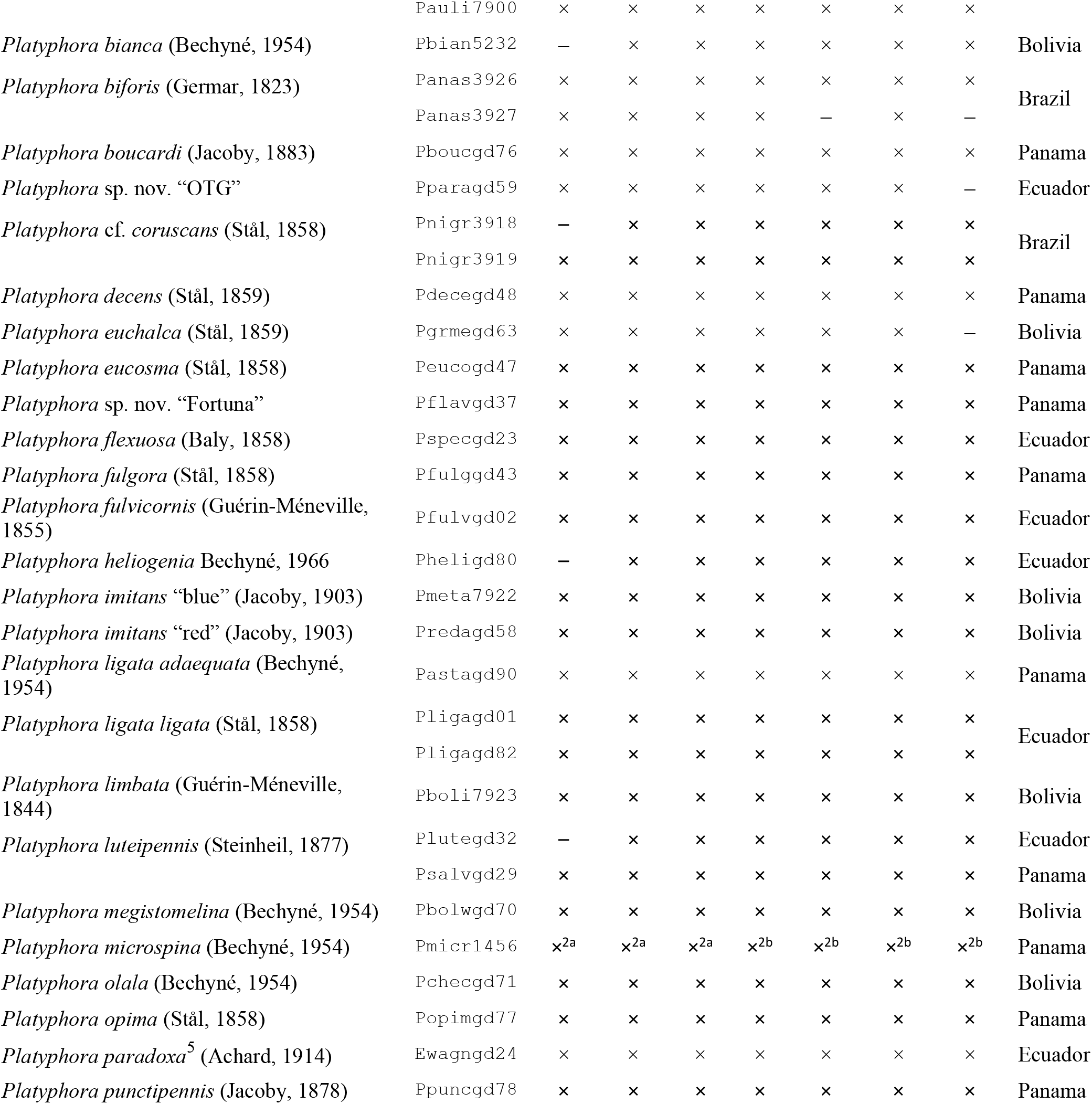
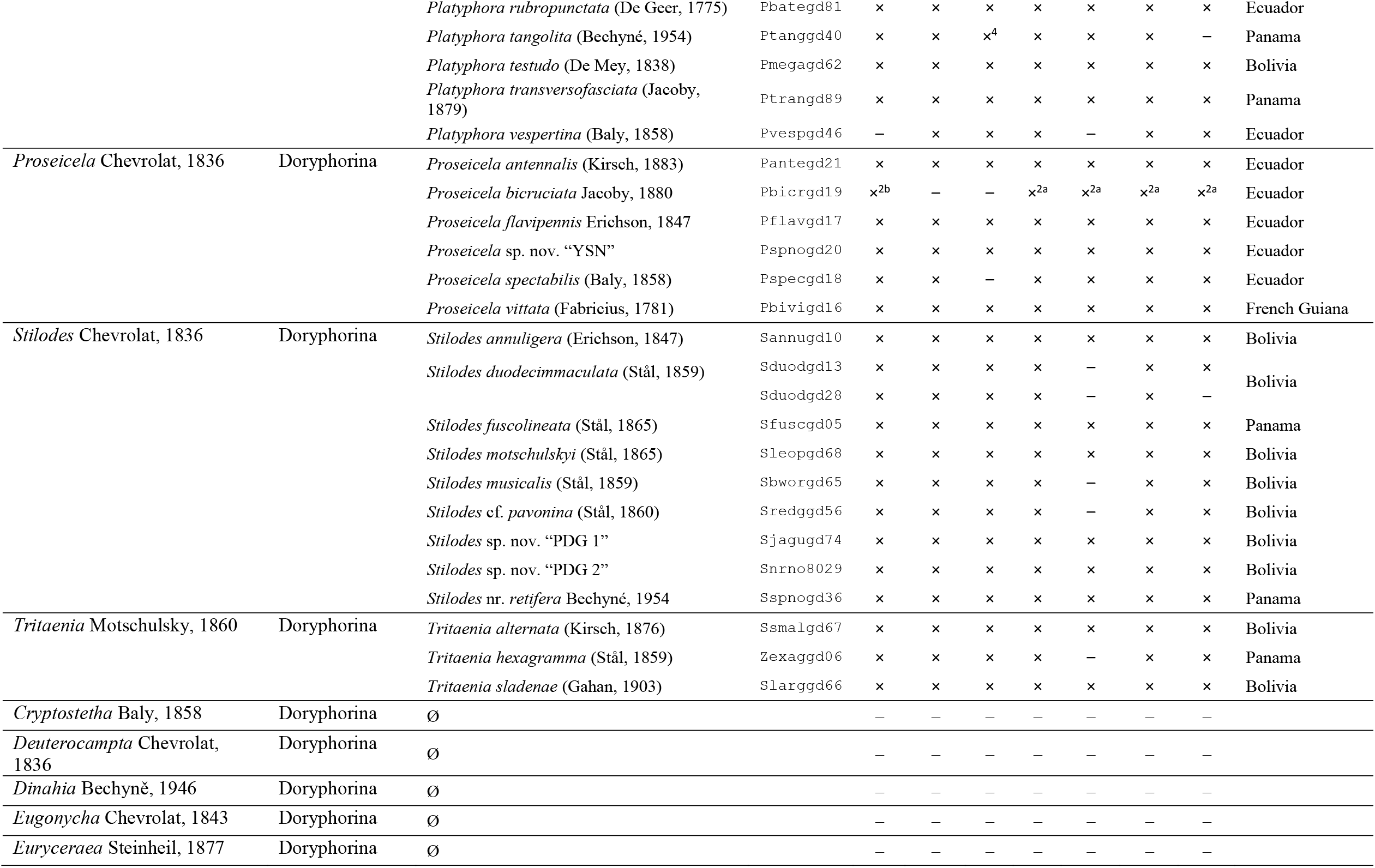
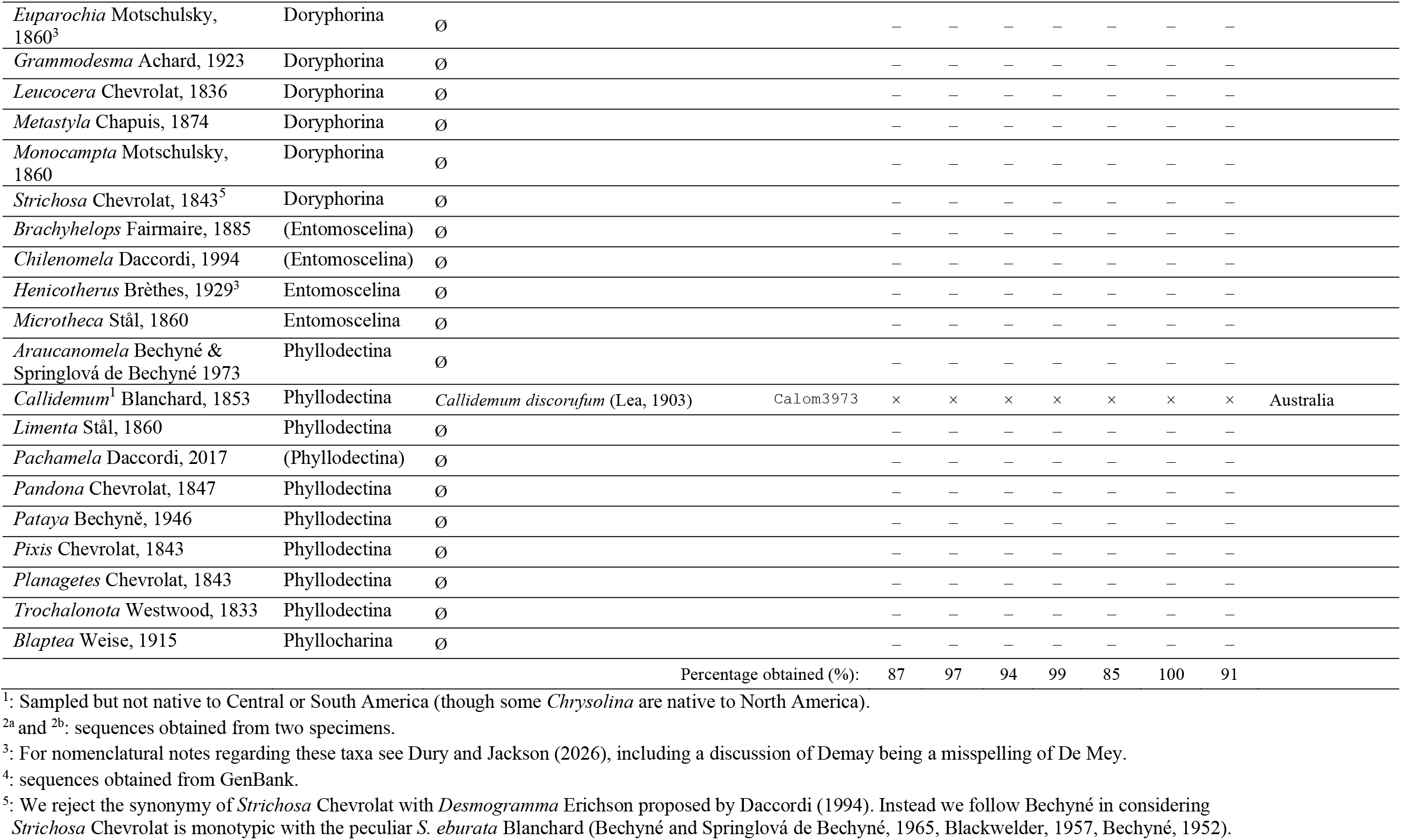
Genera of tribe Chrysomelini native to South or Central America according to Daccordi (1994; 2017; incorporating recent changes), subtribes sensu Daccordi (1982), species sampled, location of collection, and genes that were amplified for each taxon.

## Materials andMethods

### Taxonomic Sampling

Beetle specimens were collected in the Neotropics (Panama, Ecuador, Bolivia, French Guiana, and Brazil), Nearctic (Canada), Palaearctic (France and Mongolia), and Australasian regions (Australia; Table 1). The majority of specimens were collected through inspection of host plants, known through published records or personal experience. Observation/collection sites were documented through Global Positioning System (GPS) co-ordinates or, in some cases, the use of satellite photos. Live insects and host plants were photographed. Adult beetles found without larvae were kept in the laboratory to confirm host plant use through feeding, and rearing of offspring, on the putative host plant. Larvae found in the field were also reared to maturity on foliage from the plant on which they were found, enabling identification from adult characteristics. For all samples except those from Ecuador, we deposited specimen vouchers of insects and samples of host plants in the Smithsonian Tropical Research Institute, Tupper Research and Conference Center, Panama City, Panama. Beetle specimens were preserved in absolute ethanol and stored at −20°C. Specimen vouchers of beetles from Ecuador were pinned and deposited in the collection of the Quito Catholic Zoology Museum at the Pontificia Universidad Católica del Ecuador.

### DNA extraction and sequencing

Total genomic DNA was extracted from adult thoracic muscle tissue or from legs using the DNeasy Blood & Tissue kit (Qiagen Inc., Valencia, CA, U.S.A.), following insect tissue protocol (Qiagen, 2006). Polymerase chain reaction (PCR) was used to amplify two nuclear genes, *28S* rDNA and *CAD* (carbamoyl-phosphate synthetase 2, aspartate transcarbamylase and dihydroorotase; see Figure S1 for segments used and overlap), and three mitochondrial genes, *12S* rDNA, cytochrome c oxidase subunit I (*COI*), and cytochrome c oxidase subunit II (*COII*) (3709 bp in total; Table S1). PCR amplifications were performed using an Eppendorf epgradient S or Mastercycler thermal cycler (Eppendorf, Hamburg, Germany). Reactions were done using 0.03 Unit/µL of *Taq* DNA polymerase and 1X standard *Taq* buffer (New England Biolabs, Ipswich, MA, U.S.A.), with 0.4–1.2 ng/µL of sample DNA and the following modifications: of Taq DNA polymerase, 80 µM dNTPs (New England Biolabs), 0.4 µM of each primer and water to create a final volume of 25 µL. Conditions for the amplification each gene are listed in Table S2. Volume was doubled for low yielding amplifications and were decreased by two thirds for the first amplification of nested PCR (Table S2).

PCR products were separated either on a 1% agarose gel or a 2% agarose gel for *12S* mtDNA amplicons and stained with either ethidium bromide (Sigma-Aldrich Corp., St. Louis, MO, U.S.A) or SYBR Safe (Invitrogen, Carlsbad, CA, U.S.A.). Desired bands were excised and purified using the QIAquick gel extraction kit (Qiagen Inc.) following the manufacturer’s instructions except that the elution buffer was heated to 70°C to increase the yield at the final step. PCR products were sequenced in both directions by the Institut de Recherche Clinique de Montréal using a Genetic Analyzer 3130xl (Applied Biosystems, Foster City, CA, U.S.A.). Forward and reverse sequencing chromatograms were compiled into contigs, reconciled and edited in Geneious version 4.8.5 (Biomatters Ltd., 2009). Sequences were deposited in GenBank under accession numbers [submission to GenBank to be done].

The majority of sequences obtained in this study were amplified using DNA from a single insect specimen. However, when amplification and sequencing failed, the DNA from a second specimen from the same population was used. This occurred for four samples: *28S* and *CAD* for *Platyphora microspina* (Bechyné), *COI* for *Proseicela bicruciata* Jacoby, and *COII* for *Chrysolina coerulans* (Scriba). Two sequences were obtained from Genbank: *12S* for *Platyphora tangolita* (Bechyné) (accession number AY055560) and *COI* for *Leptinotarsa decemlineata* (DQ649098). Outgroup sequences, basal to the entire ingroup (Gómez-Zurita et al., 2008), were chosen among the closest taxa available in Genbank: *COI* and *28S* from *Timarcha tenebricosa* (Fabricius) (Chrysomelidae: Chrysomelinae: Timarchini; AY171412 and AY171439); *COII* from *Timarcha geniculata* Germar (AJ236336); and *12S* from *Prosopodonta limbata* Baly (Chrysomelidae: Hispinae; AF097125). The outgroup sequence for *CAD* was a combination of 1030 bp from *Prosopodonta limbata* (this study) and 606 bp from *Strangalia bicolor* (Swederus) (Cerambycidae: Lepturinae; GQ265599).

### Sequence alignment and phylogenetic inference

Protein coding genes *COI, COII,* and *CAD* were aligned in Geneious by hand using translated segments. As there were no insertions or deletions in these gene regions the alignment was straightforward. Sequences from the ribosomal gene *28S* were aligned in BioEdit version 7.1.7 (Hall, 1999) using the secondary structure model for Galerucinae as outlined by Gillespie *et al.* (2004). Regions of ambiguous alignment (RAAs) and regions of expansion and contraction (RECs) were excluded from the analysis (∼14% of each *28S* sequence). Ribosomal gene *12S* was aligned using 10 iterations of MUSCLE (Edgar, 2004) in Geneious with default settings. Bayesian phylogenies were inferred with MrBayes v.3.2.1 (Ronquist et al., 2012) on the Compute Canada WestGrid high-performance computing facilities (Western Canada Research Grid). Bayesian analyses of single gene datasets (Figs S3 to S7 in the supplementary material) were completed with 1 million generations and the five-gene concatenated dataset was analysed with 10 million generations (Fig. 1). Convergence of runs and a suitable burn-in were determined using Tracer v.1.5 (Rambaut and Drummond, 2009): 5% for the concatenated dataset analysis and 10% for single-gene analyses. Maximum Likelihood (ML) phylogenies with 100 bootstrap iterations were inferred using RAxML-HPC (Stamatakis, 2006) on supercomputers of the Cyberinfrastructure for Phylogenetic Research (CIPRES) Science Gateway v.3.3 (Miller et al., 2010). Bayes factor tests were conducted in MrBayes to test support for constrained monophyly of genus *Platyphora* and to test for constrained monophyly of genera *Stilodes* and *Leptinotarsa*. For these, stepping stone analyses were conducted with 5 million generations, sampling every thousandth generation and using a burn in of 25%. For single gene analyses, the models of nucleotide substitution were determined with MrModeltest v.2.3 (Nylander, 2008) in Paup* v.4.0b10 (Swofford, 2003) using the PaupUP graphical interface (Calendini and Martin, 2005). For the dataset containing all concatenated sequences, the ideal partitioning strategy and models of nucleotide substitution were determined using PartitionFinder v.1.0.1 (Lanfear et al., 2012). This partition scheme was used for both the Bayesian and ML analyses. Twelve character-sets were pre-defined as follows: all codon positions of all coding genes (*CAD, COI* and *COII*; 9 character-sets total) and the stems of *28S*; non-pairing loops and arcs of *28S*, and the complete *12S* gene region. The ideal partitioning scheme determined using PartitionFinder divided the dataset into four partitions that included the following gene segments: Partition (1): first and second codon positions of *CAD, COI*, and *COII* and all regions of *28S* (except the excluded RAAs and RECs); Partition (2): third codon positions of *CAD*; Partition (3): third codon positions of *COI* and *COII*; Partition (4): *12S*. Data files (including alignments in nexus format) are available at the Texas Data Repository (https://dataverse.tdl.org) under DOI: [submission to Texas Data Repository will be done upon acceptance]. A comparison of gene performance was done by summing the observed posterior probabilities across the nodes and dividing by the maximum possible posterior probability (Wild and Maddison, 2008).

**Figure 1.**
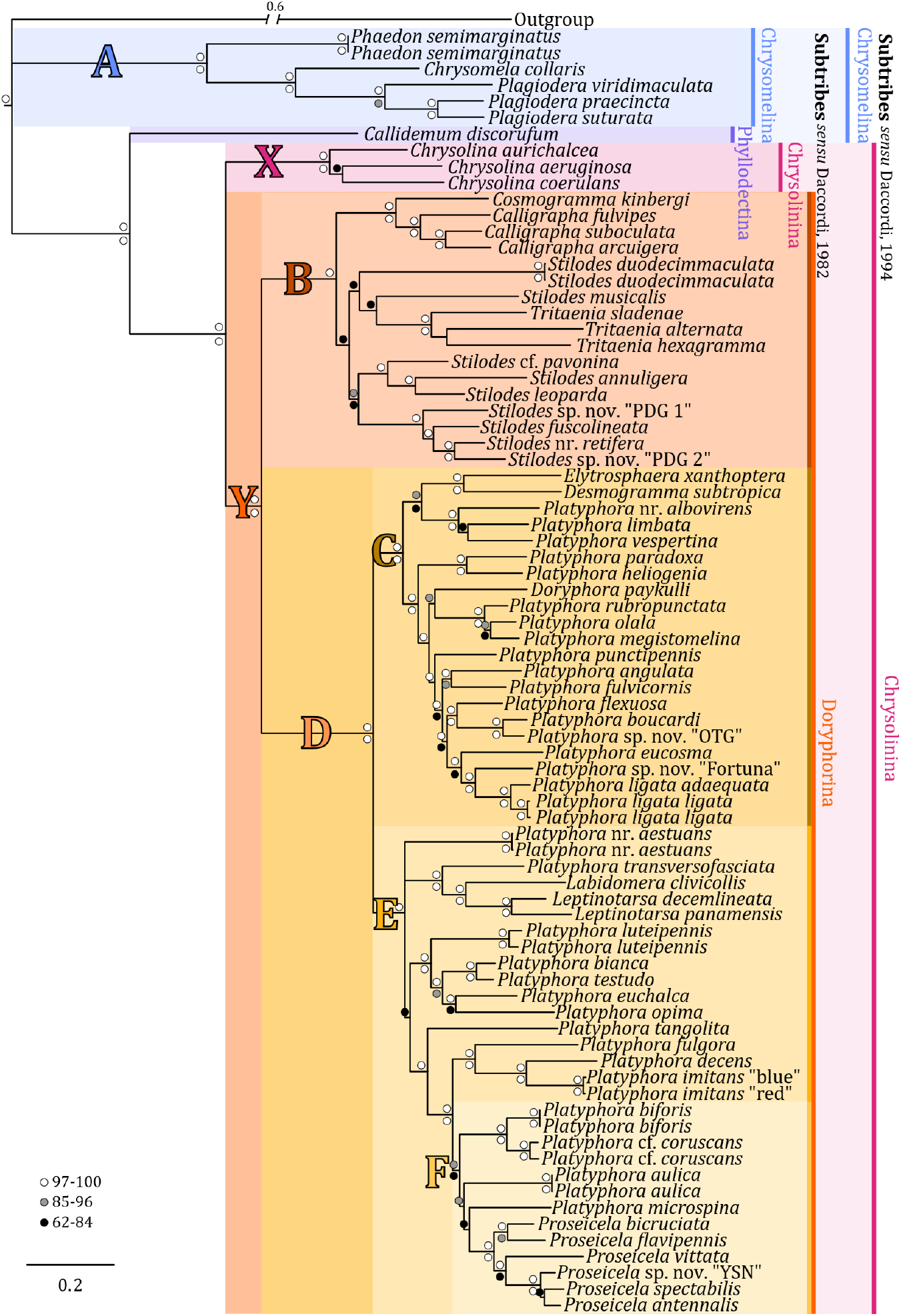
opology and support values of the inferred phylogeny of Neotropical Chrysomelinae through Bayesian analysis of all genes concatenated (with regions of ambiguity from *28S* removed). Posterior probabilities are represented by the circles on top of the nodes and bootstrap values from the maximum likelihood analysis are represented by the circles below the nodes.

### Ancestral host plant use reconstruction

Ancestral states of host plant use were reconstructed using Bayestraits V2 (Pagel et al., 2004). To account for phylogenetic uncertainty, all Bayesian trees from the MrBayes analysis were used in the ancestral state reconstruction after 5% burn-in. We used the most recent common ancestor (mrca) function, a reverse-jump model and set the exponential prior to a uniform hyper prior with the interval 0 to 5 which resulted in an acceptance rate between 20 and 40% as recommended by the developer. We sampled every thousandth generation of 10 million generations after a burn-in of 100000. The ancestral states of host plant use were also reconstructed using maximum parsimony (MP) methods in the StochChar module v.1.1 in Mesquite 3.03 (build 702) (Maddison and Maddison, 2006, 2011).

## Results andDiscussion

### Concatenated gene analyses and taxonomic implications

Bayesian and Maximum Likelihood (ML) analyses of the complete concatenated gene dataset generated similar topologies (Figs 1 and S8, respectively). Specifically, the two analyses differed in clade B (further discussed below) and only the Bayesian analysis recovered basal a polytomy in clade E, although the ML tree recovered short basal branches for clade E with very low support. The five gene segments were also analyzed individually using Bayesian inference (Figs S3 to S7). Gene trees were generally concordant with the concatenated analyses, but the *COII* phylogeny was the least concordant with the concatenated analyses (Fig. S6). However, removal of the *COII* segment from the concatenated dataset analysis did not significantly change the tree topology or our general conclusions. Gene sequences had varying utilities (Fig. S2), with *CAD* being the most useful, probably owing in part to its proportional length in the concatenated dataset (45% when including gaps). Mitochondrial sequences were better at recovering shallowest nodes, while nuclear sequences were better at recovering deeper nodes, which is usual for molecular phylogenies (Moritz et al., 1987). Daccordi’s classifications of Chrysomelinae subtribes (1982 and 1994) are labelled on Fig. 1 to facilitate comparison with the current results. The subtribes (sensu Daccordi, 1982) Chrysomelina (Clade A), Chrysolinina (Clade X), and Doryphorina (Clade Y), and Phyllodectina (represented by a single taxon), were all recovered as robustly supported monophyletic clades (and are further discussed below). In contrast, Gómez-Zurita et al. (2007) did not find these subtribes to be monophyletic: in their study, Doryphorina was polyphyletic, with a paraphyletic Chrysolinina nested within and their polyphyletic Chrysomelina and polyphyletic Phyllodectina were interspersed within each other. That said, of the thirteen nodes informative at the subtribe level, only three had support values above 51%, and therefore their study does not provide robust information pertaining to subtribal relationships.

In contrast to Chrysomelina (sensu Daccordi, 1982), subtribe Chrysomelina (sensu Daccordi, 1994) is paraphyletic because the Australian taxon (*Callidemum discorufum* (Lea)) was recovered as sister to Chrysolinina (Clade X) + Doryphorina (Clade Y) with strong support (posterior probability (PP) = 1.0 and bootstrap = 100) (Fig. 1). Before being placed in Chrysomelina, most of the Australian taxa were placed in the subtribe Phyllodectina (sensu Daccordi, 1982). The subtribe Phyllodectina was synonymized by Daccordi (1994) because he proposed that the analysis of more Australian taxa would likely demonstrate the subtribe to be artificial, based on convergent traits rather than evolutionary history (Weise, 1915, Daccordi, 1994). Daccordi’s argument may be correct, however, given the placement of *Callidemum discorufum*, Daccordi’s (1994) proposed solution is found to be imperfect.

We find the Chrysomelina (sensu Daccordi, 1982) to be monophyletic with *Phaedon* Latreille sister to (*Chrysomela* Linnaeus + *Plagiodera* Chevrolat; clade A, Fig. 1). This agrees with the phylogenetic analysis by Termonia et al. (2001). Since our specimens of *Chrysomela collaris* Linnaeus were from Mongolia and the specimens of *Phaedon* and *Plagiodera* were from Bolivia and Ecuador, this suggests that Chrysomelina had a global range that later split through vicariance events. This hypothesis is also supported by some maps in Daccordi (1994).

### Combining molecular and chemical traits

Chemical analysis of Chrysomelinae defensive secretions may also be an important trait in understanding Chrysomelinae evolution as suggested by Pasteels (1993). These defensive secretions are released from exocrine glands of chrysomelines and are either synthesized by the beetles or derived from precursors that are widely distributed in plants (Pasteels et al., 2003). As a consequence, their defensive chemicals are rarely constrained by host plant, increasing their potential utility for inferring taxonomic grouping (Pasteels et al., 2003). Even more importantly for taxonomic inferences, the compounds require biosynthetic pathways and metabolic mechanisms that are very distinct, and therefore unlikely to be the result of convergent or parallel evolution (Pasteels et al., 2003). The defensive secretions of several of the insect taxa studied here have been previously characterized.

Species in subtribe Chrysomelina (sensu Daccordi, 1982; clade A in Fig. 1) secrete nitropropanoic acid and isoxazolinone glucosides (Pasteels et al., 1994, 2004). Subtribe Chrysolinina (sensu Daccordi, 1982), represented in our study by the genus *Chrysolina* Motschulsky (clade X, Fig. 1), is monophyletic and sister to Doryphorina (sensu Daccordi, 1982). Within the Doryphorina, species in clade B secrete cardenolides or polyoxygenated steroids (Daloze et al., 1991, 1995, Pasteels et al., 1982, 2003, Timmermans et al., 1992), and species in clade D secrete triterpene saponins (Timmermans et al., 1992, Pasteels et al., 2001, 2003, 2004, Plasman et al., 2000a, 2000b). Doryphorina was merged into Chrysolinina by Daccordi (1994), since we find that Doryphorina (sensu Daccordi, 1982) to be monophyletic (clade Y, Fig. 1) and distinct from Chrysolinina in terms of its defensive chemistry, we therefore suggest that Doryphorina maintain its subtribe status. Thus, for the remainder of the paper we will restrict comparisons of our results to Daccordi’s (1982) classification.

Daccordi (personal comm. in Pasteels et al., 2004) noted that the genera of Neotropical chrysomelines are badly in need of revision. However, this task is hindered by frequent convergent evolution in the subfamily (Daccordi et al., 1999). For example, whether tarsal claws are fused or connate tarsal claws (*i.e.*, claws that are in contact with each other for much of their length but not fused) are used as a diagnostic criterion: genera *Doryphora*, and *Tritaenia* have connate tarsal claws (Clark et al., 2024), whereas many other genera have fused tarsal claws (Sampaio and da Fonseca, 2023). This trait also varies within genus *Calligrapha*: subgenus *Zygogramma* Chevrolat has connate tarsal claws, but all other subgenera of *Calligrapha* have fused tarsal claws (Clark et al., 2024). Our phylogeny suggests this character may be more labile than anticipated and not ideal as a distinguishing character.

Doryphorina (Clade Y, PP and bootstrap ≥ 99) is separated into clades B and D (Fig. 1). Clade B contains (*Cosmogramma* Erichson + *Calligrapha* Chevrolat) sister to (*Stilodes* Chevrolat + *Tritaenia* Motschulsky). Our findings completely support the taxonomic changes proposed by Clark et al. (2024) of treating species of the former genus *Zygogramma* as either species of *Calligrapha* subgenus *Zygogramma* or as members of genus *Tritaenia.* Our analysis included members of three the current six subgenera of *Calligrapha*, namely subgenus *Zygogramma* with *C. arcuigera* Stål (Clark et al., 2024), subgenus *Calligrapha* with *C. fulvipes* (Gistel) (Gómez-Zurita, 2021), and subgenus *Erythrographa* Gómez-Zurita with *C. suboculata* Stål (Gómez-Zurita, 2018). Our analysis also included three species of *Tritaenia*; these were recovered nested within *Stilodes* (Fig. 1); however, Bayesian support for these branches is relatively low (75 ± 5) and specific topology differed between the ML and Bayesian analyses. In the ML analysis (Fig. S8), we recovered (*Cosmogramma* + *Calligrapha*) as nested within *Stilodes* rather than sister to *Stilodes*. Finally, our analyses consistently suggest that *Stilodes* is paraphyletic with *Tritaenia* nested within.

Clade D was further divided into two large clades C and E (PP and bootstrap = 100; Fig. 1). These species secrete triterpene saponins (Timmermans et al., 1992, Pasteels et al., 2001, 2003, 2004, Plasman et al., 2000a, 2000b). One potential exception to this is *Proseicela antennalis* (Kirsch); preliminary analysis of its secretions tentatively detected cardenolides and did not find triterpene saponins (Pasteels et al., 2004). In the light of *Proseicela*’s phylogenetic placement in a clade secreting triterpene saponins and not cardenolides, *Proseicela* secretions should be studied again (J. Pasteels, personal comm. March 2013). Clade C, contained (*Desmogramma* Erichson + *Elytrosphaera* Chevrolat) sister to (18 spp. of *Platyphora* + *Doryphora paykulli* (Stål)). Clade E contained a polytomy of three branches: (*Platyphora* nr. *aestuans* (Linnaeus)), (*Platyphora transversofasciata* (Jacoby), *Labidomera clivicollis* (Kirby)*, Leptinotarsa decemlineata* (Say) *+ Leptinotarsa panamensis* Tower), and (13 *Platyphora* spp. + 6 *Proseicela* spp.), this polytomy is absent in the ML analysis (Fig. S8), in which *Pl.* nr. *aestuans* is sister to a clade containing all other species of clade E. This polytomy does not affect the monophyly of the genera in the clade, and *Platyphora* is polyphyletic in both cases. In Gómez-Zurita et al. (2007), *Proseicela* is sister to *Doryphora* but support was under 50% for this in their parsimony tree, and those two species are sister to (*Leptinotarsa* Chevrolat + *Labidomera* Chevrolat), with similarly low support. We recovered *Leptinotarsa* and *Proseicela* as well-supported monophyletic clades (PP and bootstrap = 100; Fig. 1) nested within *Platyphora*. Recovered within clade E, was a well-supported clade consisting of all Solanaceae-feeding species included in the analysis (clade F), except *Leptinotarsa* spp. The Bayes factor test the likelihood of *Platyphora* monophyly gave a log difference between constrained and unconstrained trees of 426.9. A log difference above five is considered very strong evidence in favor of the better model (Kass and Raftery, 1995). In other words, there is very strong evidence for rejecting monophyly of *Platyphora*. This polyphyletic state of *Platyphora* is unlikely to be new, as the genus seems to be delimited from other genera using homoplastic traits and mostly defined by symplesiomorphies.

Flowers (2004) synonymised *Stilodes* and *Leptinotarsa*, and gave *Stilodes* priority over *Leptinotarsa* (see Dury & Jackson, 2026). In light of our results, members of *Stilodes* and *Leptinotarsa* clearly fall in different subclades of Doryphorina (Fig. 1). The log difference between the constrained tree for monophyly of *Stilodes* and *Leptinotarsa* from the Bayes factor test is 462.5; very strong evidence rejecting the synonymy of *Leptinotarsa* and *Stilodes*. While most authors rejected this synonymy, we nonetheless formally restore genus *Leptinotarsa* Chevrolat (type species *Leptinotarsa heydeni* Stål) as distinct from *Stilodes* Chevrolat (type species *Chrysomela humeralis* Gory). This means that *Stilodes decemlineata* (Say) is an invalid combination and, fortunately for the applied entomological literature, we restore the combination *Leptinotarsa decemlineata* (Say)—this species is likely the most economically significant pest of all the Chrysomelidae.

Another example of morphological convergence is the presence or absence of a mesosternal projection. The mesosternum of *Platyphora* and *Doryphora* is modified into a forward projecting horn or spine, with some species only having a small tubercle (Bechyné and Springlová de Bechyné, 1965). Our phylogeny suggests that presence of this horn is the ancestral state for the clade (clade D in Fig. 1) but was secondarily lost in (*Desmogramma* + *Elytrosphaera*), in (*Labidomera* + *Leptinotarsa*), and in *Proseicela*. The mesosternum of *Proseicela* species is produced into a short lobe (Daccordi et al., 1999), *Pr. bicruciata* has one of the larger mesosternal projection forming a small tubercle. It is, thus, logical that *Platyphora microspina* (Bechyné)— named for the diminutive size of its sternal horn, reduced in that species to a short thick spine—is sister to *Proseicela*. The function of the sternal horn is still unclear but Eberhard (1981) has observed male *Doryphora* sp. near *punctatissima* using it in male aggressive behaviours. Females also have a horn, which could help them compete for food plants, although this has not been observed (Eberhard, 1981).

Termonia et al. (2002) studied defensive secretions of *Platyphora* in a phylogenetic context and inferred a clear clade in which all species sequester pyrrolizidine alkaloids and host-plant derived triterpene saponins. We identified a similar topology for this clade (clade C) with five of the same species included in our study (*Pl. vespertina* (Baly), *Pl. heliogenia* Bechyné*, Pl. boucardi* (Jacoby), *Pl. ligata adaequata* (Bechyné) = their *Pl. ligata*, and *Pl. eucosma* (Stål)). Our topology for species that do not sequester pyrrolizidine alkaloids (clade E) is distinct from that obtained by Termonia et al. (2002) (*Pl. transversofasciata* = their *Pl. salviny, Pl. tangolita* (Bechyné) = their *Pl. decorata, Pl. microspina,* and *Pl. opima* (Stål)) (Fig. 1). Despite these differences, the main findings of Termonia et al. (2002) remain valid: “dual sequestration could be the key mechanistic means by which transitions among ecological specializations (i.e., restricted host-plant affiliations) are made possible.” Overall, chemical traits reported in the literature support our molecular taxonomic classifications of the Chrysomelinae.

### Ancestral host plant use

Chrysomelinae have long been used to study host specificity in herbivorous insects (Strauss, 1988). Given the numerous species not sampled, it is difficult to estimate the number of host plant switches that have occurred in the Chrysomelinae. For example, *Calligrapha* species feed on Malvaceae, but also feed on at least nine other plant families (Clark et al., 2004). Our ancestral host plant reconstruction estimated the likely host plant of the common ancestor of all Chrysomelinae (Fig. 2) but this estimate should not be taken as informative; it may be a sampling artefact, since genus *Timarcha*, sister to subtribe Chrysomelina (*sensu* Daccordi, 1982), feeds on Rosaceae, Rubiaceae and Plantaginaceae, and species of Chrysomelina feed on Salicaceae or Asteraceae, but also on Betulaceae, Brassicaceae, Polygonaceae, Malvaceae, and many other families (Clark et al., 2004, Jolivet and Hawkeswood, 1995, Termonia et al., 2001). Despite the uncertainty of host associations at the deepest nodes, host switches at more shallow nodes can be inferred when a clade containing species that eat plants from a certain family is either nested within, or derived from, a clade eating plants from a different family (Fig. 2). Most of the plant families appear to have been colonized once, with two exceptions: Malpighiaceae colonized by (*Platyphora bianca* (Bechyné) + *Pl. testudo* (De Mey)) and (*Stilodes* + *Tritaenia*) and Solanaceae colonized by *Leptinotarsa* and (*Platyphora* spp. + *Proseicela*). The common ancestor of *Leptinotarsa* likely fed on Asclepiadoideae, like its sister genus *Labidomera*, while species of the *Leptinotarsa* now mainly feed on Solanaceae and sometimes Zygophyllaceae or Asteraceae (Clark et al., 2004, Hsiao, 1988). In *Leptinotarsa*, Asteraceae-feeding represents a reversal to a more ancestral food plant. Other reversals to Asteraceae-feeding are apparent in a few clades of genus *Platyphora*: (*Pl. angulata* (Stål) + *Pl. fulvicornis* (Guérin-Méneville)) and (*Pl. eucosma* (Stål) + *Pl. ligata* (Stål)). The alternative appears less likely: these two clades may have maintained Asteraceae-feeding and be sister to three clades, (*Doryphora* + *Platyphora* sp.), (*Pl. punctipennis* (Jacoby)) and (*Pl. boucardi* (Jacoby)), which all independently switched to Apocynoideae (Fig. 2).

**Figure 2.**
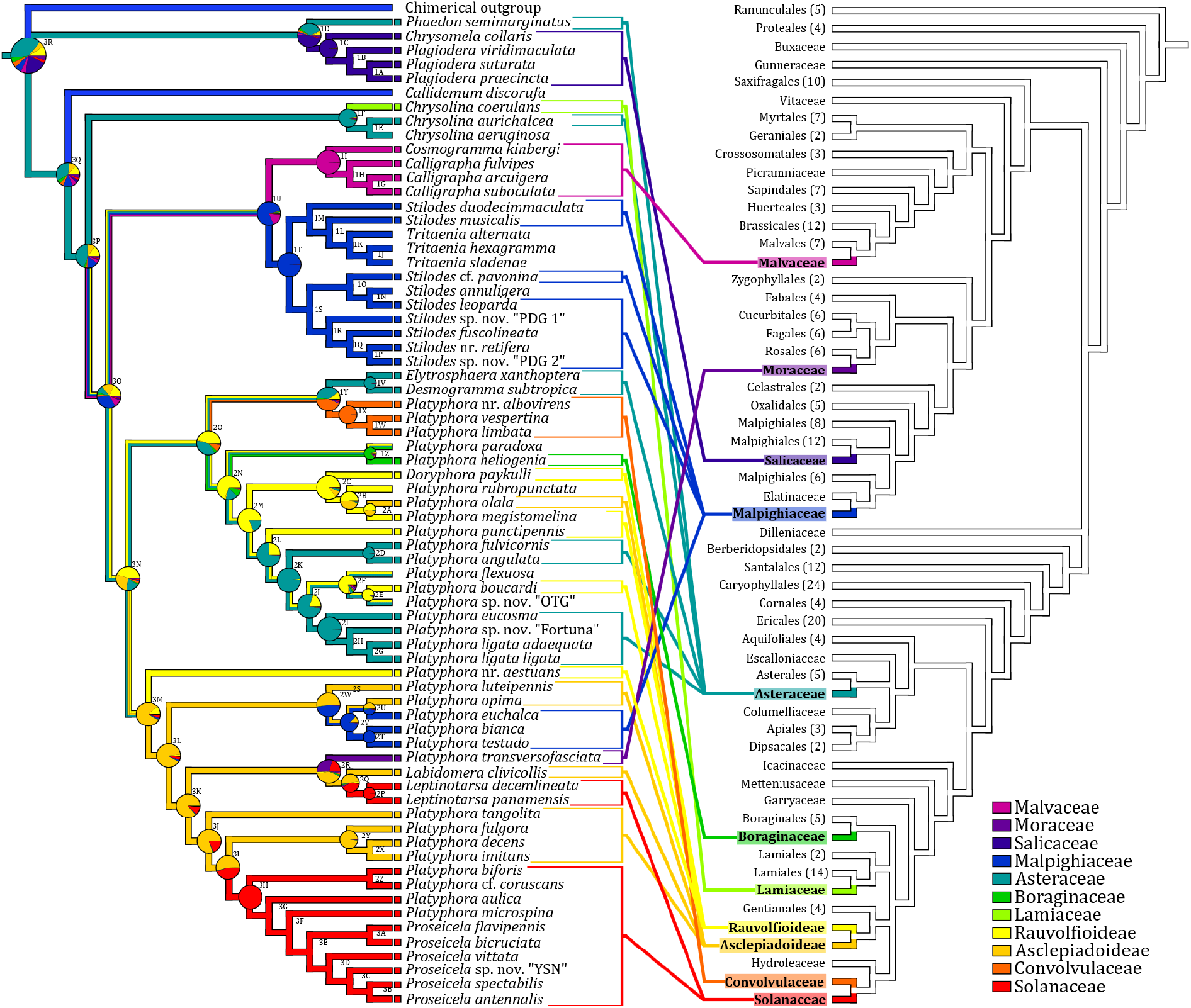
Reconstruction of ancestral host plant families (left) on a maximum a posteriori probability (MAP) Bayesian tree of Neotropical Chrysomelinae from analysis of the concatenated gene dataset. Phylogenetic relations of the host plant families (right) according to version 13 of the Angiosperm Phylogeny Group website. Colours on branches are from Maximum Parsimony reconstruction, pie charts at nodes represent likelihood of each ancestral state as reconstructed in Bayestraits for the most recent common ancestor of taxa at the tips. Squares at the tips of branches on the beetle cladogram indicate available data.

Broadly, the phylogeny of the beetles does not follow the phylogeny of the host plants; there are several switches to host plants very distant from each other, for example, Asteraceae (Asterales) to Convolvulaceae (Solanales) and Asclepiadoidae (Gentiales) to Malpighiaceae (Malpighiales) (Fig. 2 and APG III 2009). Despite having access to over 240 plant families (left side of Fig. 2), to our knowledge, Chrysomelinae of the Neotropics only feed on plants in nine of those families.

## Conclusion

Using a five gene molecular phylogeny, we revealed the phylogenetic relationships among 70 in-group taxa of largely Neotropical Chrysomelinae. The resulting phylogenetic hypothesis identified problems with the most recent arrangement of Chrysomelinae genera and subfamilies. *Stilodes* was paraphyletic with respect to *Tritaenia*. Genus *Platyphora* was polyphyletic and clearly divided into two clades, one with *Doryphora* nested within and sister to *Desmogramma* and *Elytrosphaera*; the other clade contained a polytomy between *Labidomera, Leptinotarsa* and other *Platyphora* species in which the genus *Proseicela* was nested. Our phylogeny is supported by chemical defense traits and underscores the need for a revision of the Chrysomelinae genera of the Americas, especially *Platyphora*. The results provide a first phylogenetic framework for the study of chemical, morphological, and natural history traits in these leaf beetles.

## Author contributions

**Guillaume J. Dury:** Conceptualization; methodology; investigation; data curation; formal analysis; validation; project administration; funding acquisition; resources; writing - original draft; writing - review & editing. **Donald M. Windsor:** Conceptualization; methodology; investigation; data curation; validation; project administration; supervision; funding acquisition; resources; writing - review & editing. **Barbara J. Sharanowski:** Methodology; formal analysis; resources; validation; writing - review & editing. **Lukáš Sekerka:** Data curation; validation; writing - review & editing. **Jacqueline C. Bede:** Conceptualization; methodology; validation; project administration; supervision; funding acquisition; resources; writing - review & editing.

## Acknowledgments

We are grateful to C. Galdames (STRI) for aid in the identification of host plants and to M. Daccordi (Museo Civico di Storia Naturale) for aid in identification of leaf beetles. We thank the following scientist for specimens: J. Vasconcellos-Neto (Universidade Estadual de Campinas) for Brazilian species, J. Pasteels (Université Libre de Bruxelles) for specimens from France and French Guiana, and Y. Pelletier (Agriculture and Agri-Food Canada) for potato beetle specimens. We thank M. Hersh and K. Ashfar for assistance with molecular work, M. Jackson for assisting in the interpretation of the International Code of Zoological Nomenclature. For help in Ecuador, we thank C. Kiel and G. Onore (Pontificia Universidad Católica del Ecuador), and staff of the Yasuní Research Station and Yanayacu Biological Station and Center for Creative Studies. We thank J. Ledezma and J. Aramayo (Museo de Historia Natural Noel Kempff Mercado) for institutional support and assistance in obtaining collecting and export permits in Bolivia (MMAyA-VMA-DGBAP N∘1735/2011, MHNNKM-OF-N∘ 441/2012). Studies in Panama were conducted under the research and collecting permits issued by the Autoridad Nacional del Ambiente (ANAM; SE/A-16-12, SE/A-54-13; DMW principal investigator). Studies in Ecuador were conducted under the research and collecting permits issued by the Ministerio del Ambiente (N°001-10 IC-FAU-DNB/MA and 06-2011-FAU-DPAP-MA). This work was funded by the Natural Sciences and Engineering Research Council of Canada fellowship (CGS M; GJD) and research grant (JCB), and the Fonds de Recherche du Québec Nature et technologies sector (master’s training scholarship; GJD), and DKRVO 2024–2028/5.I.b by Ministry of Culture of the Czech Republic to the National Museum, 00023272 (LS).

