## supplementary materials for "Molecular phylogenetics of Neotropical chrysomeline beetles: Evidence for a constrained history of host plant use"

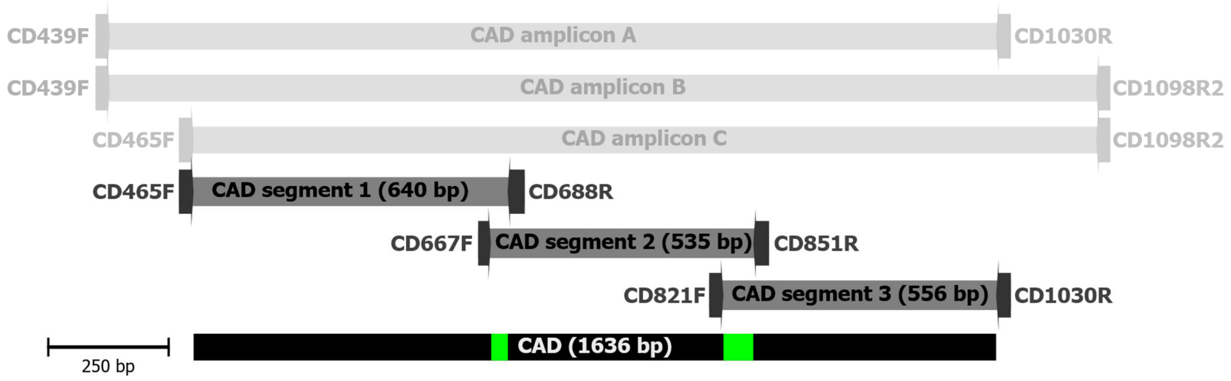


**Figure S1.** *CAD* (*carbamoyl-phosphate synthetase 2, aspartate transcarbamylase and dihydroorotase*) segments used in this study and their respective primers (to scale). The three different amplicons used for fully-nested PCR amplification are represented in pale grey; three CAD gene segments are in dark grey; the combined CAD sequence is represented in black, with internal segment overlap in green (pale grey in greyscale).

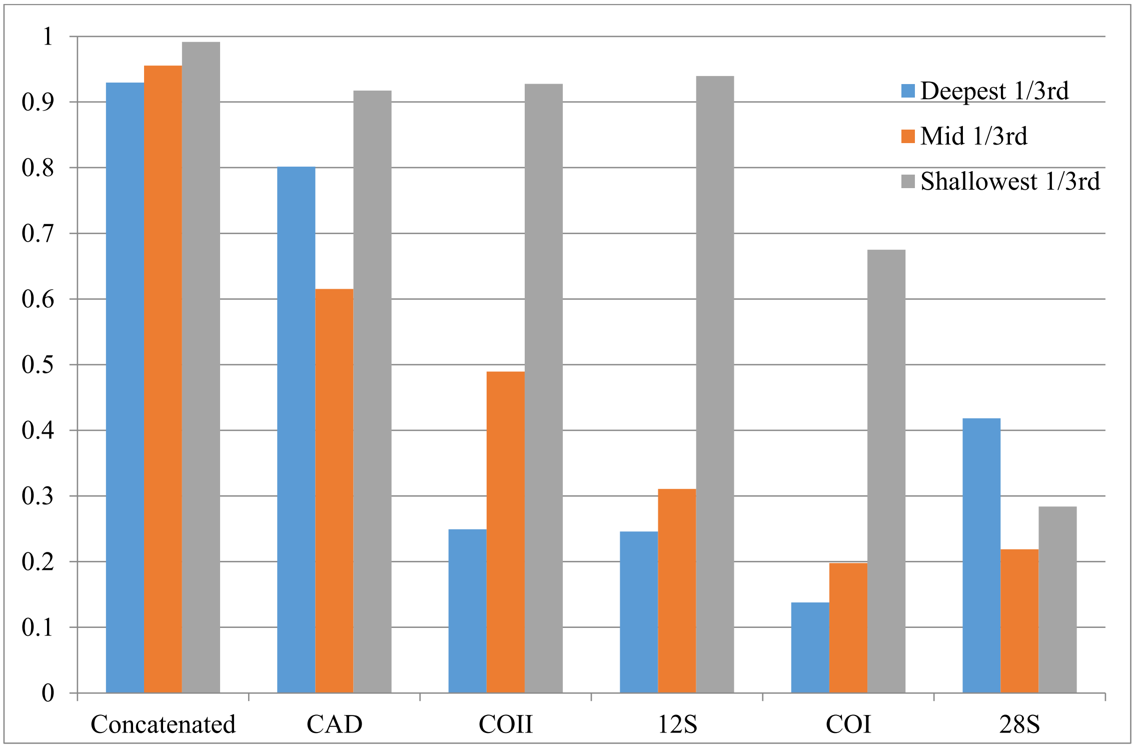


**Figure S2.** Gene performance measured by summing the observed posterior probabilities across the nodes and dividing by the maximum possible posterior probability. Nodes are separated in three groups using number of base substitutions on the concatenated tree, as counted from the tips of the branches. Genes are ordered in terms of average utility.

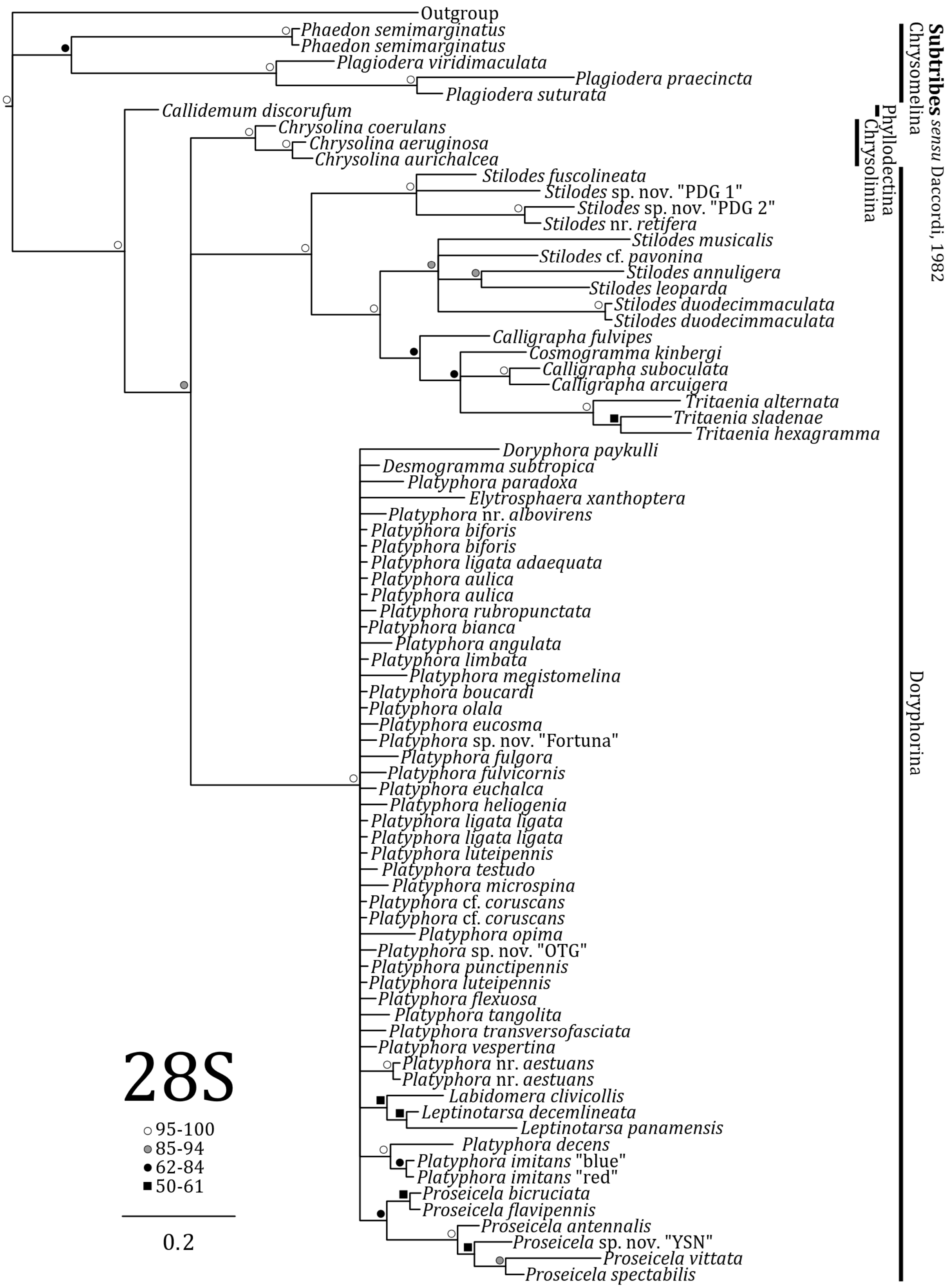


**Figure S3.** Topology and posterior probabilities values of the inferred phylogeny of Neotropical Chrysomelinae through Bayesian analysis of nuclear *28S* ribosomal DNA.

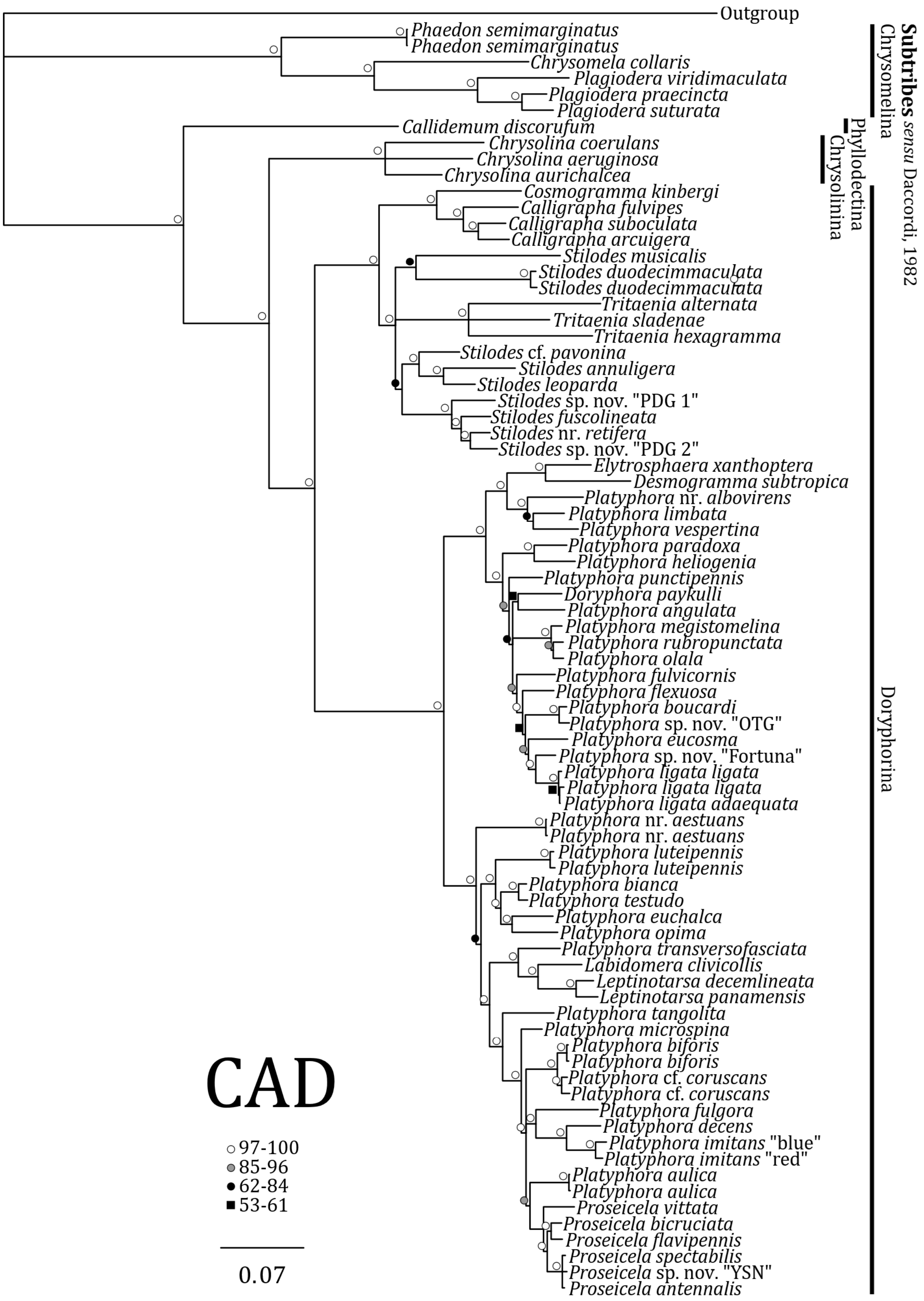


**Figure S4.** Topology and posterior probabilities values of the inferred phylogeny of Neotropical Chrysomelinae through Bayesian analysis of the nuclear protein-coding *CAD* (*carbamoyl-phosphate synthetase 2, aspartate transcarbamylase and dihydroorotase*) amplicons.

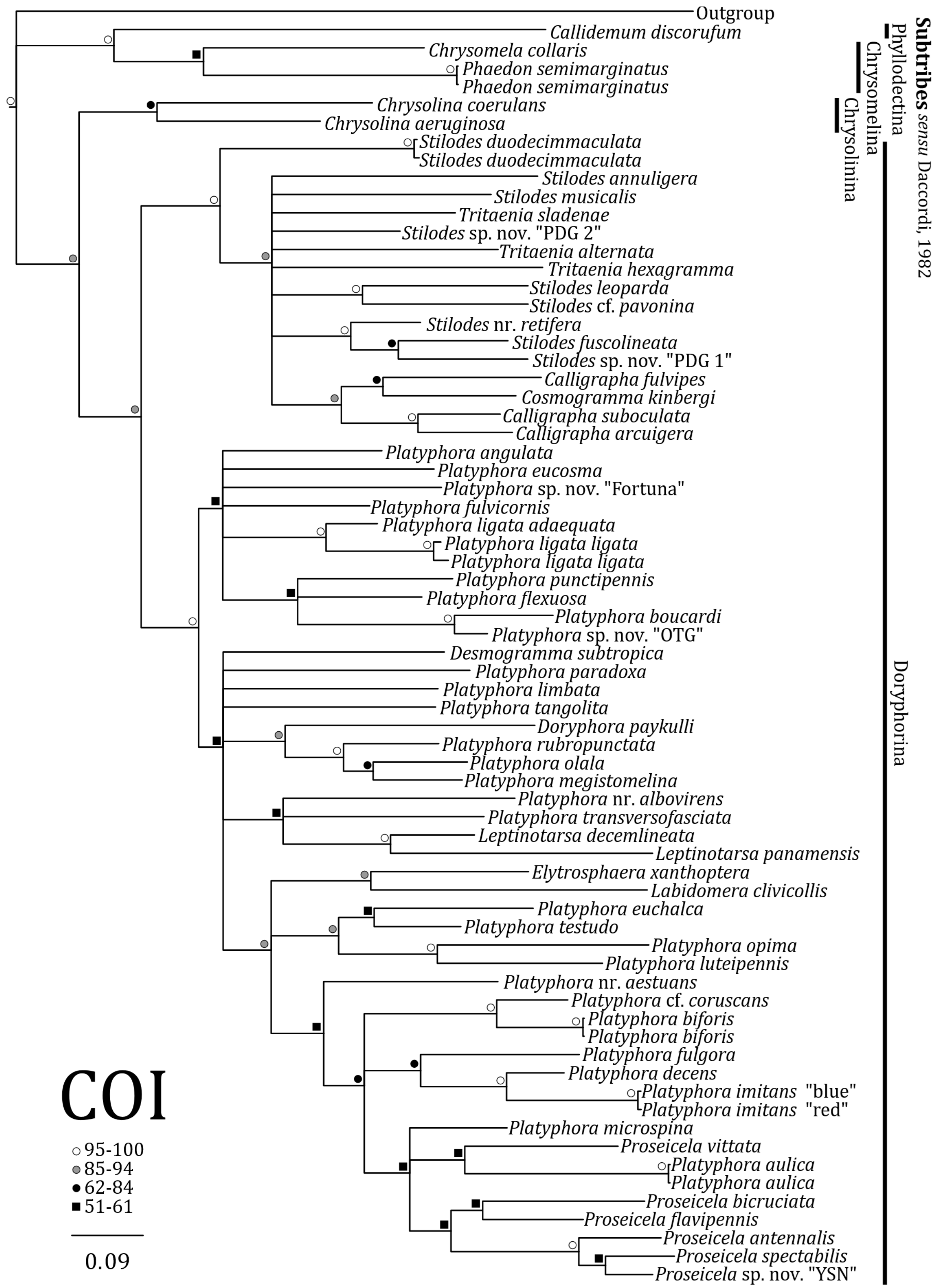


**Figure S5.** Topology and posterior probabilities values of the inferred phylogeny of Neotropical Chrysomelinae through Bayesian analysis of mitochondrial protein-coding *cytochrome oxidase subunit I* (*COI*).

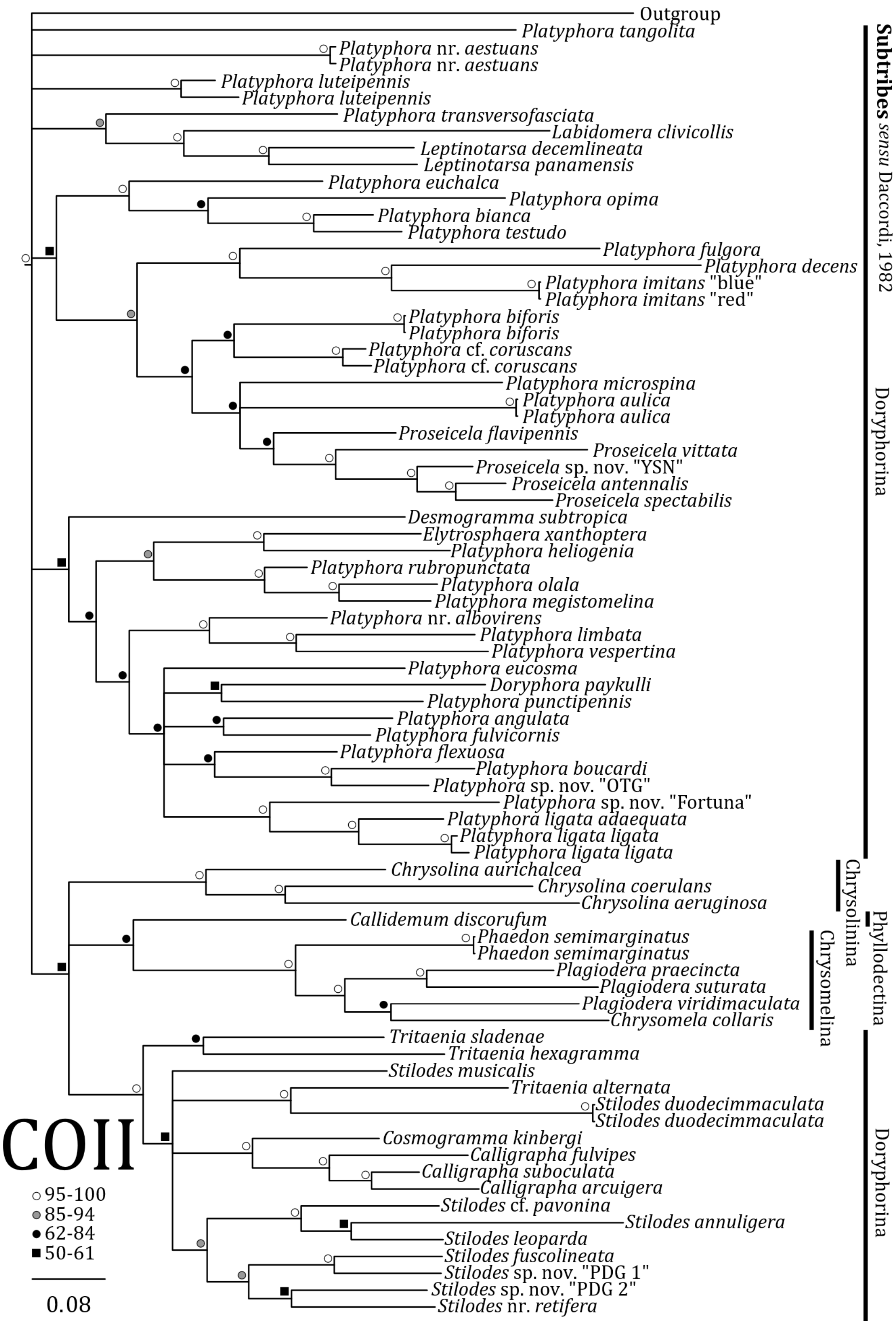


**Figure S6.** Topology and posterior probabilities values of the inferred phylogeny of Neotropical Chrysomelinae through Bayesian analysis of mitochondrial protein-coding *cytochrome oxidase subunit II* (*COII*).

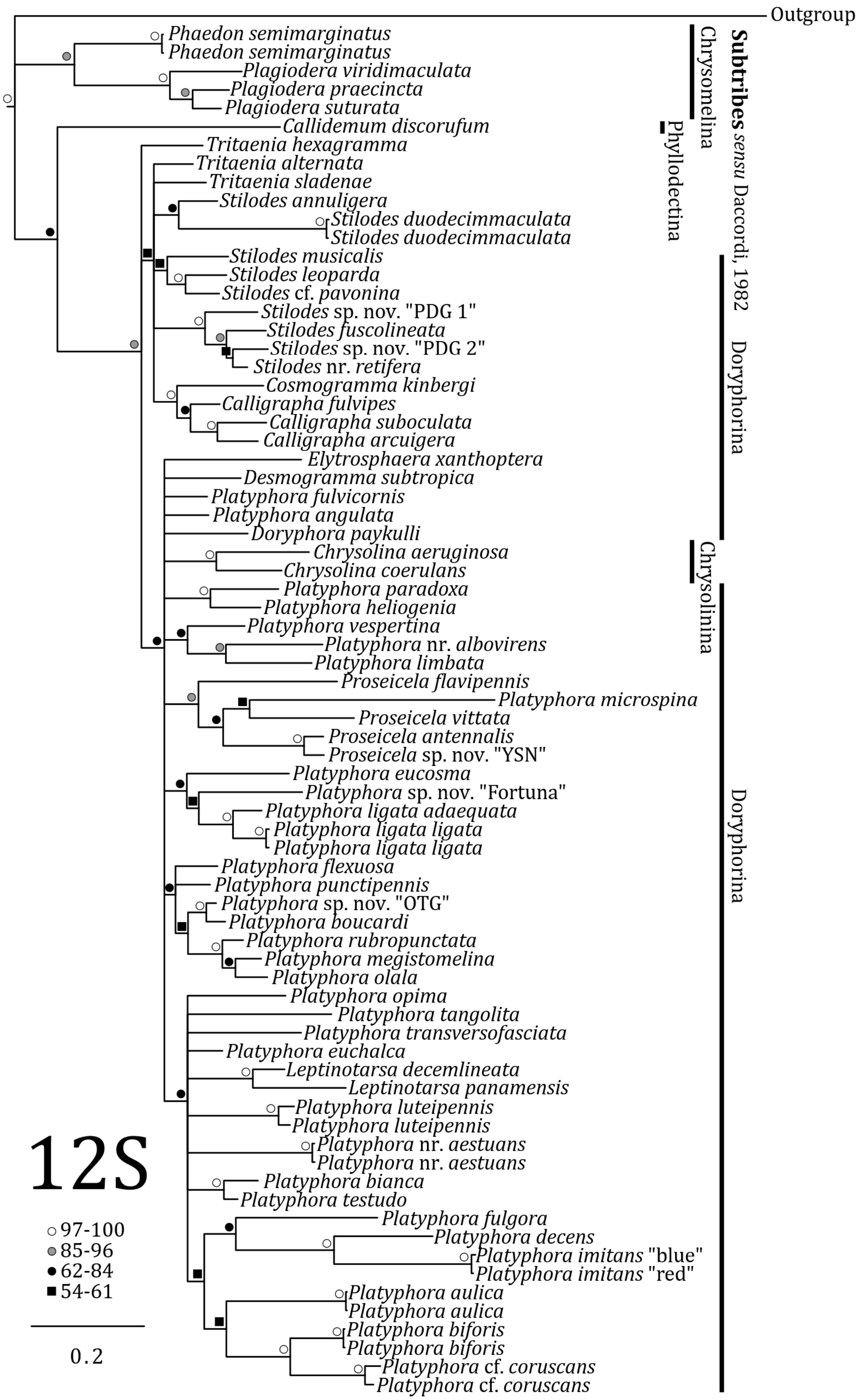


**Figure S7.** Topology and posterior probabilities values of the inferred phylogeny of Neotropical Chrysomelinae through Bayesian analysis of mitochondrial *12S* ribosomal DNA.

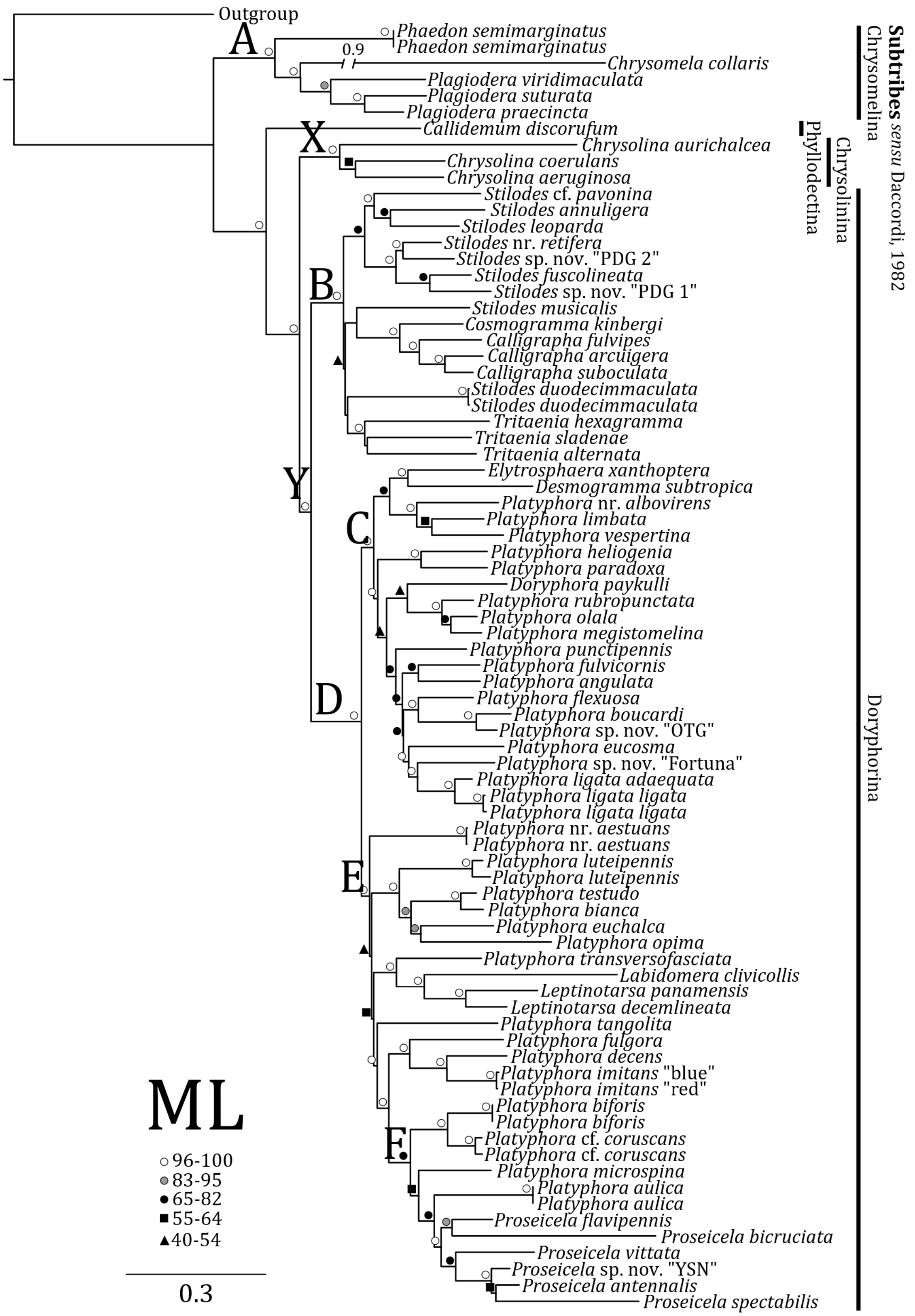


**Figure S8.** Topology and support values of the inferred phylogeny of Neotropical Chrysomelinae through Maximum Likelihood analysis of all genes concatenated (with regions of ambiguity from *28S* removed). Bootstraps values are represented by the shapes on top of the nodes.

**Table S1:** Target regions for amplification and primers used in this study

| Gene segment (length) | | Primer (length) | Primer sequence (5' to 3') | Source |
|---|---|---|---|---|
| *12S* mt rDNA (504-550 bp) | | SR-N-14756 (21-mer) | GAC AAA ATT CGT GCC AGC AGT | Simon *et al.*, (1994) |
| | | SR-J-14233 (20-mer) | AAG AGC GAC GGG CGA TGT GT | |
| *28S* nu rDNA (440-482 bp) | | D2 UP-4 (23-mer) | GAG TTC AAG AGT ACG TGA AAC CG | Gillespie *et al.*, (2003) |
| | | D2 DN-B (21-mer) | CCT TGG TCC GTG TTT CAA GAC | |
| *COI* mt protein-coding (472 bp) | | C1-J-1718F (26-mer) | GGA GGA TTT GGA AAT TGA TTA GTT CC | Simon *et al.*, (1994) |
| | | C1-N-2191 (26-mer) | CCC GGT AAA ATT AAA ATA TAA ACT TC | |
| *COII* mt protein-coding (617 bp) | | modTL2-J-3037 (20-mer) | ATG GCA GAT TAG TGC AWT RG | Termonia *et al.*, (2001) |
| | | modC2-N-3661 (24-mer) | CCA CAA ATT TCW GAA CAT TGA CCA | |
| *CAD* nu protein-coding* (1636 bp) | | CD439F (29-mer) | TTC AGT GTA CAR TTY CAY CCH GAR CAY AC | Wild and Maddison (2008) |
| | 640 bp | CD465F (30-mer) | ACC YAA RAA ART KYT VAT AAT TGG TTC WGG | This study |
| | | CD688R (35-mer) | TGT ATA CCT AGA GGA TCD ACR TTY TCC ATR TTR CA | Wild and Maddison (2008) |
| | 535 bp | CD667F (26-mer) | GGA TGG AAG GAA GTD GAR TAY GAR GT | Wild and Maddison (2008) |
| | | CD851R (32-mer) | GGA TCG AAG CCA TTH ACA TTY TCR TCH ACC AT | |
| | 556 bp | CD821F (29-mer) | AGC ACG AAA ATH GGN AGY TCN ATG AAR AG | Wild and Maddison (2008) |
| | | CD1030R (30-mer) | CWS RGC AYA CCA RTC RAA CTC WAC WGA GCT | This study |
| | | CD1098R2 (29-mer) | GCT ATG TTG TTN GGN AGY TGD CCN CCC AT | Wild and Maddison (2008) |

* *CAD* (*Carbamoyl-phosphate synthetase 2, Aspartate transcarbamylase and Dihydroorotase*) was amplified in three partially overlapping segments using fully nested PCR and primers CD439F or CD465F and CD1098R2 or CD1030R (Fig. S1 in the supplementary material).

**Table S2:** Polymerase chain reaction (PCR) conditions used to amplify the selected gene segments. Amplifications began with an initial 2 minute denaturation step at 94°C and ended with a final extension step of 5 minutes at 72°C.

| Gene | Cycling conditions | | | |
|---|---|---|---|---|
| | Steps* | Temperature (°C) | Time (s) | Cycles |
| *12S* | Denaturation | 93 | 35 | 45 |
| | Annealing | 55 | 35 | |
| | Extension | 72 | 60 | |
| *CAD* amplicon for nested PCR | Denaturation | 94 | 35 | 45 |
| | Annealing | 53 | 60 | |
| | Extension | 72 | 120 | |
| *28S* & *CAD* segment 2 (a)<br>*CAD* segments 1 & 3 (b) | Denaturation | 95 | 30 | 45 |
| | Annealing | 53 (a) or 52 (b) | 60 | |
| | Extension | 72 | 60 | |
| *COII* | Denaturation | 94 | 25 | 45 |
| | Annealing | 52 | 60 | |
| | Extension | 72 | 60 | |
| *COI* | Denaturation | 94 | 30 | 10 |
| | Annealing | 46 | 30 | |
| | Extension | 72 | 30 | |
| | Denaturation | 94 | 30 | 30 |
| | Annealing | 48 | 30 | |
| | Extension | 72 | 40 | |

### Supplementary materials – Photos of species used

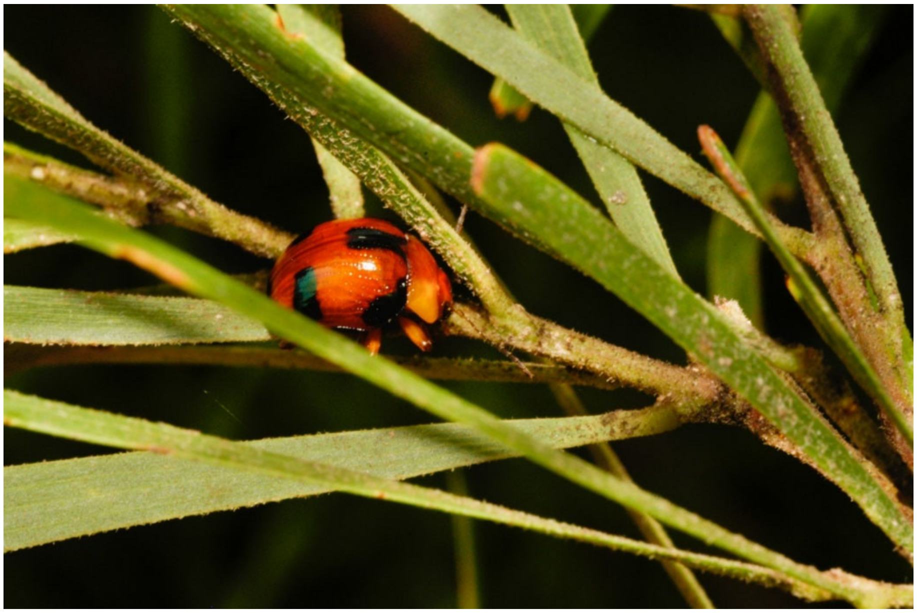

*Callidemum discorufum* (Lea, 1903), photo by DWM.

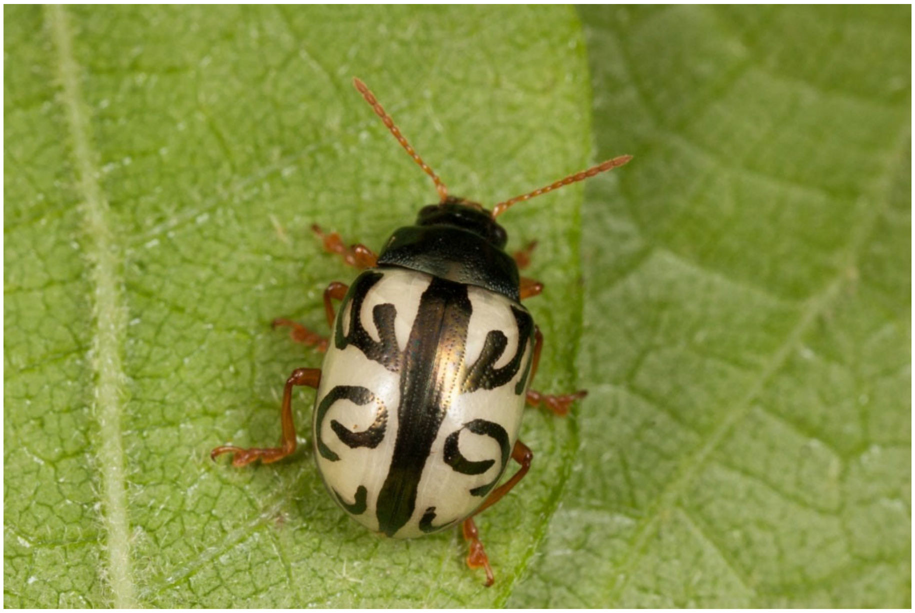

*Calligrapha arcuigera* Stål, 1859, photo by GJD (under CC BY-NC 2.0).

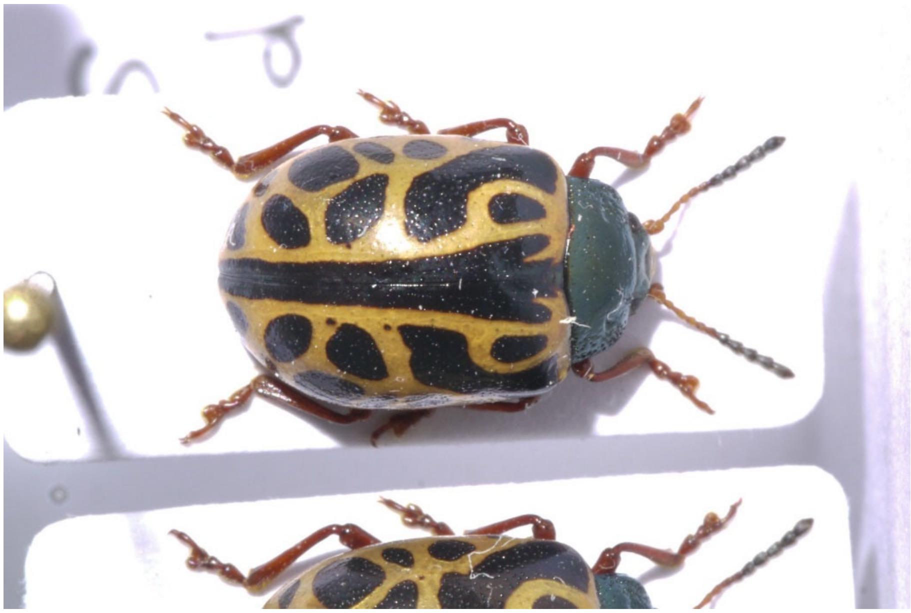

*Calligrapha fulvipes* (Gistel, 1848), photo by LS.

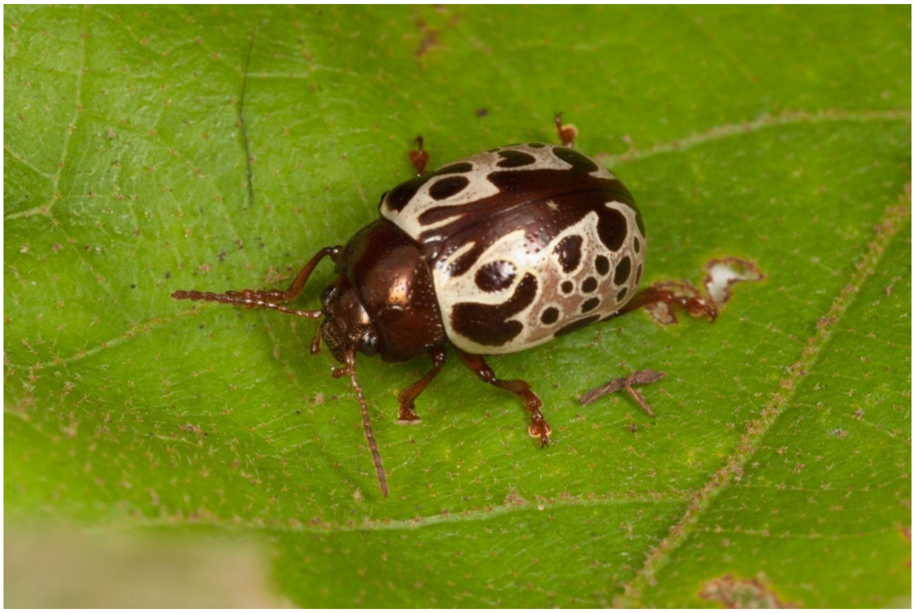

*Calligrapha suboculata* Stål, 1859, photo by GJD (under CC BY-NC 2.0).

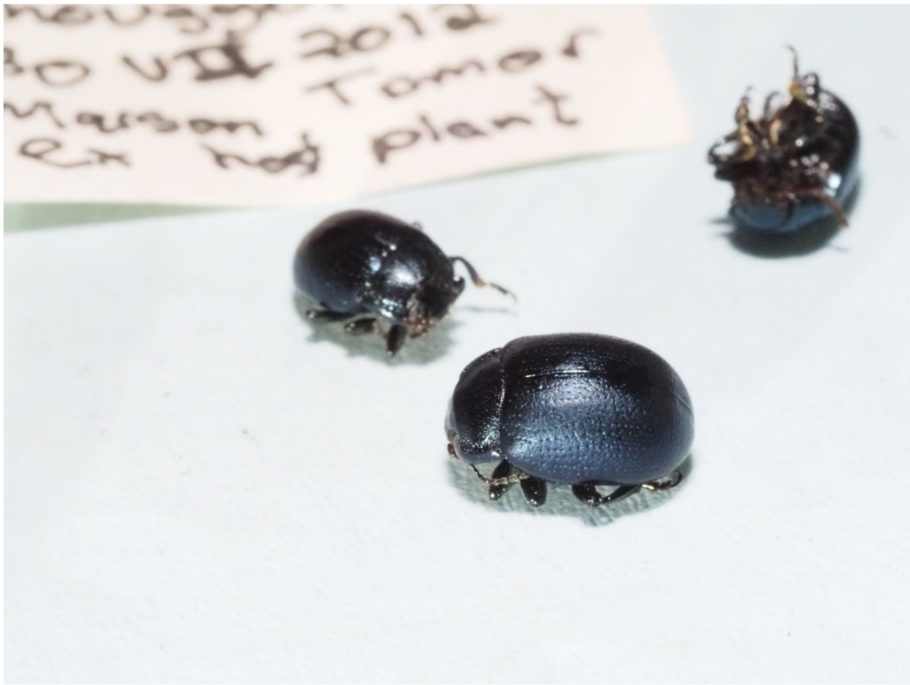


*Chrysolina aeruginosa* (Faldermann, 1835)

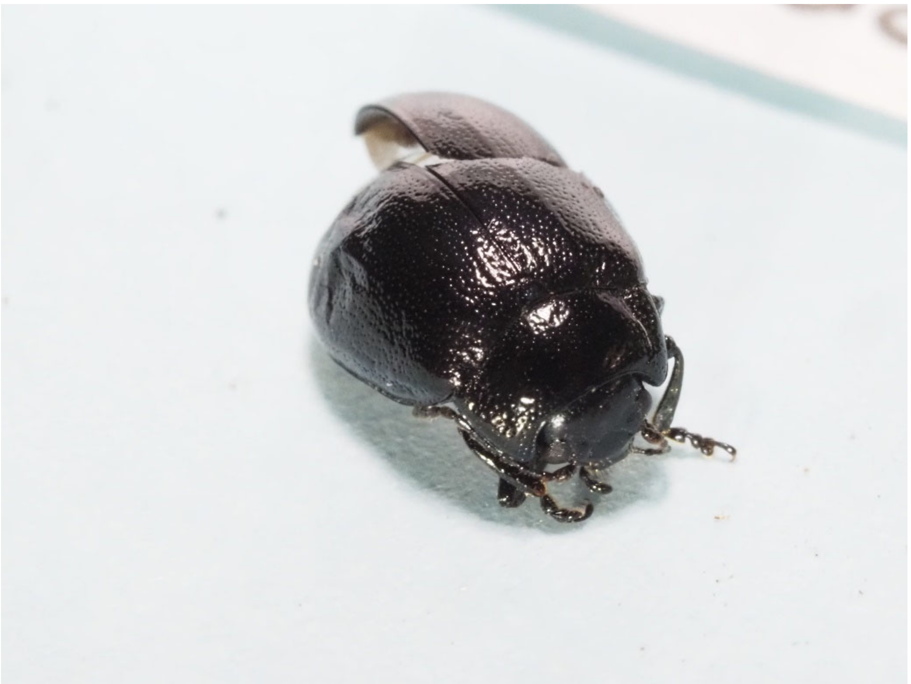

*Chrysolina aurichalcea* (Mannerheim, 1825)

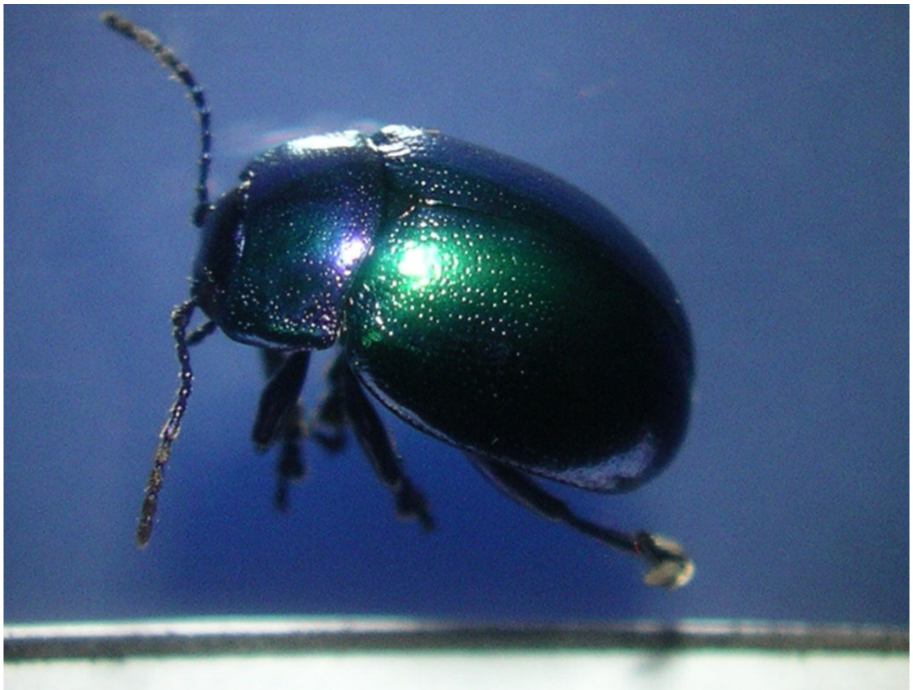

*Chrysolina coerulans* (Scriba, 1791)

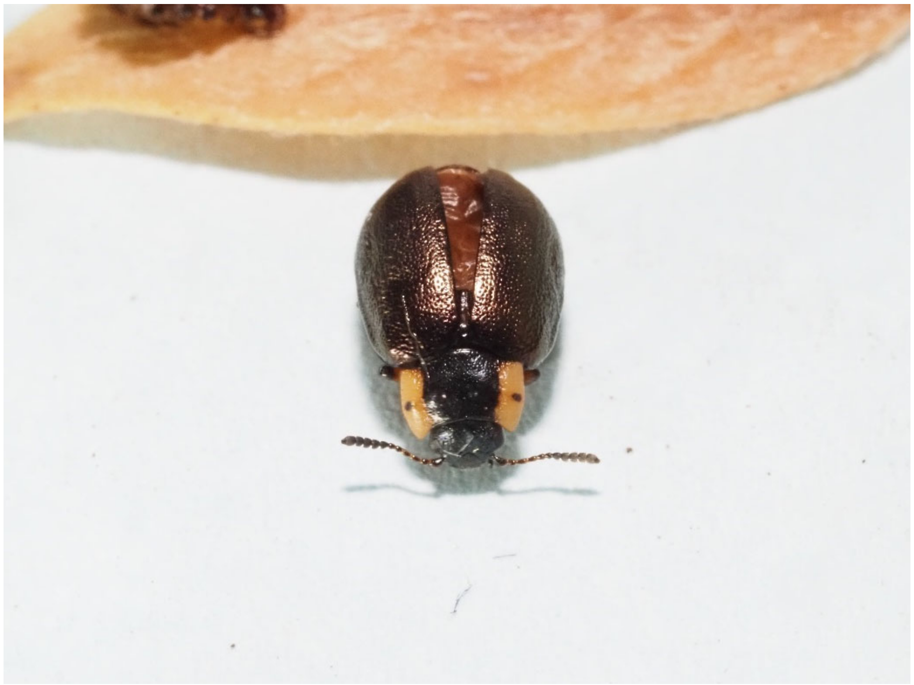

*Chrysomela collaris* Linnaeus, 1758

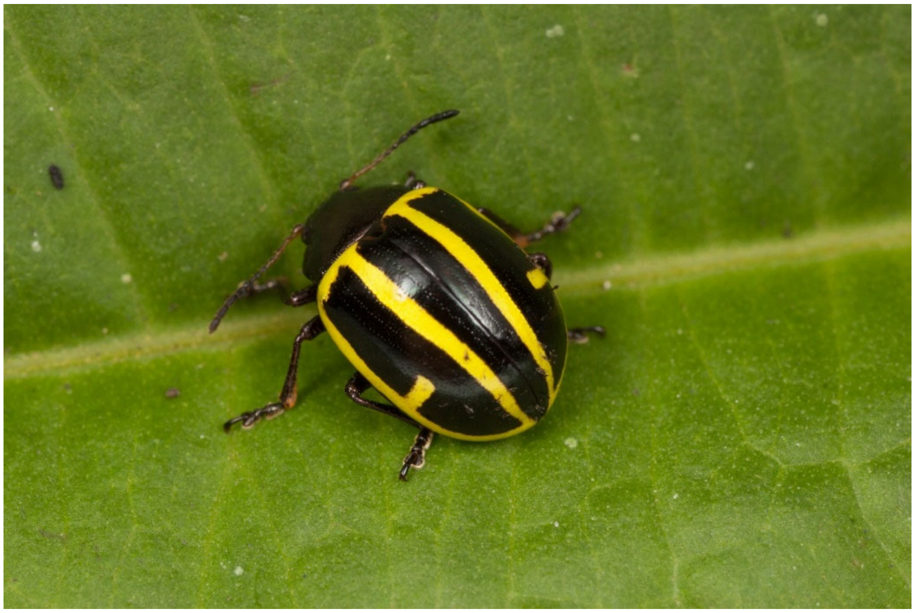

*Cosmogramma kinbergi* (Boheman,1858), photo by GJD (under CC BY-NC 2.0).

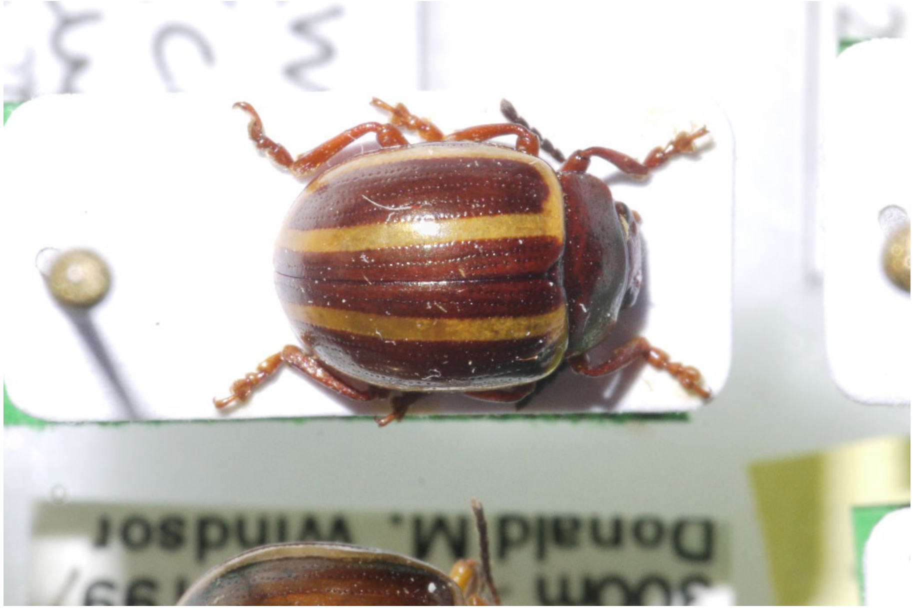

*Desmogramma subtropica* Bechyně, 1946, photo by LS.

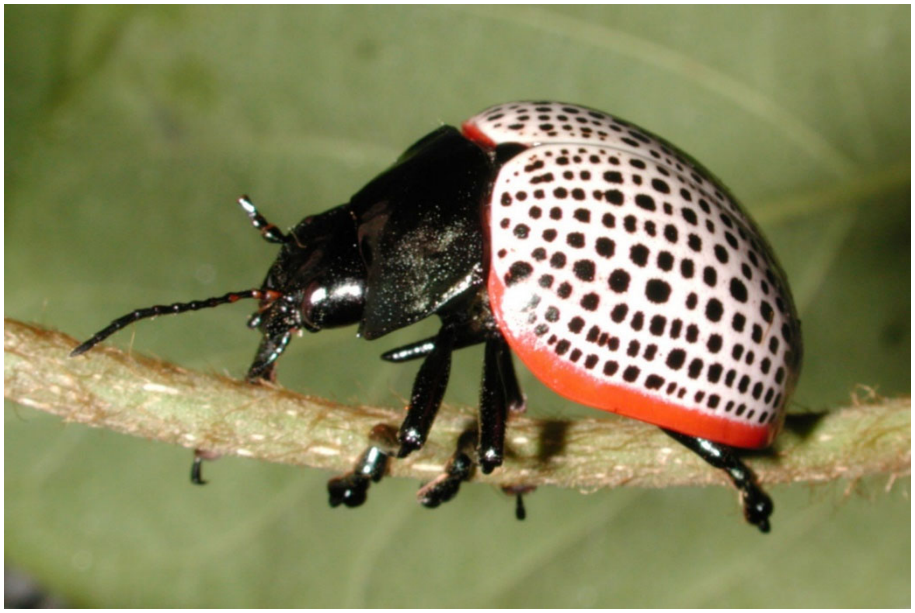

*Doryphora paykulli* (Stål, 1859), photo by DWM.

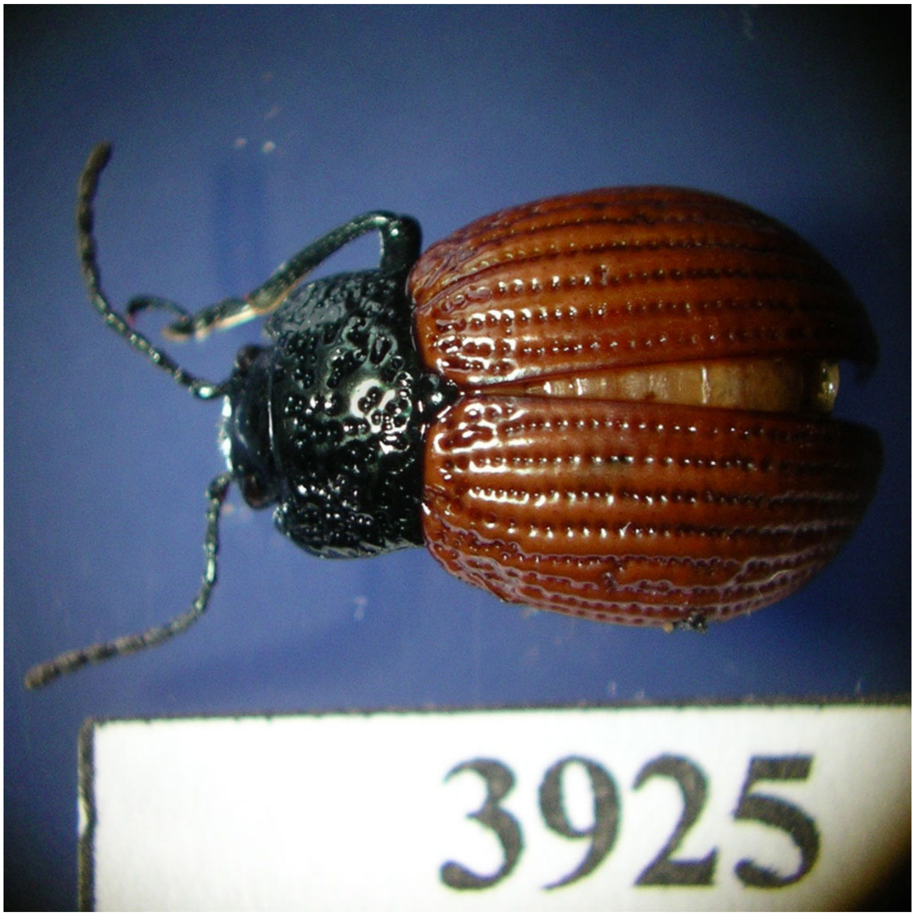


*Elytrosphaera xanthoptera* (Gistel, 1848), photo by DWM.

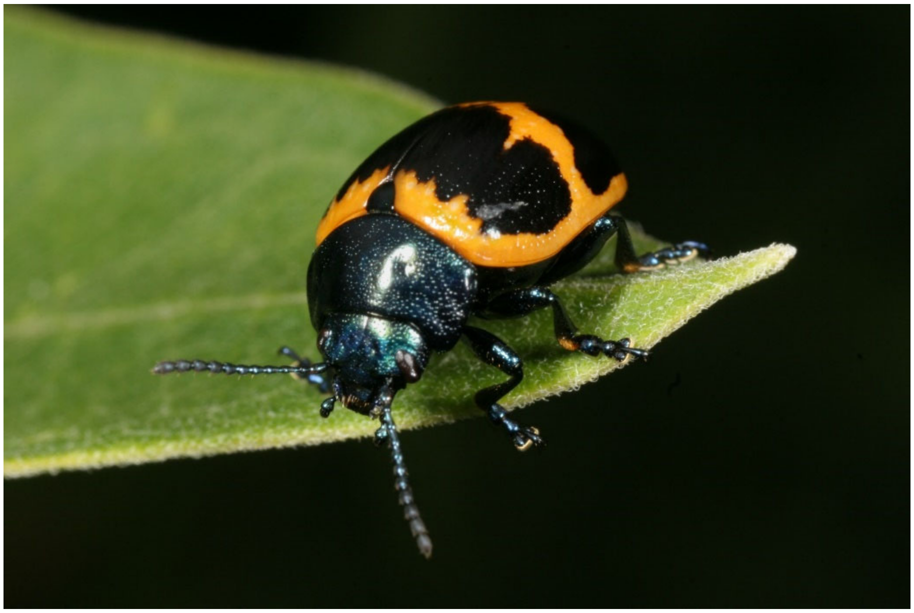

*Labidomera clivicollis* (Kirby, 1837), photo by GJD (under CC BY-NC 2.0).

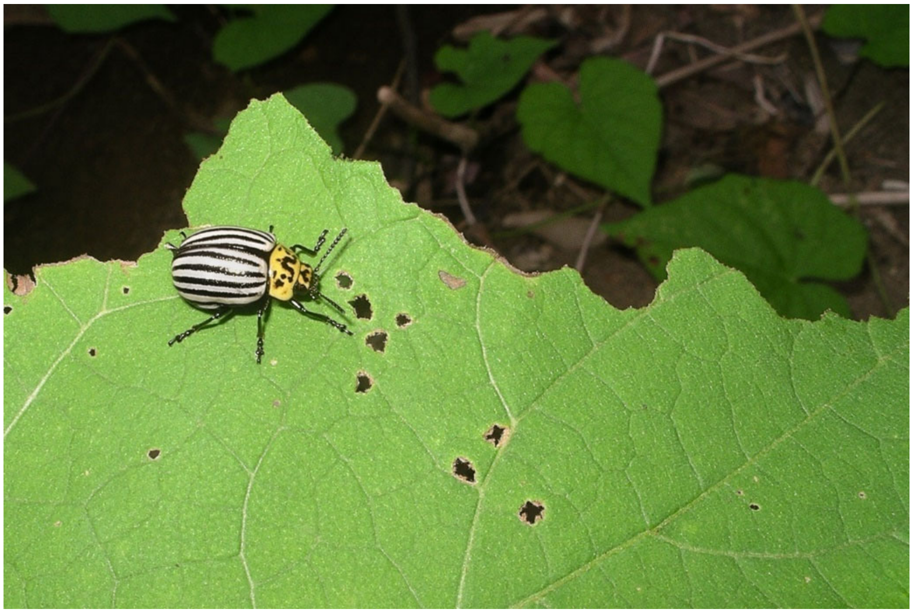

*Leptinotarsa panamensis* Tower, 1918, photo by DWM.

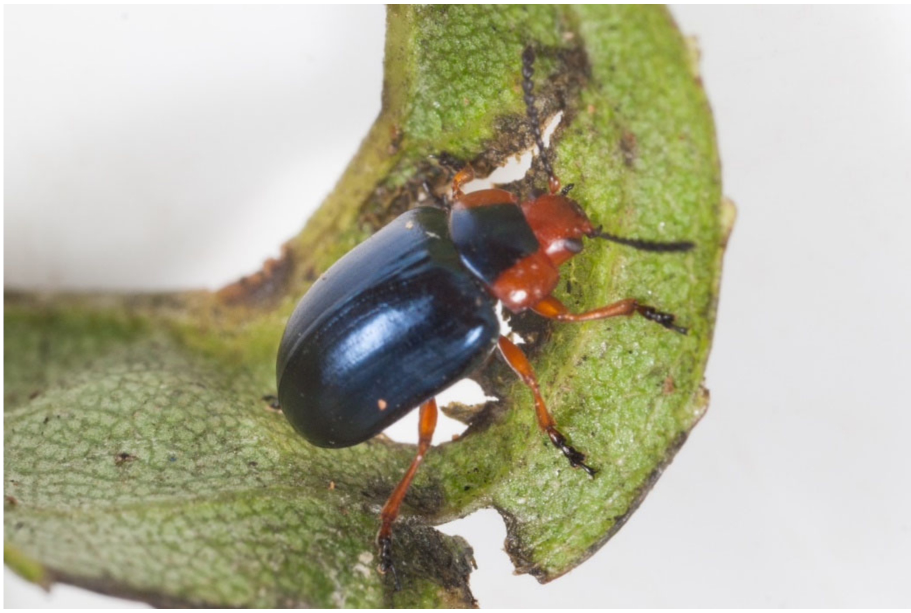

*Phaedon semimarginatus* (Latreille, 1811), photo by GJD (under CC BY-NC 2.0).

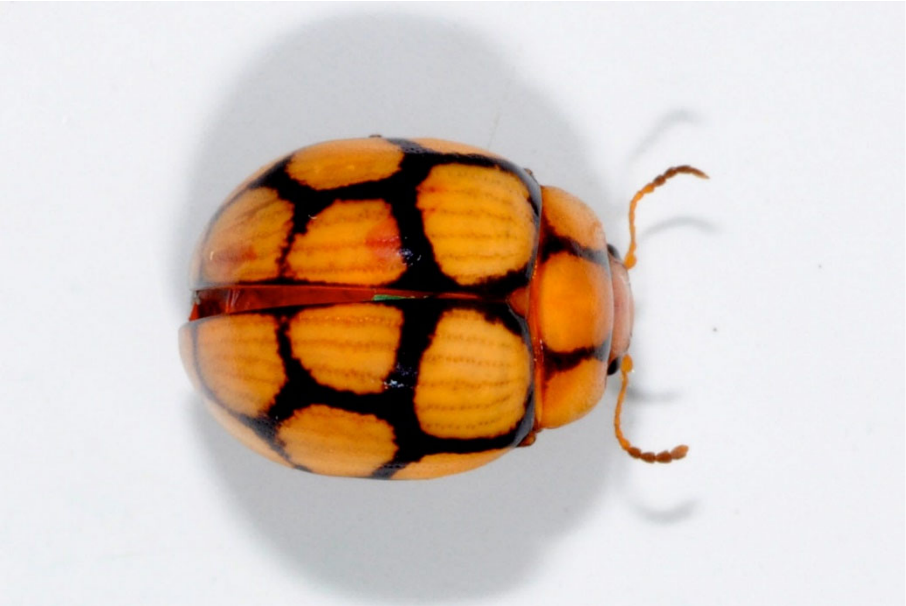

*Plagiodera praecincta* Erichson, 1847, photo by DWM.

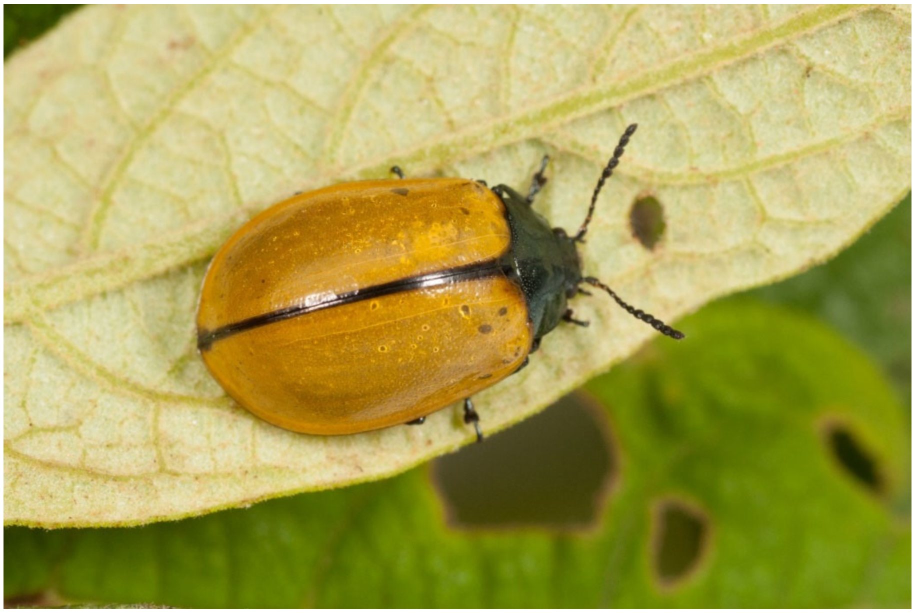

*Plagiodera suturata* Achard, 1925, photo by GJD (under CC BY-NC 2.0).

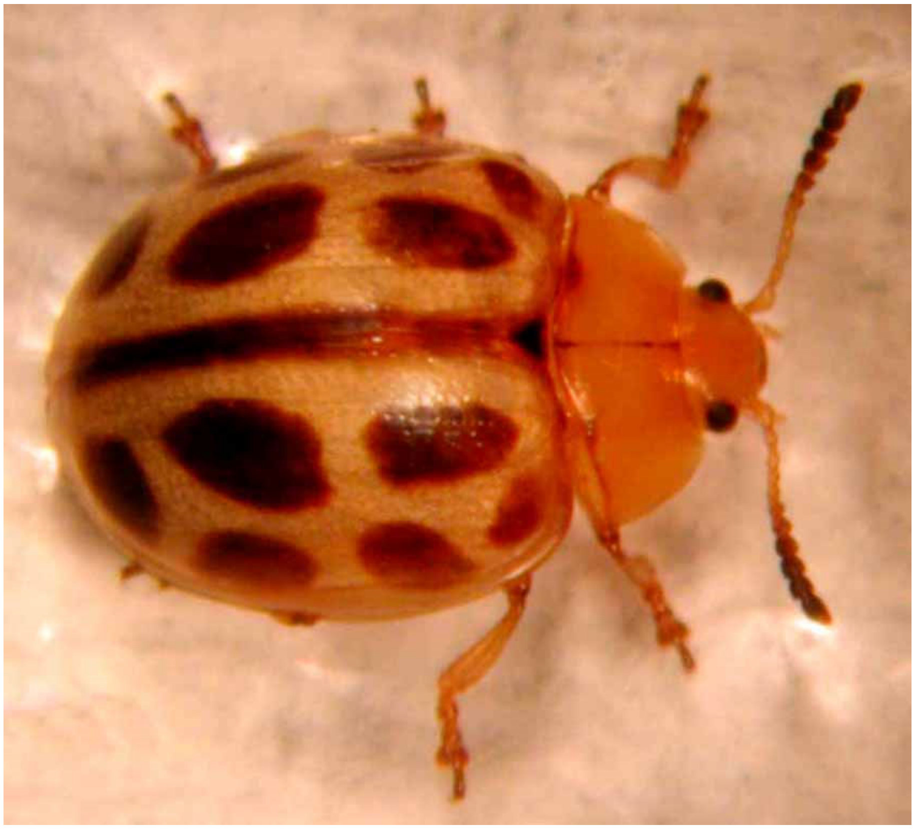

*Plagiodera viridimaculata* Jacoby, 1891, photo by DWM.

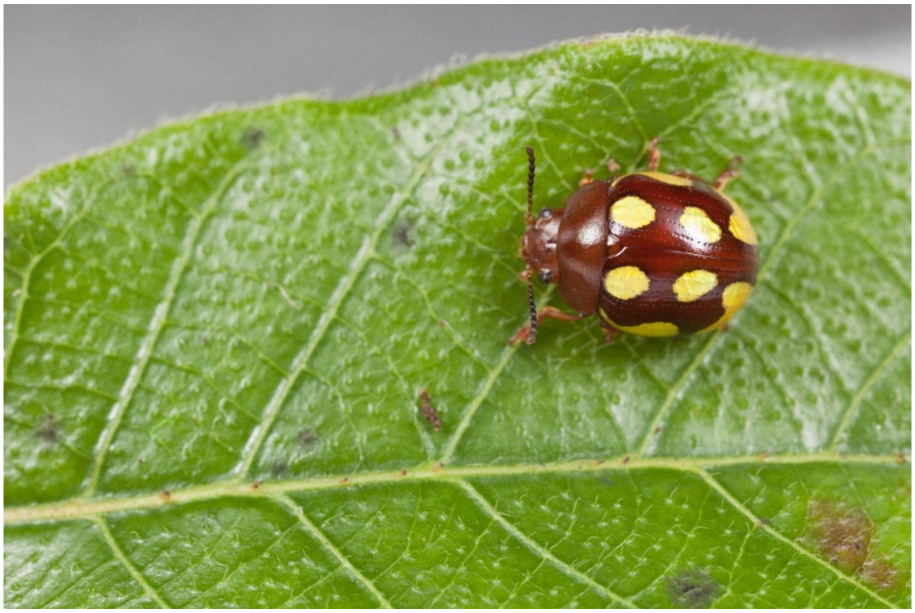

*Platyphora* nr. *aestuans* (Linnaeus, 1758), photo by GJD (under CC BY-NC 2.0).

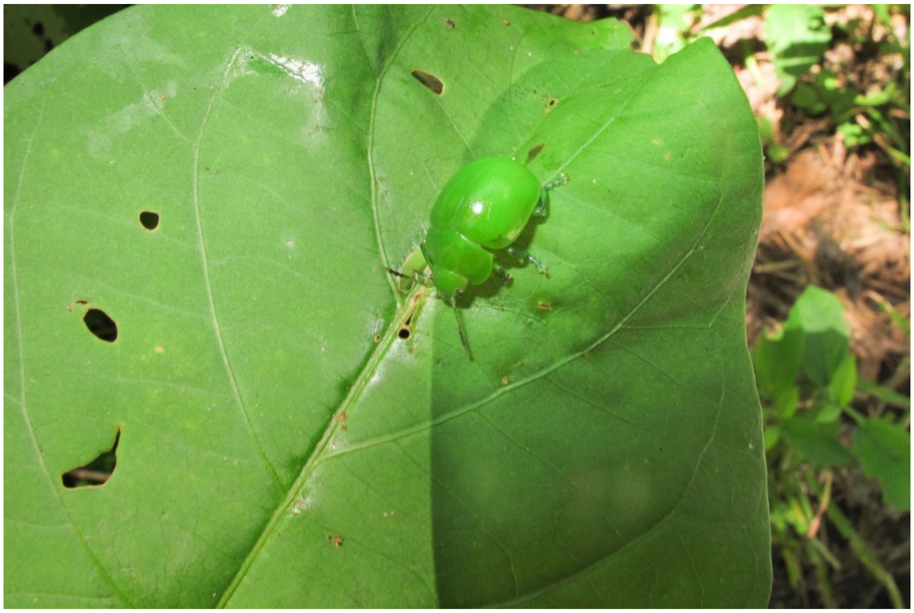

*Platyphora* nr. *albovirens* (Stål, 1857), photo by DWM.

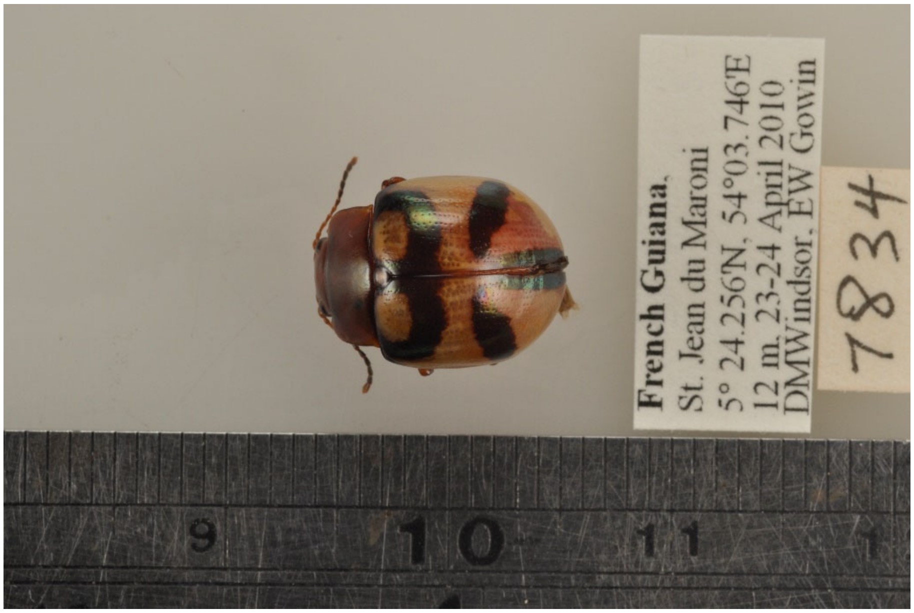

*Platyphora angulata* (Stål, 1858), photo by DWM.

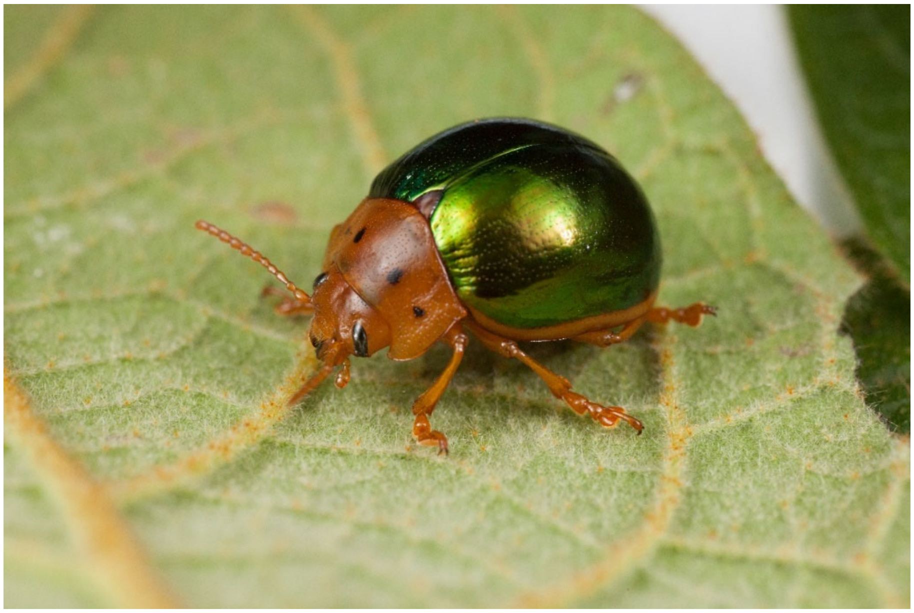
*Platyphora aulica* (Olivier, 1807), photo by GJD (under CC BY-NC 2.0).

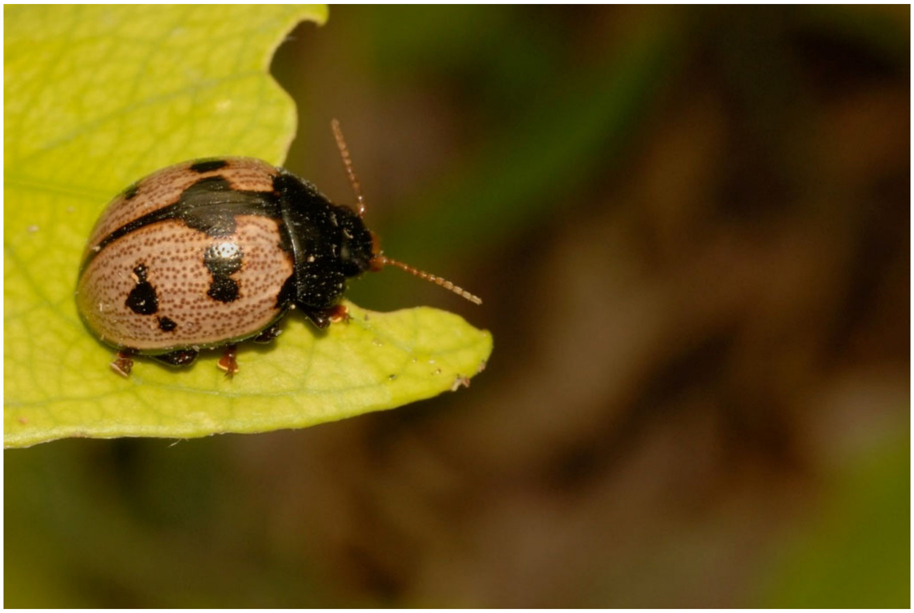

*Platyphora bianca* (Bechyné, 1954), photo by DWM.

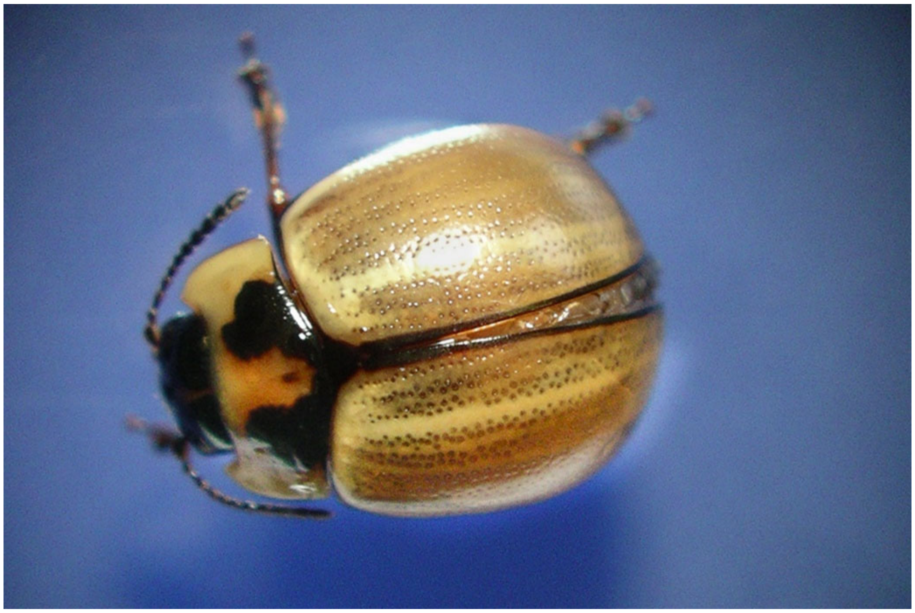

*Platyphora biforis* (Germar, 1823), photo by DWM.

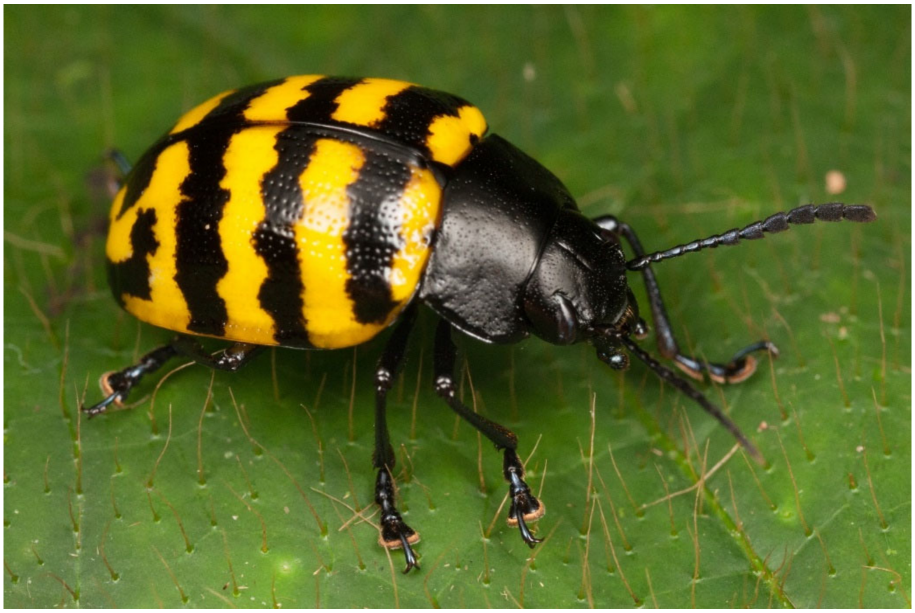

*Platyphora boucardi* (Jacoby, 1883), photo by GJD (under CC BY-NC 2.0).

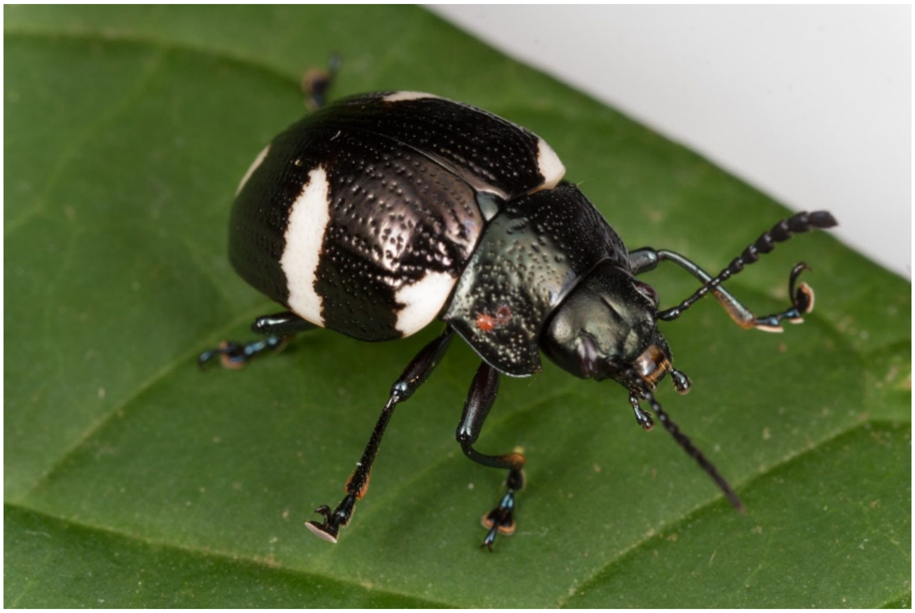

*Platyphora* sp. nov. “OTG”, photo by GJD (under CC BY-NC 2.0).

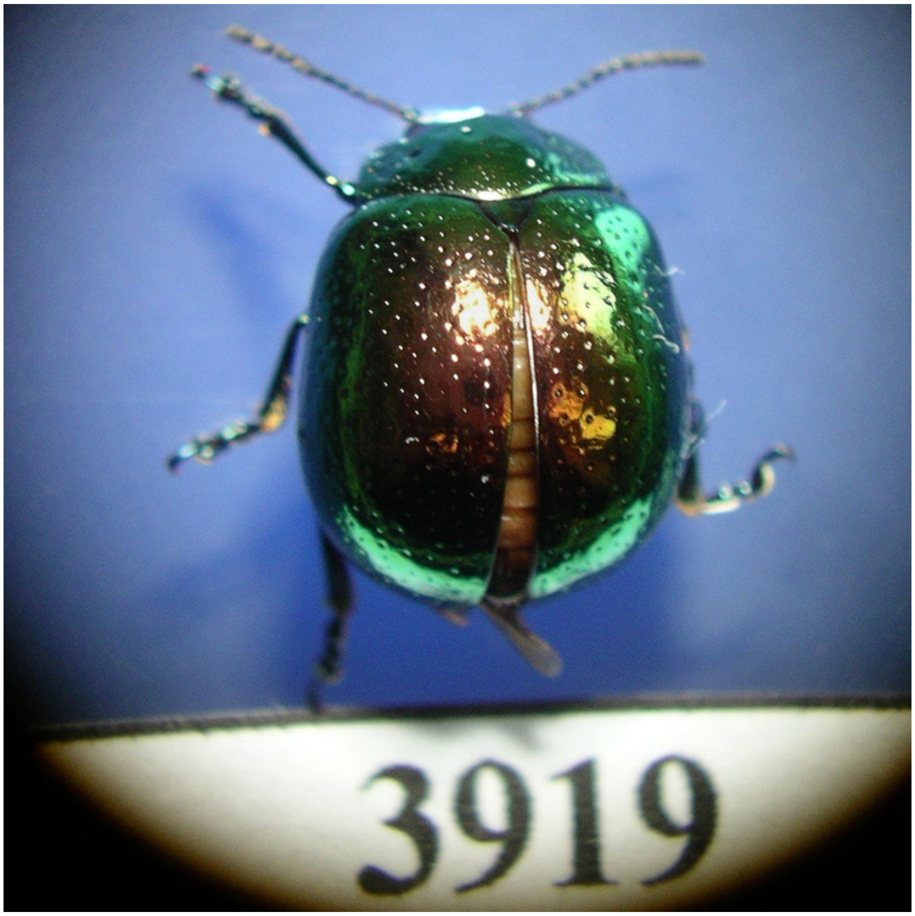


*Platyphora* cf. *coruscans* (Stål, 1858), photo by DWM.

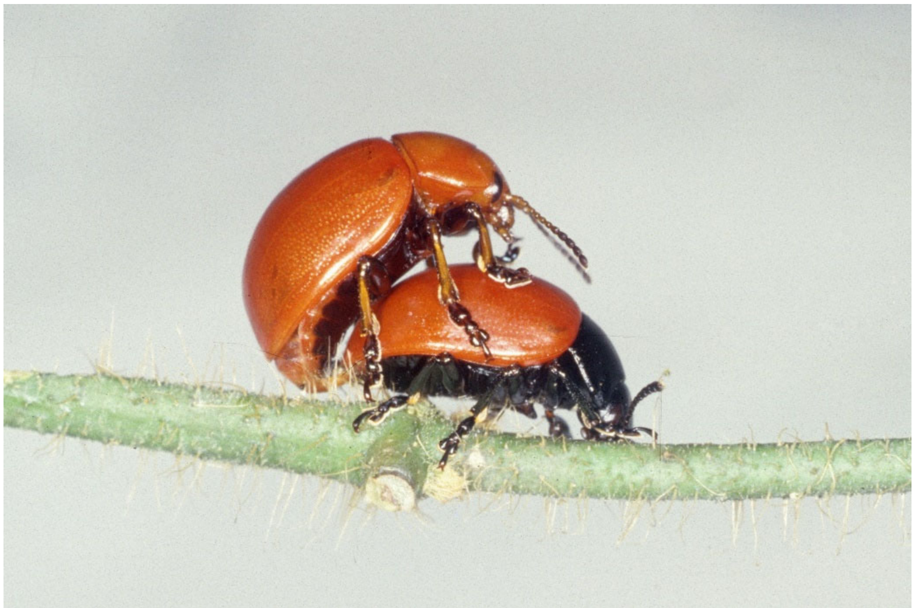

*Platyphora decens* (Stål, 1859), photo by DWM.

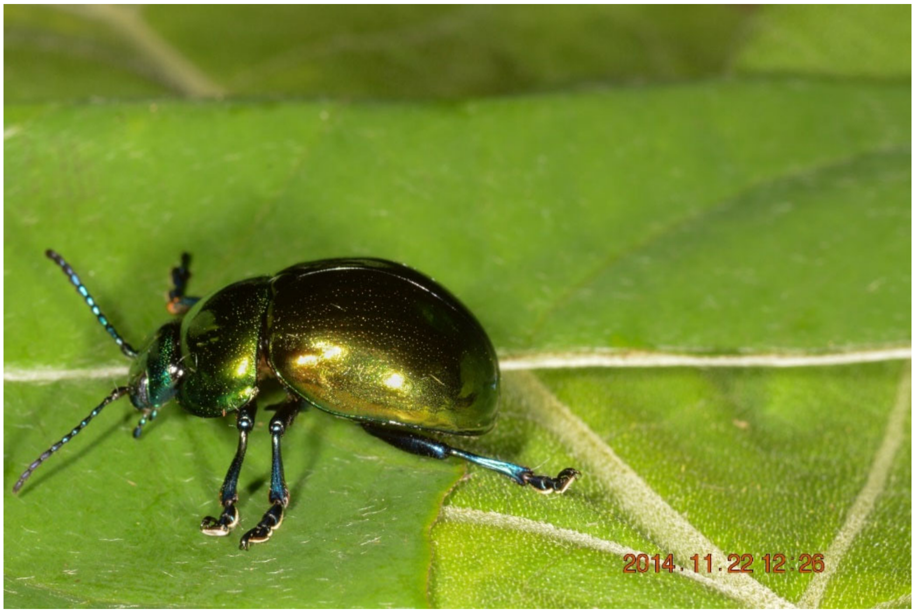


*Platyphora euchalca* (Stål, 1859), photo by DWM.

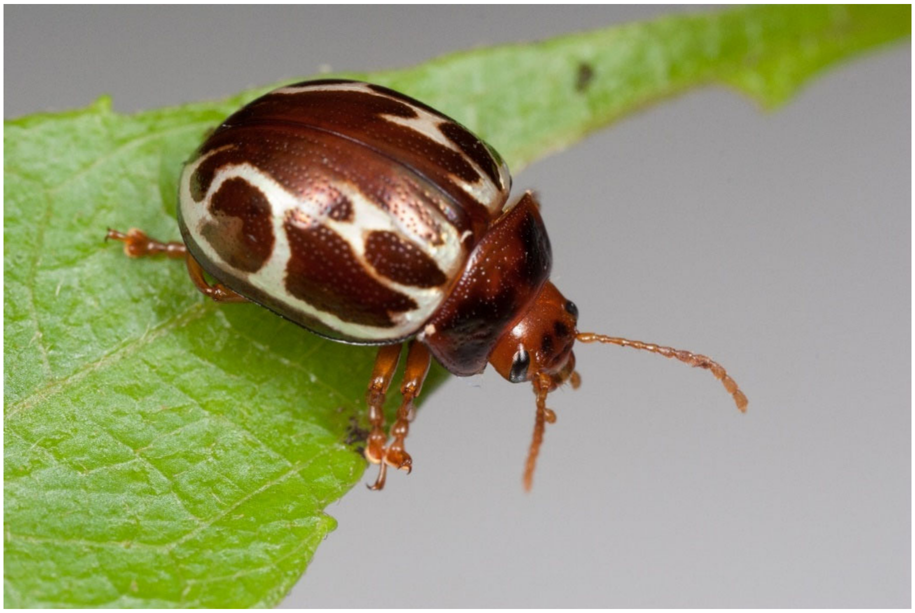

*Platyphora eucosma* (Stål, 1858), photo by GJD (under CC BY-NC 2.0).

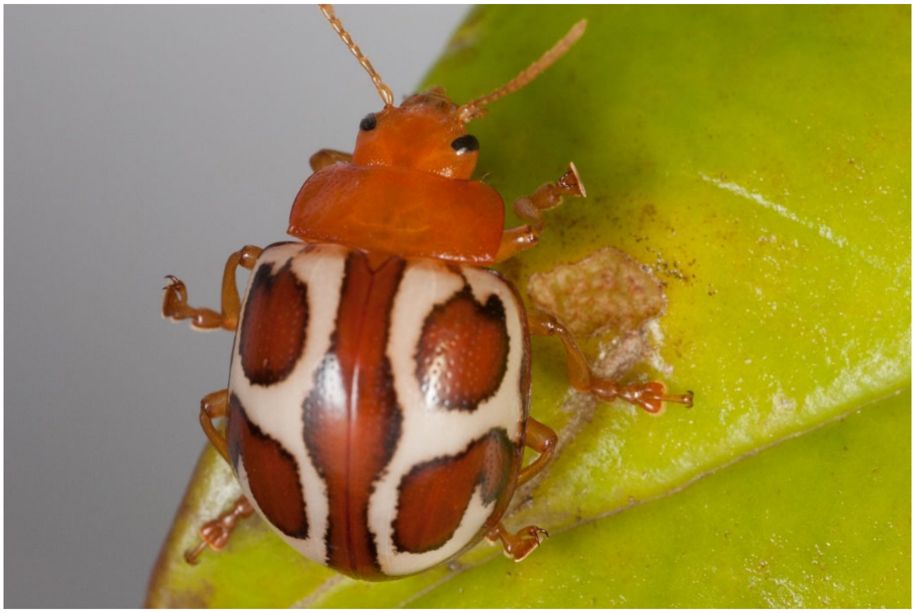

*Platyphora* sp. nov. “Fortuna”, photo by GJD (under CC BY-NC 2.0).

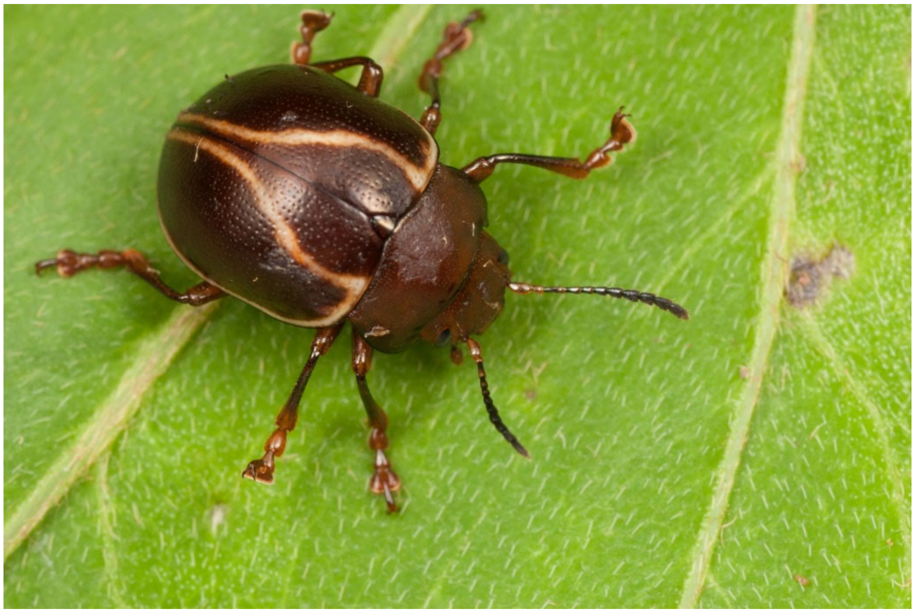

*Platyphora flexuosa* (Baly, 1858), photo by GJD (under CC BY-NC 2.0).

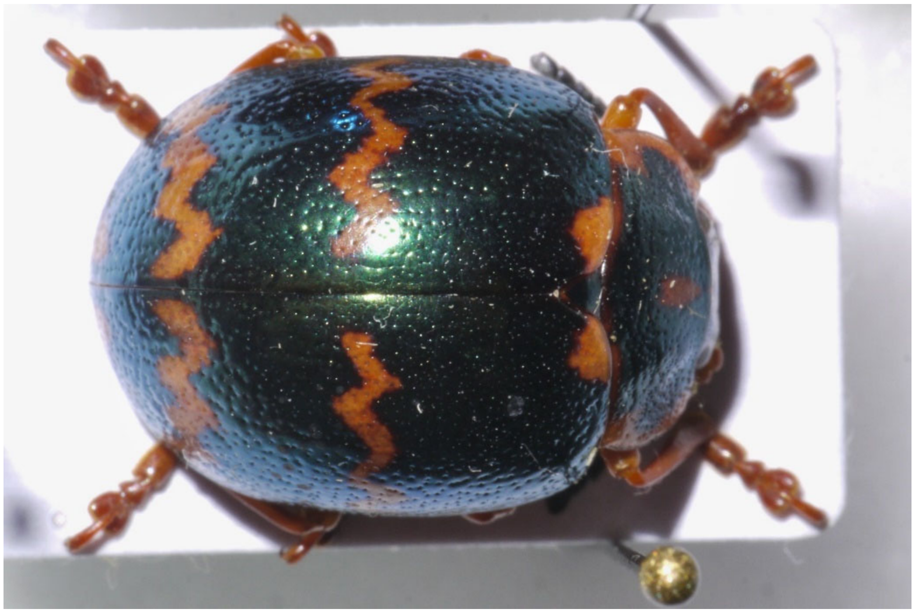

*Platyphora fulgora* (Stål, 1858), photo by DWM.

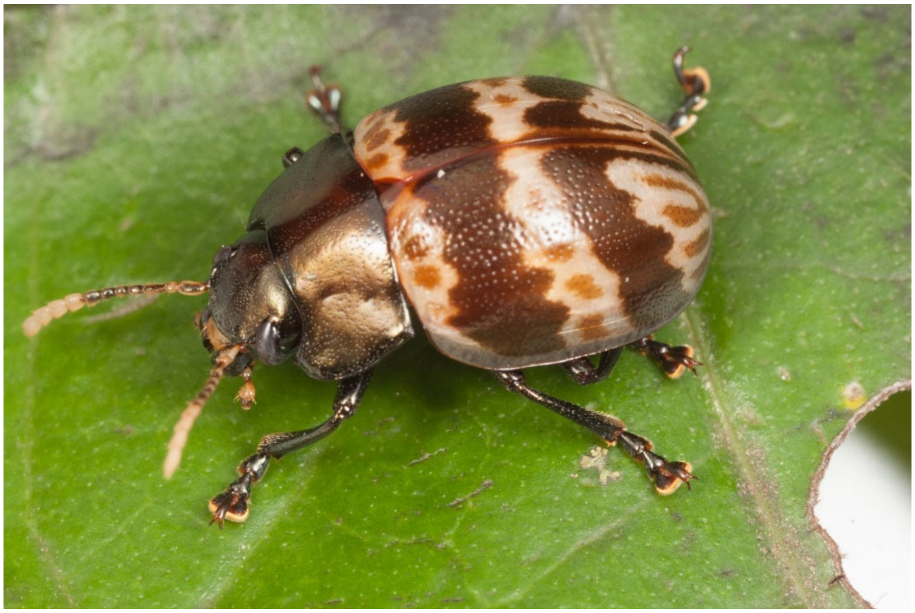

*Platyphora fulvicornis* (Guérin-Méneville, 1855), photo by GJD (under CC BY-NC 2.0).

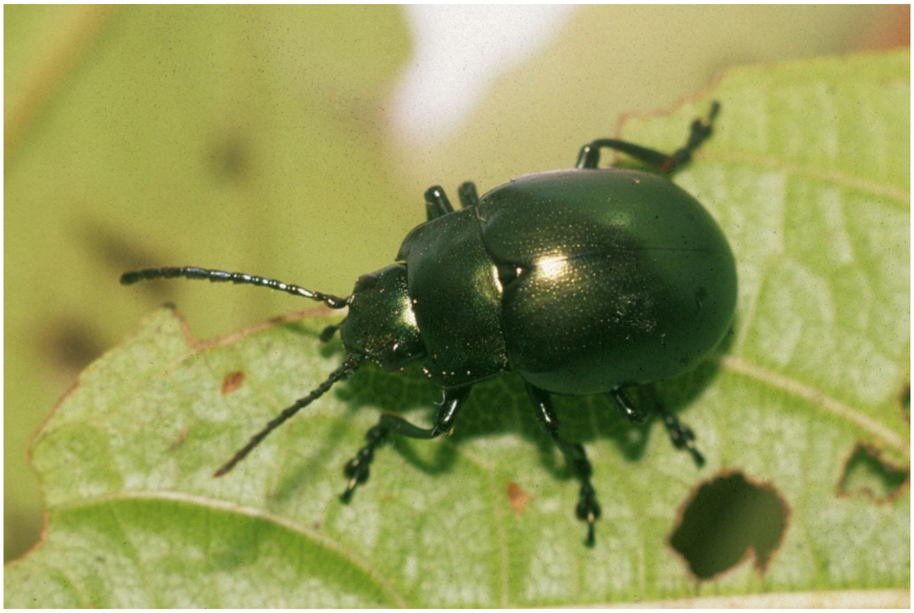

*Platyphora heliogenia* Bechyné, 1966, photo by DWM.

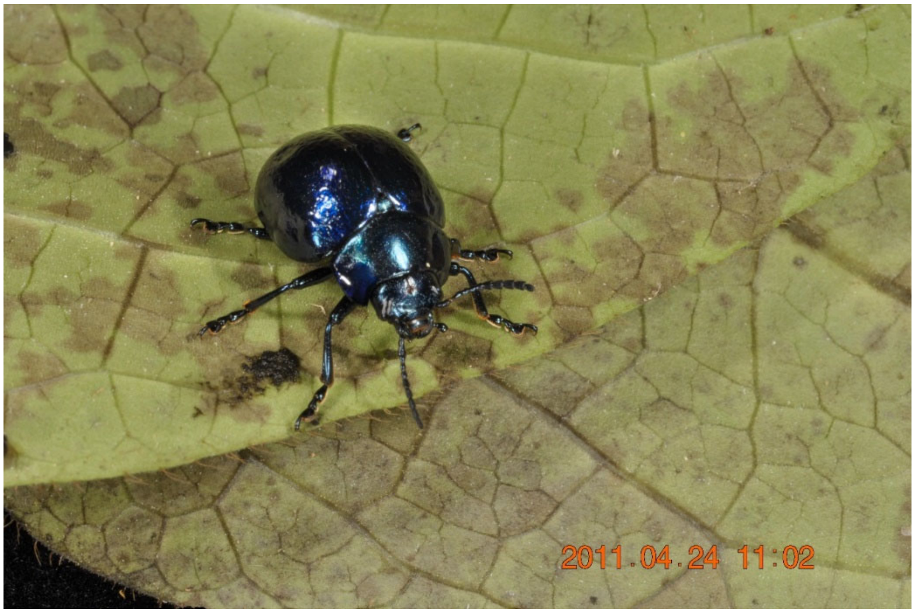


*Platyphora imitans* "blue" (Jacoby, 1903), photo by DWM.

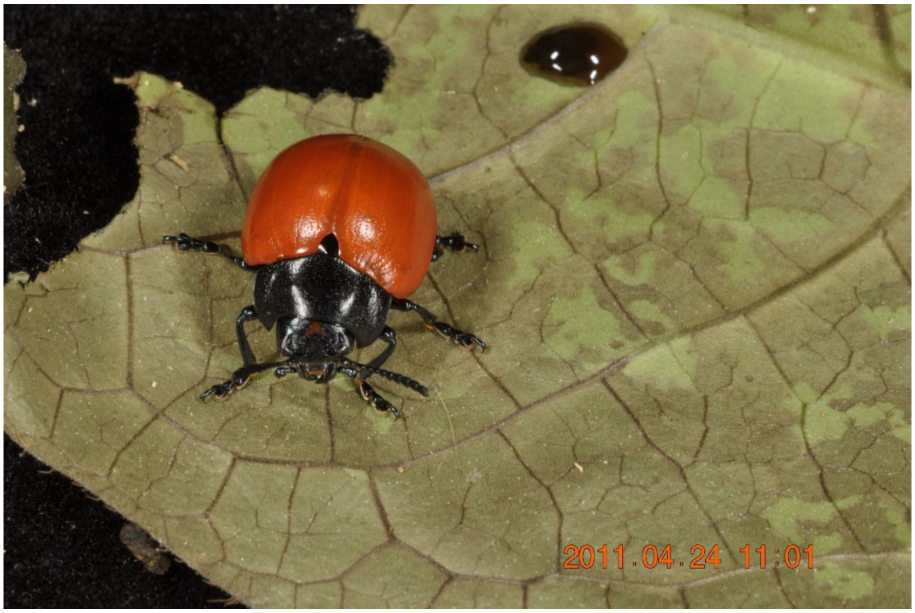


*Platyphora imitans* "red" (Jacoby, 1903), photo by DWM.

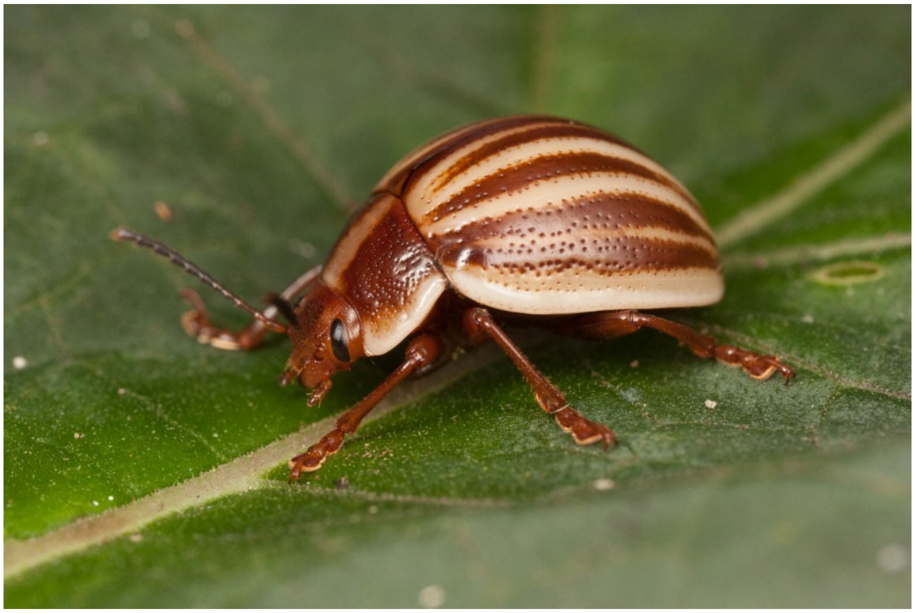

*Platyphora ligata adaequata* (Bechyné, 1954), photo by GJD (under CC BY-NC 2.0).

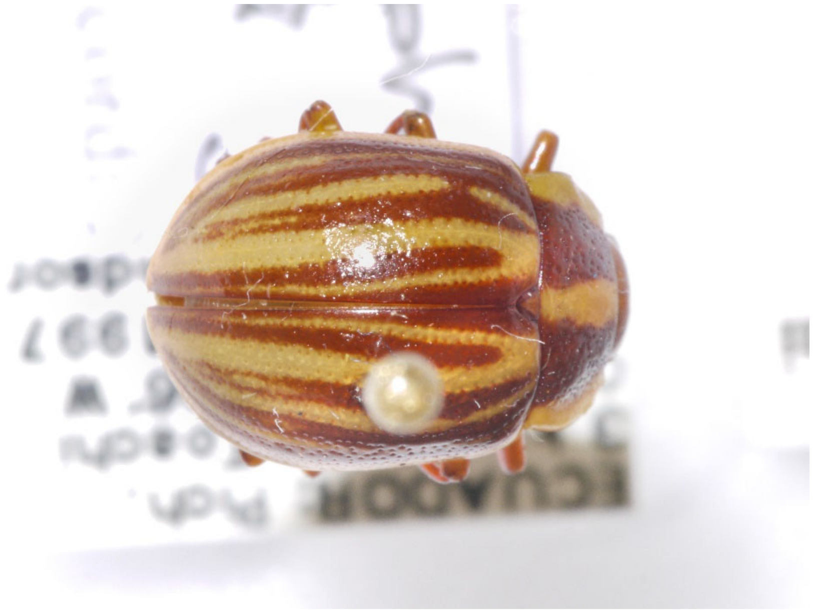

*Platyphora ligata ligata* (Stål, 1858), photo by DWM.

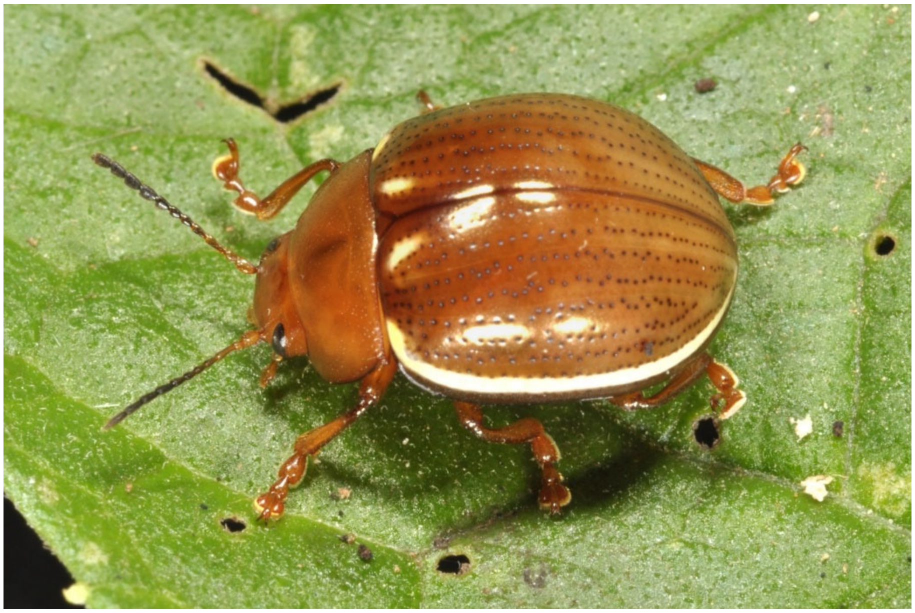

*Platyphora limbata* (Guérin-Méneville, 1844), photo by DWM.

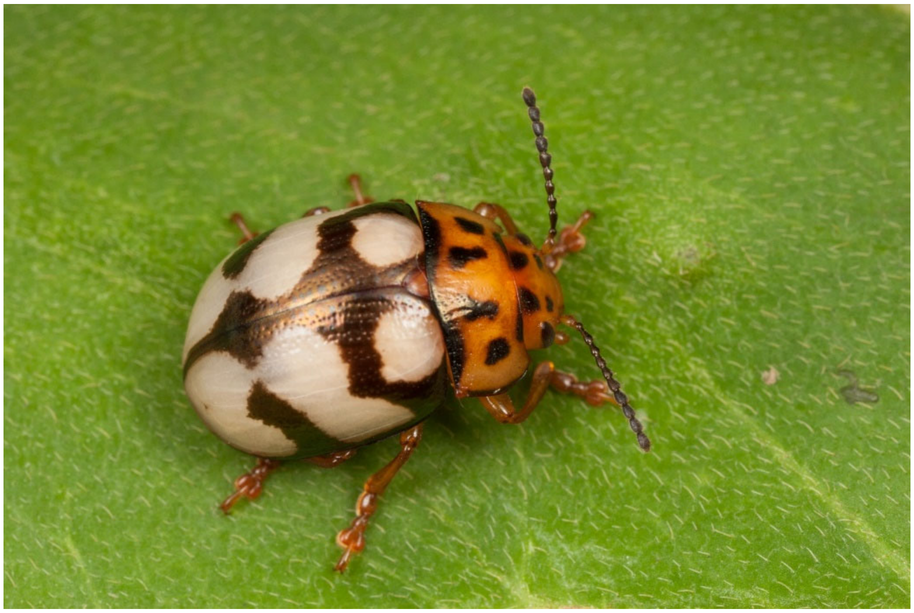

*Platyphora luteipennis* (Steinheil, 1877), photo by GJD (under CC BY-NC 2.0).

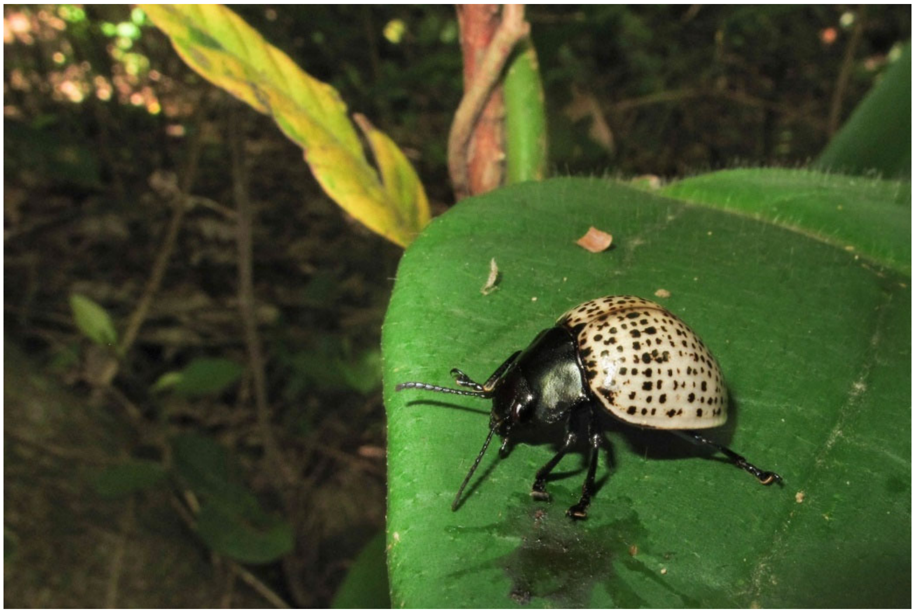

*Platyphora megistomelina* (Bechyné, 1954), photo by DWM.

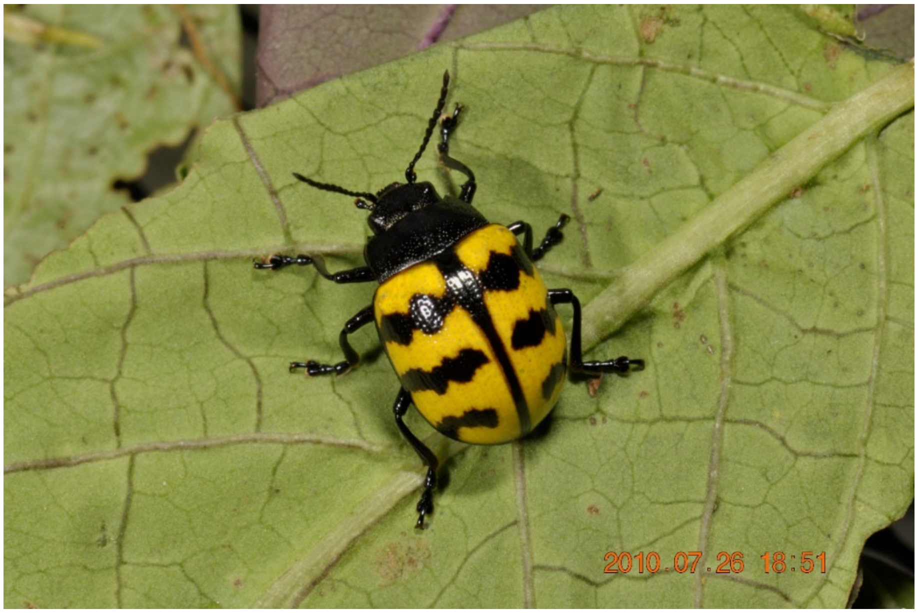


*Platyphora microspina* (Bechyné, 1954), photo by DWM.

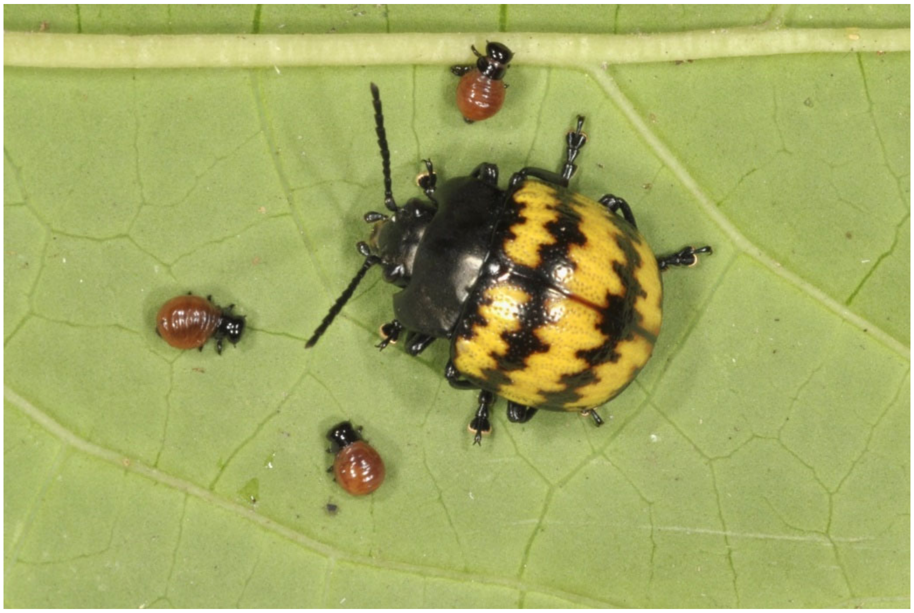

*Platyphora olala* (Bechyné, 1954), photo by DWM.

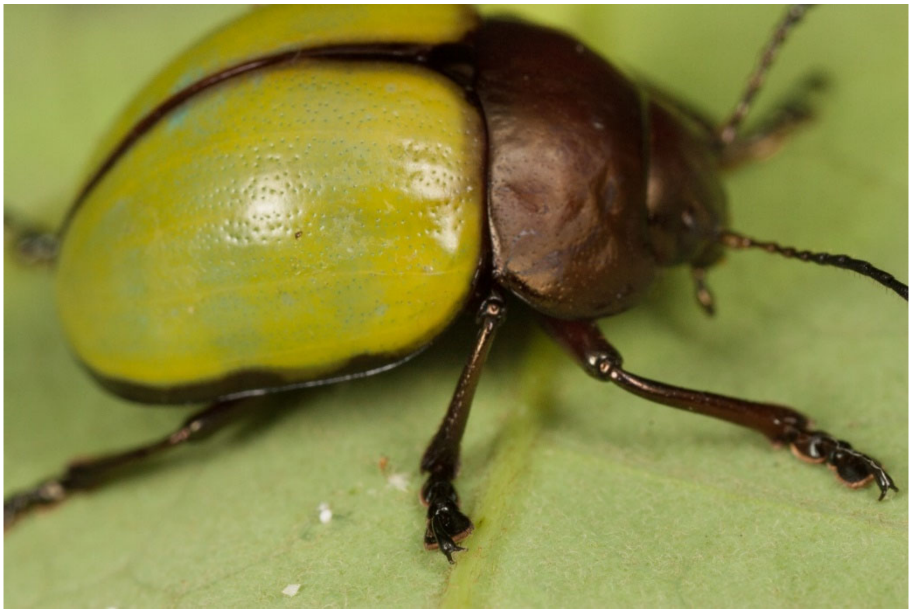

*Platyphora opima* (Stål, 1858), photo by GJD (under CC BY-NC 2.0).

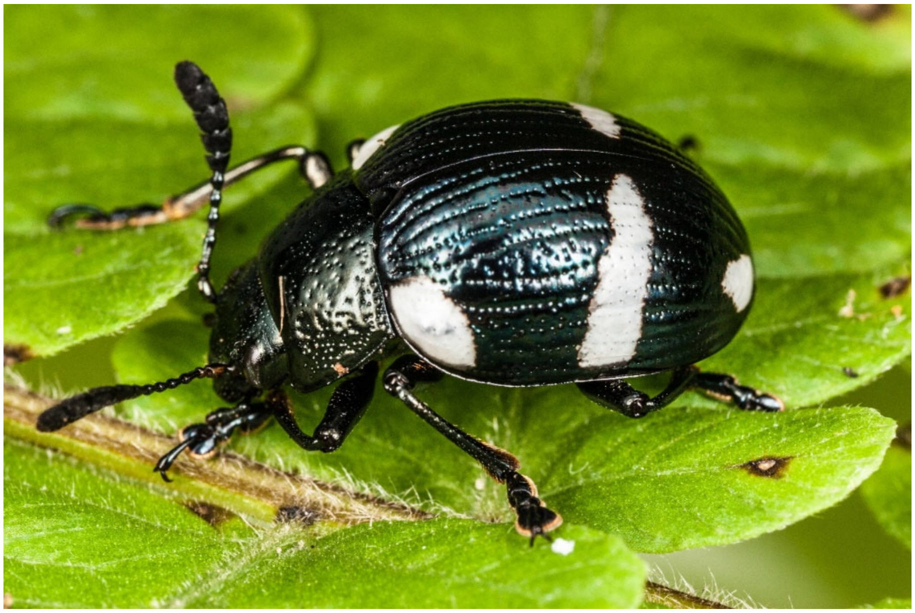

*Platyphora paradoxa* (Achard, 1914), photo by GJD (under CC BY-NC 2.0).

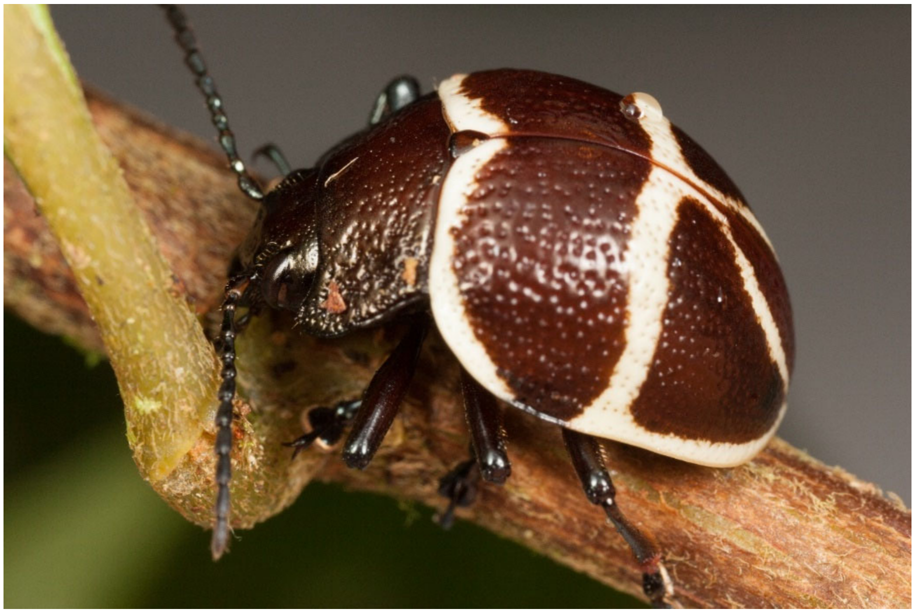

*Platyphora punctipennis* (Jacoby, 1878), photo by GJD (under CC BY-NC 2.0).

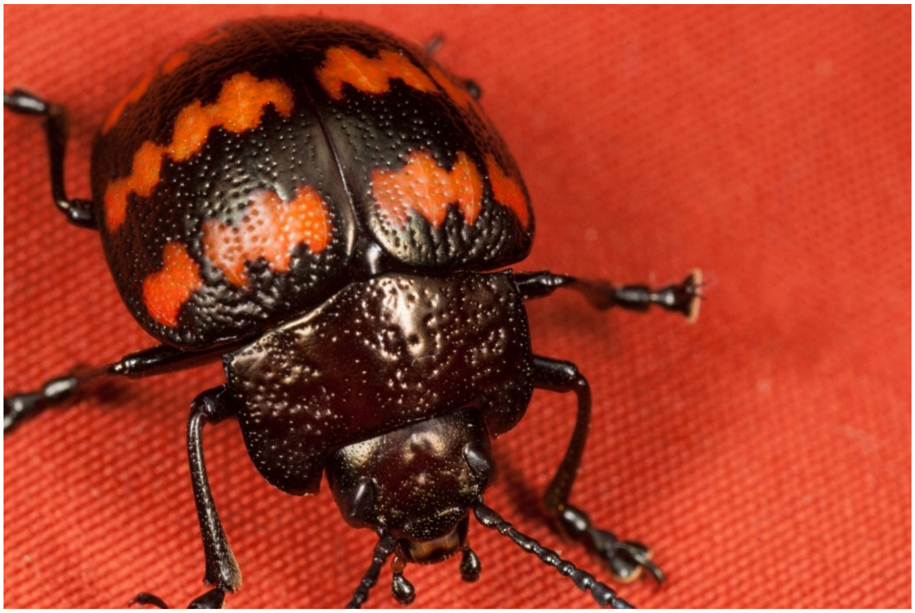

*Platyphora rubropunctata* (De Geer, 1775), photo by GJD (under CC BY-NC 2.0).

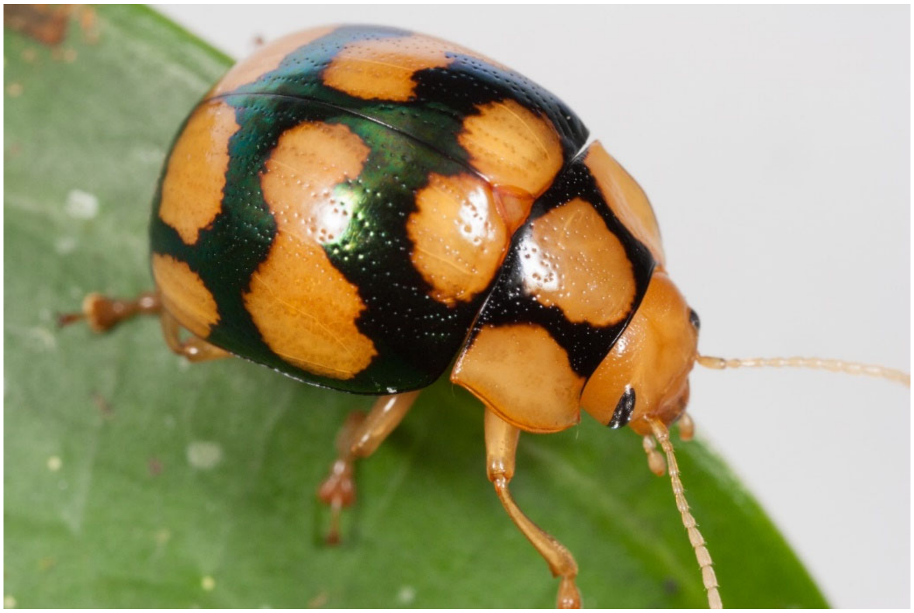

*Platyphora tangolita* (Bechyné, 1954), photo by GJD (under CC BY-NC 2.0).

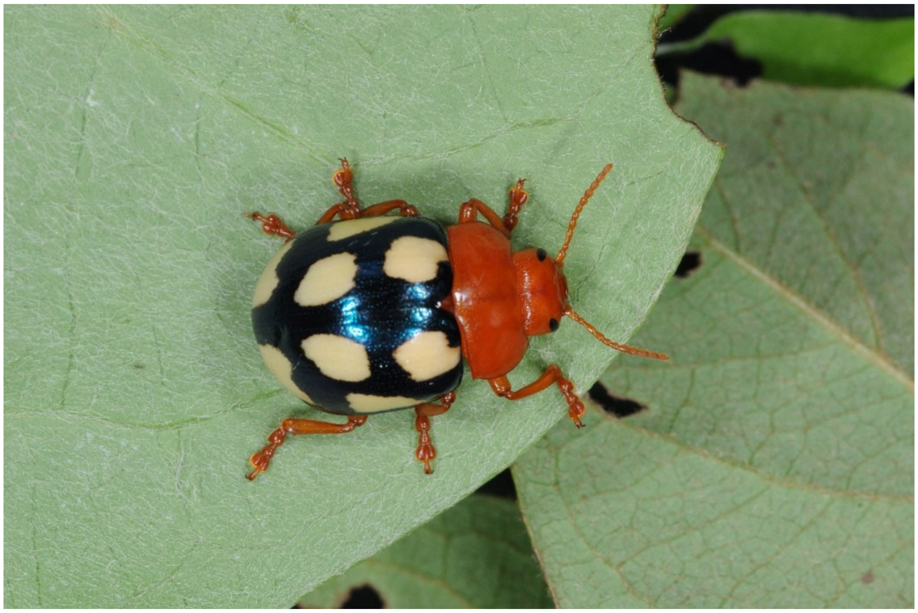

*Platyphora testudo* (De Mey, 1838), photo by DWM.

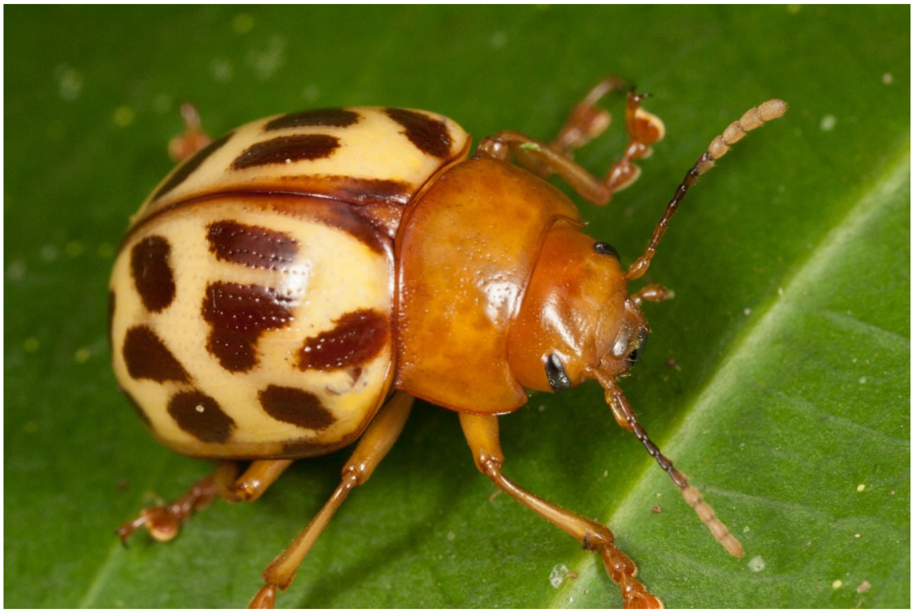

*Platyphora transversofasciata* Jacoby, 1883, photo by GJD (under CC BY-NC 2.0).

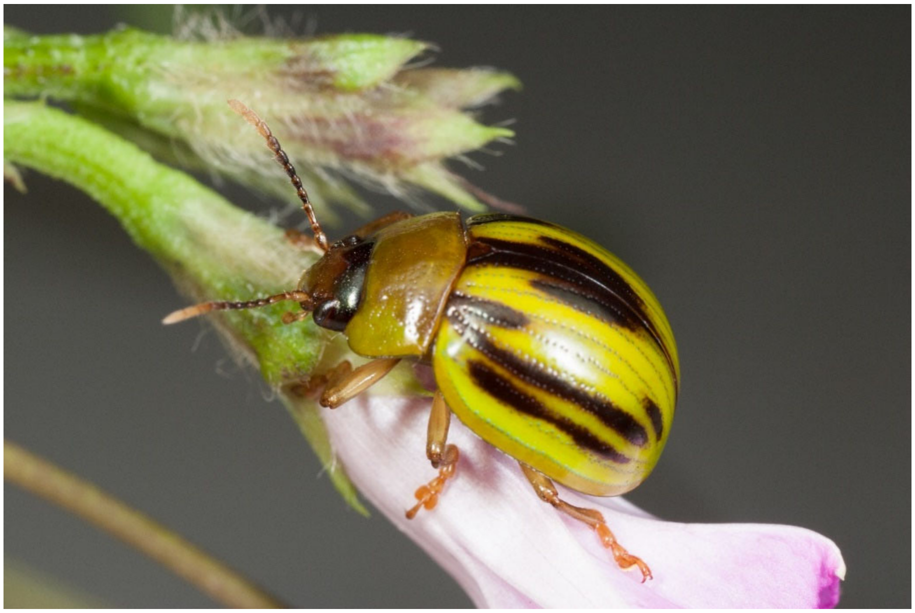

*Platyphora vespertina* (Baly, 1858), photo by GJD (under CC BY-NC 2.0).

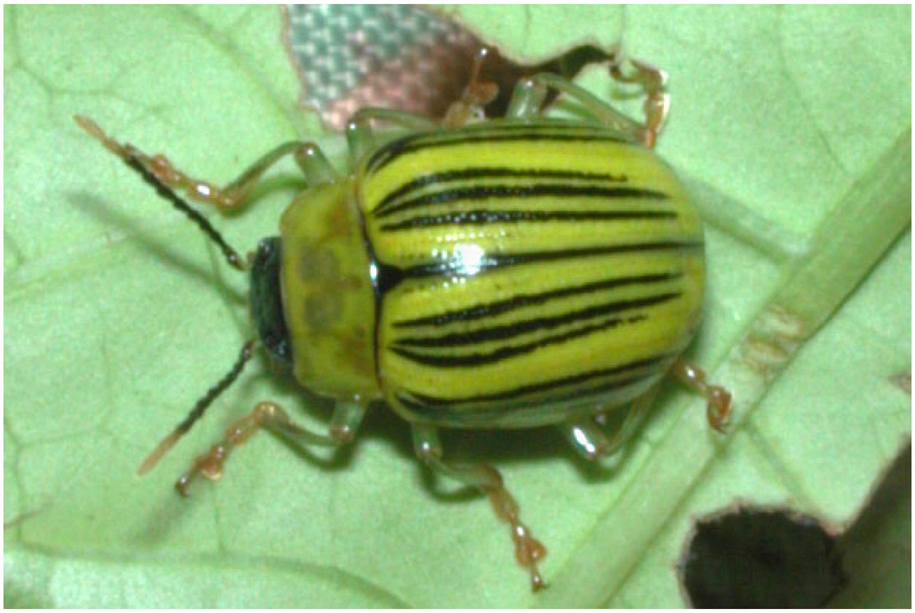

*Proseicela antennalis* (Kirsch, 1883), photo by DWM.

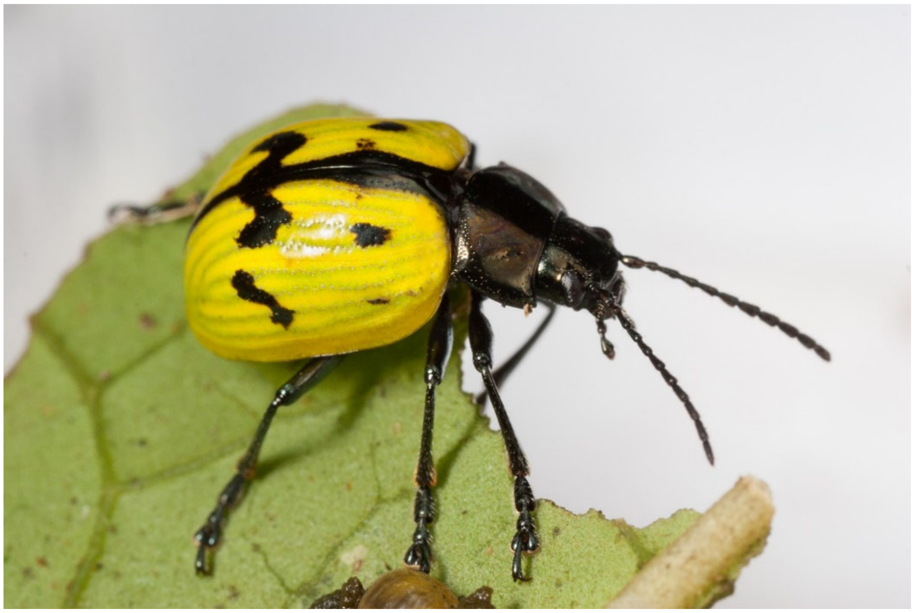

*Proseicela bicruciata* Jacoby, 1880, photo by GJD (under CC BY-NC 2.0).

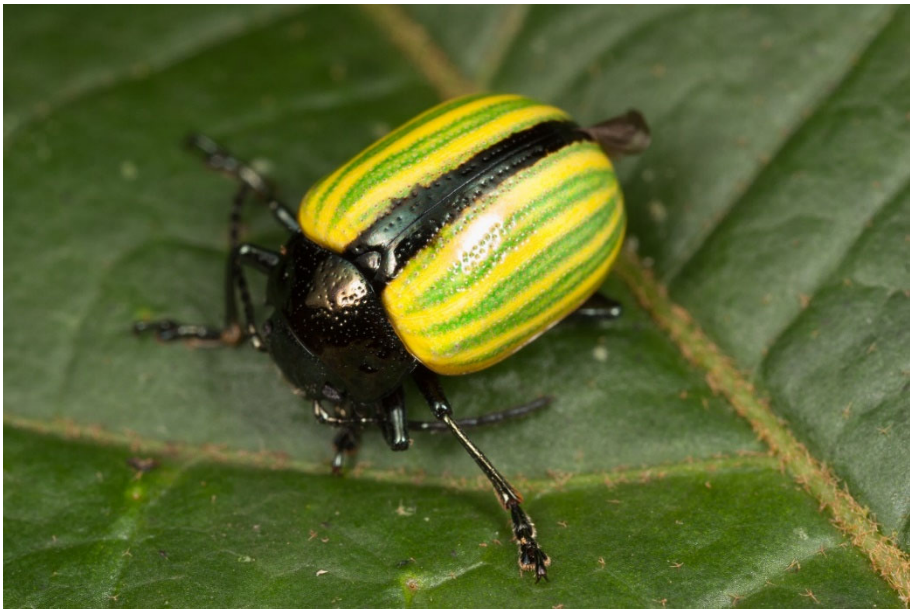

*Proseicela flavipennis* Erichson, 1847, photo by GJD (under CC BY-NC 2.0).

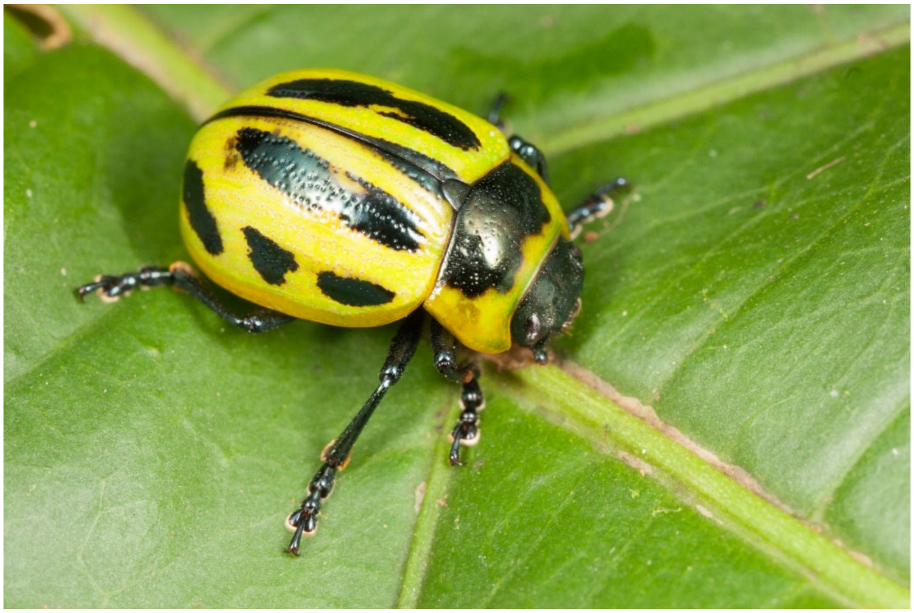

*Proseicela spectabilis* (Baly, 1858), photo by GJD (under CC BY-NC 2.0).

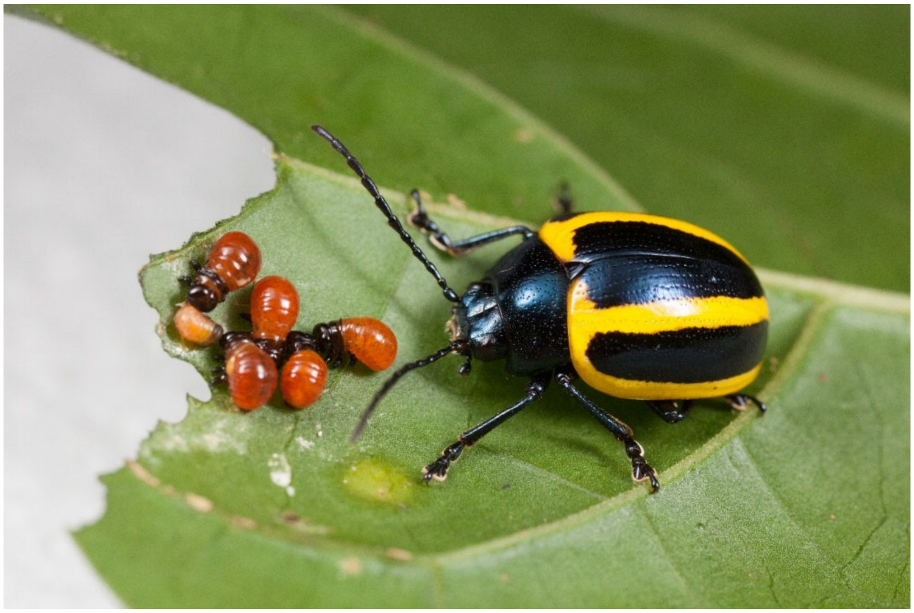

*Proseicela vittata* (Fabricius, 1781), photo by GJD (under CC BY-NC 2.0).

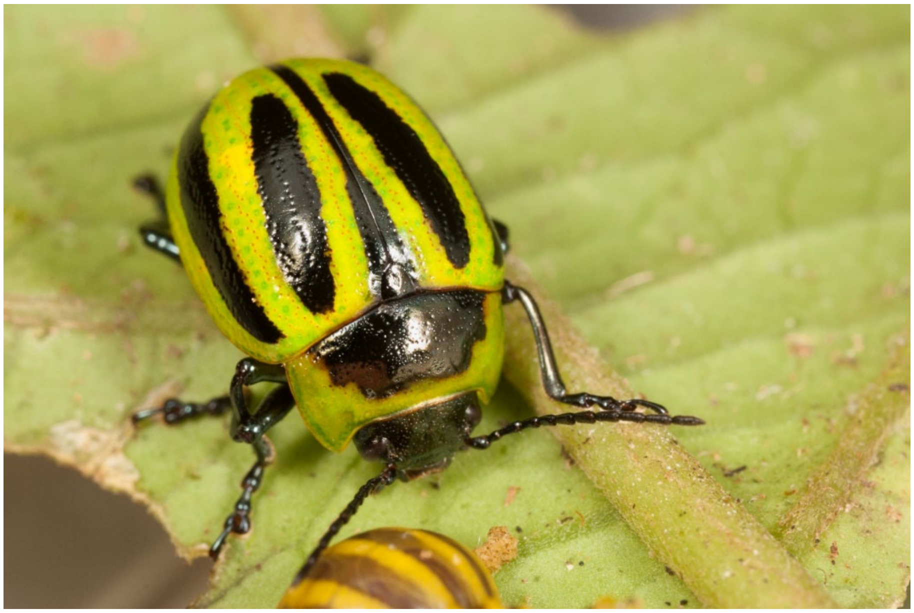

*Proseicela* sp. nov. "YSN", photo by GJD (under CC BY-NC 2.0).

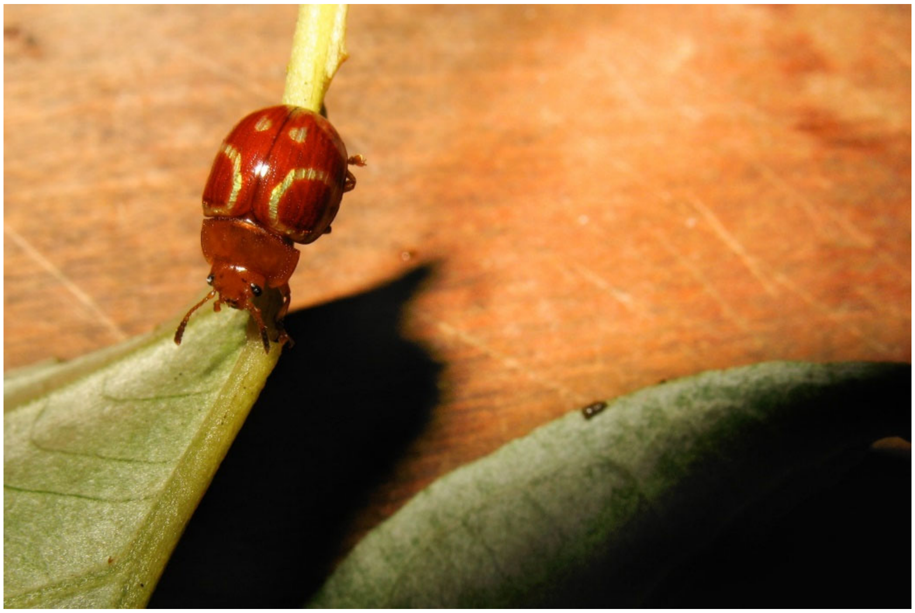

*Stilodes annuligera* (Erichson, 1847), photo by DWM.

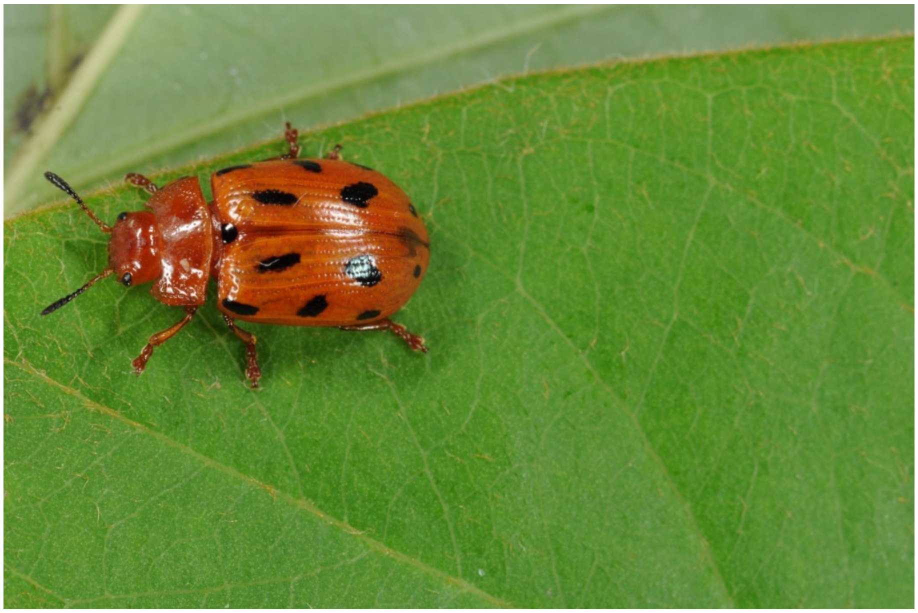

*Stilodes duodecimmaculata* (Stål, 1859), photo by DWM.

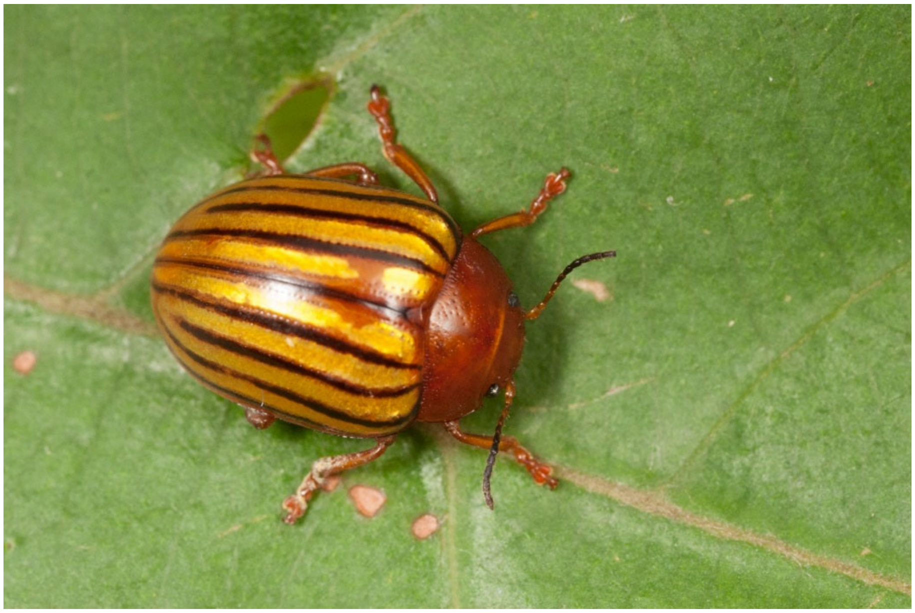

*Stilodes fuscolineata* (Stål, 1865), photo by GJD (under CC BY-NC 2.0).

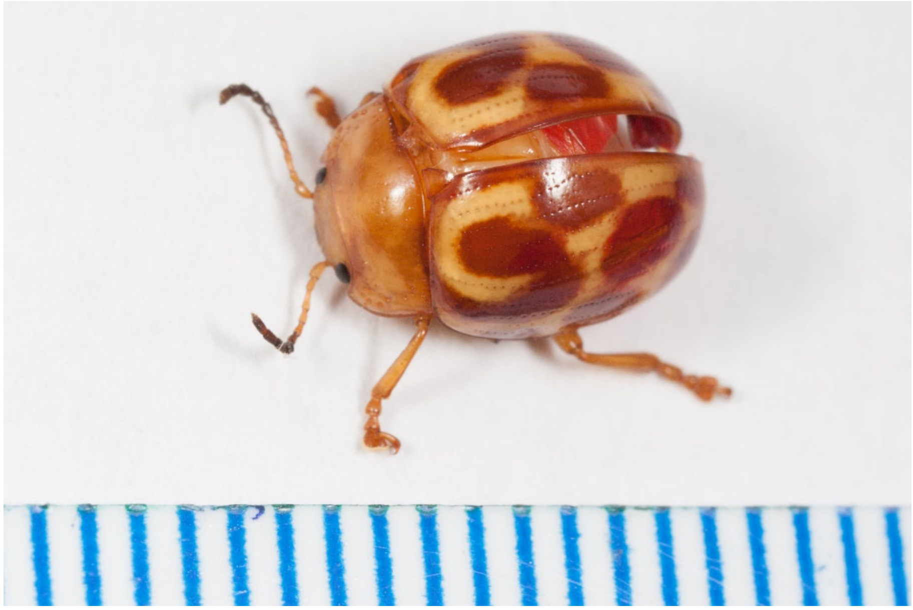

*Stilodes motschulskyi* (Stål, 1865), photo by GJD (under CC BY-NC 2.0).

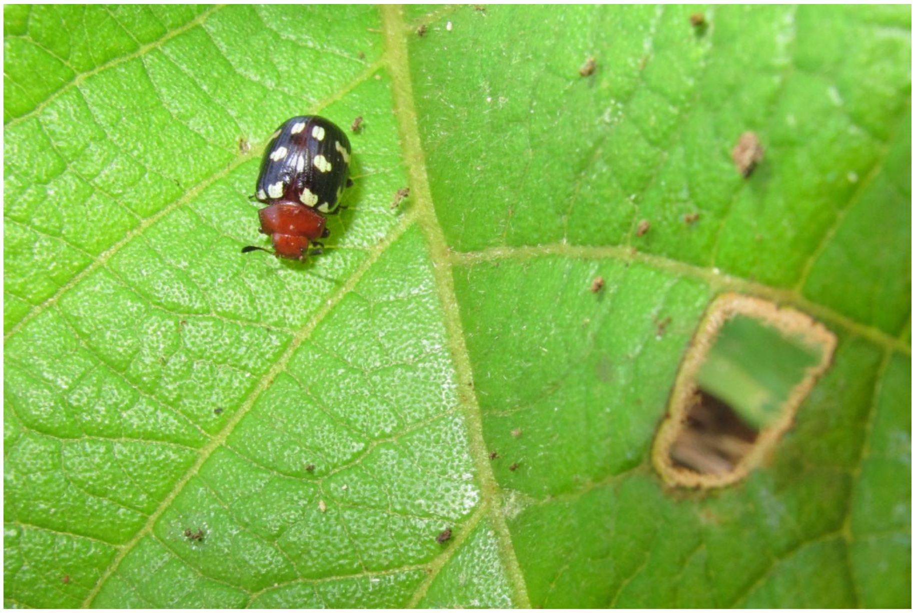

*Stilodes* (*Linographa*) *musicalis* (Stål, 1859), photo by DWM.

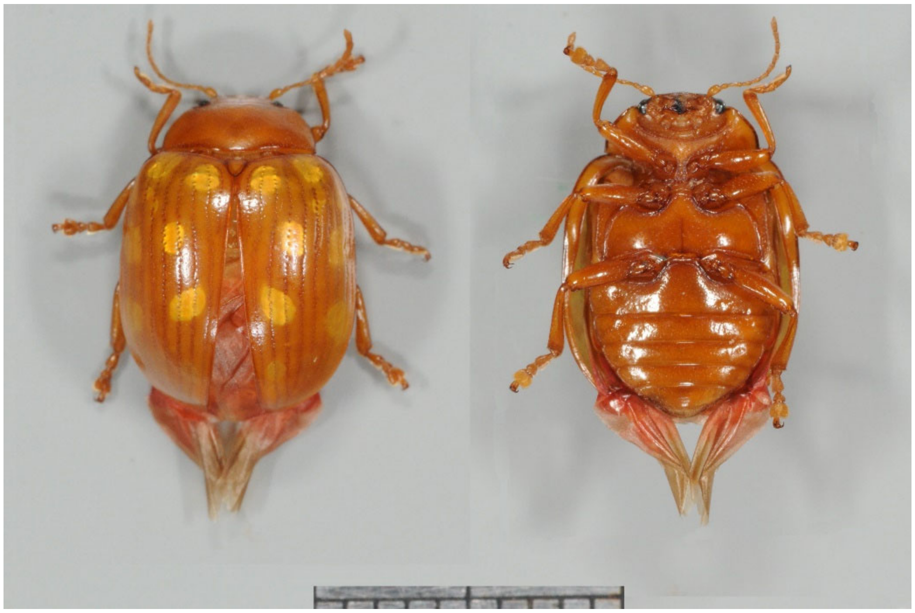

*Stilodes* cf. *pavonina* (Stål, 1860), photo by DWM.

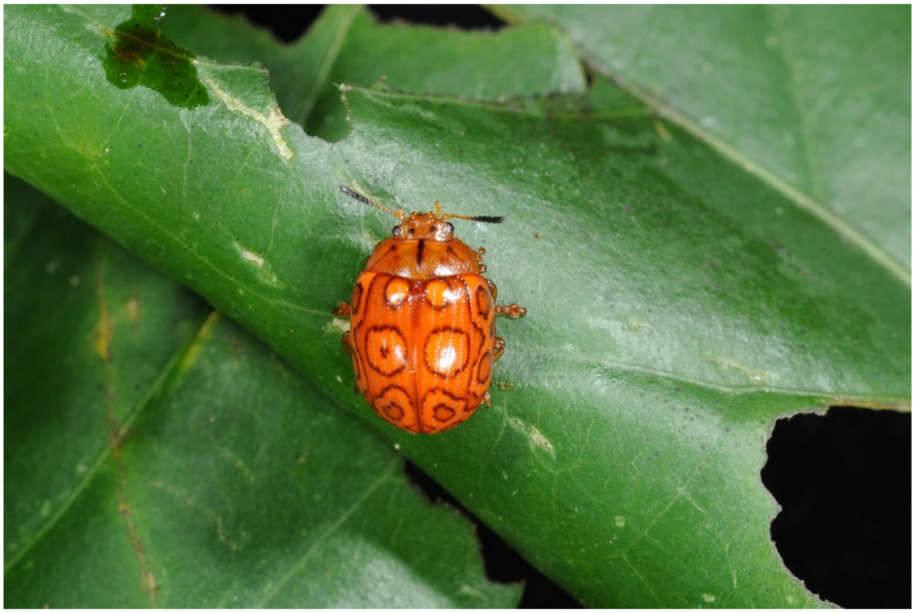

*Stilodes* sp. nov. “PDG 1”, photo by DWM.

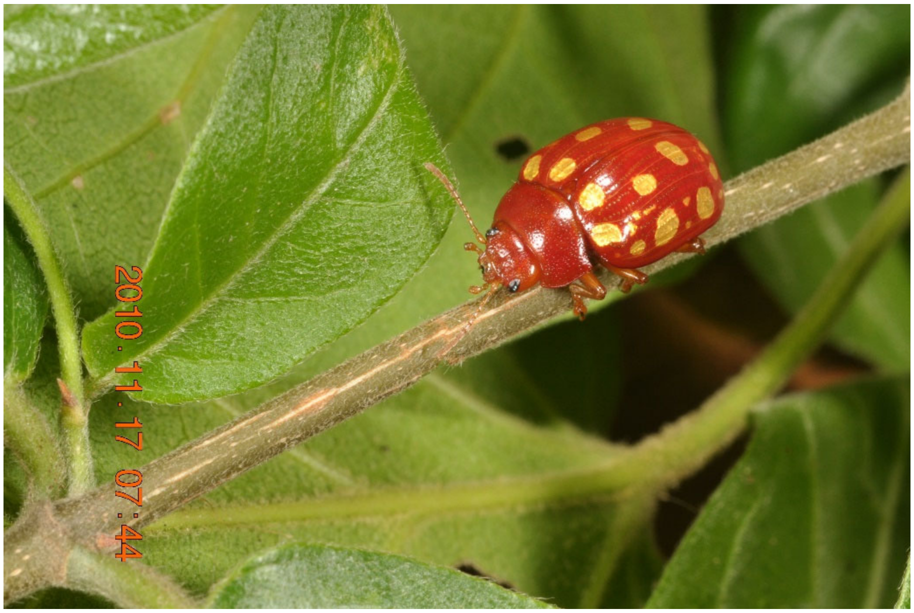


*Stilodes* sp. nov. “PDG 2”, photo by DWM.

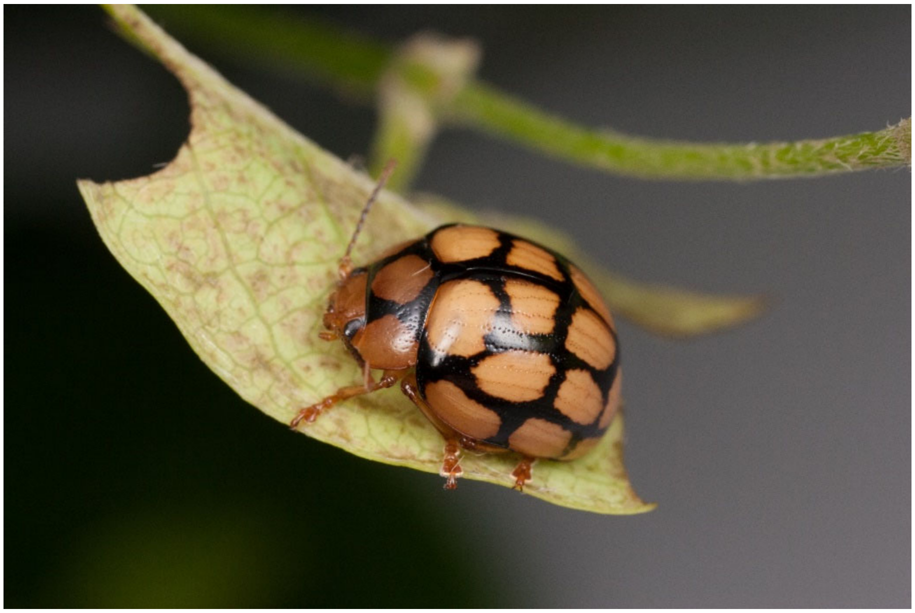

*Stilodes* nr. *retifera* Bechyné, 1954, photo by GJD (under CC BY-NC 2.0).

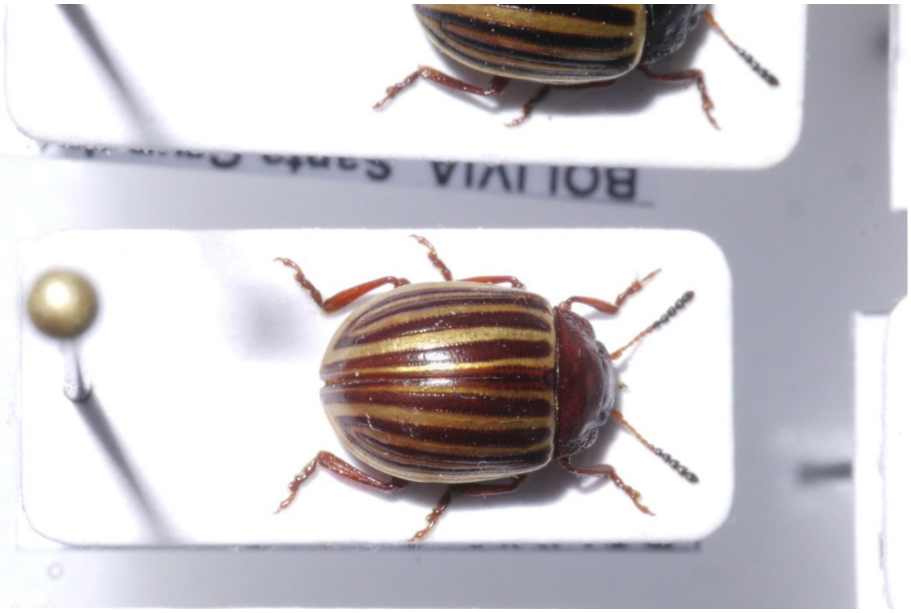


*Tritaenia alternata* (Kirsch, 1876), photo by LS.

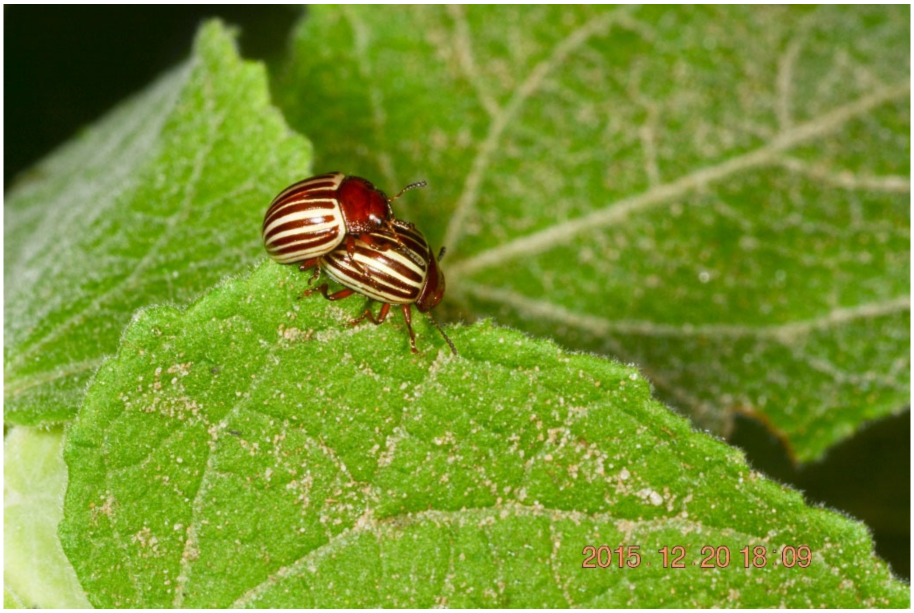


*Tritaenia alternata* (Kirsch, 1876), photo by DWM.

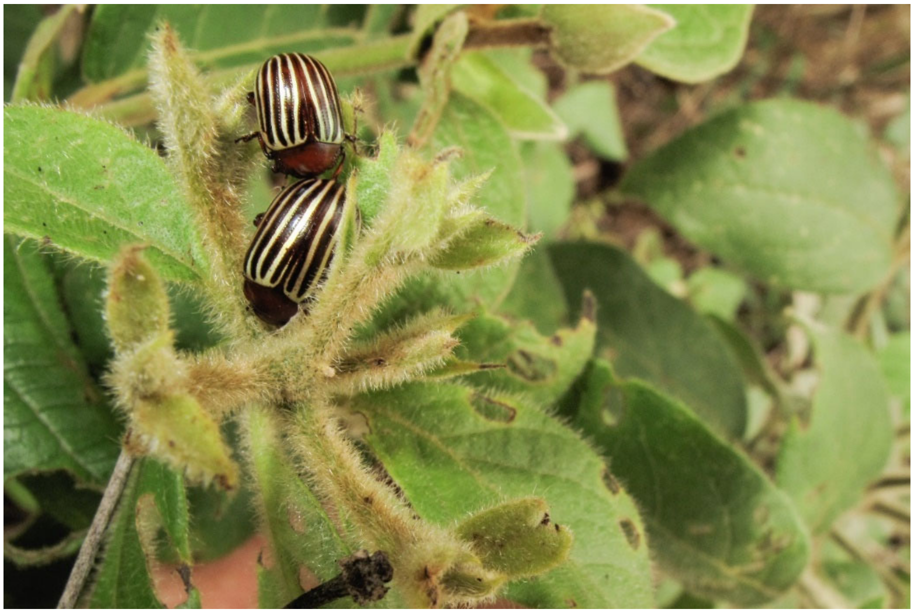

*Tritaenia sladenae* (Gahan, 1903), photo by DWM.
